# Real time quantification reveals novel dynamic processes in pancreatic lumenal network establishment and remodeling

**DOI:** 10.1101/2023.08.18.552936

**Authors:** Abigail Laura Jackson, Silja Heilmann, Christine Ebeid, Jelena Miskovic Krivokapic, Jose Alejandro Romero Herrera, Henrik Semb, Pia Nyeng

**Author notes:** co-first authors (Abigail Laura Jackson, Silja Heilmann).

## Abstract

During embryogenesis dynamic changes in tissue architecture transform primitive anlages to functional organs. Here we document in real time how pancreatic lumens are derived and transformed using a new apical-polarity mouse reporter. Our 4D imaging data reveals dynamic remodeling of apical proteins and lumens to primarily drive each stage of pancreatic duct development. Furthermore, we pinpoint two unique transitions during lumenogenesis. Contrary to current “*de novo*” models of polarity acquisition, we show that expansion and rearrangement of the pre-existing central primary lumen drives early network growth. We also document how the endocrine promoting niche - a “plexus” of interconnected ducts - is resolved. We show that an arborized network forms by gradual closing of ductal loops, rather than via pruning. These novel tissue dynamics provide a new framework within which cell and molecular signaling can be investigated to better understand the interplay between organ architecture and cell fate.

## Introduction

Tubes are fundamental building blocks of an organism and essential for the function of many organs that transport vital gasses and enzymes around the body. Understanding how tubes form in development can help us better understand the process that goes awry in congenital tube defects (Dimitriou et al. 2018; Bergmann et al. 2018; Cnossen and Drenth 2014).

Tubulogenesis has been studied in 3D models based on cell lines such as the Madin-Darby canine kidney (MDCK) cells (Martín-Belmonte et al. 2008) and during organogenesis of the lung, salivary gland, pancreas etc (Lubarsky and Krasnow 2003). Based on these studies, tube formation is traditionally divided into two main categories: A) tubes that emerge from an already polarized epithelium, and B) tubes that emerge independently from a non-polarised epithelium (Hogan and Kolodziej 2002). In (A) the center of a tube - the lumen - is created through processes such as budding (e.g. the lung) and wrapping (e.g. the neural tube). In (B) lumens form *de novo* either by cavitation, where central cells undergo apoptosis (e.g. in mammary gland end buds (Debnath et al. 2002), by hollowing, where cords of lumens form within cells, or by cells acquiring “de novo” apical polarity (e.g. in 3D cell culture of MDCK cells, (Martín-Belmonte et al. 2008; Mangan et al. 2016))

The pancreas is a complex endocrine and exocrine organ containing a branched network of tubes. To achieve a postnatal exocrine function the pancreas must form a tree-like ductal structure, ensuring that acinar derived enzymes, fluid and bicarbonate can flow from the terminal end buds, through the ductal network, and into the duodenum (Reichert and Rustgi 2011). The formation of this network and its subsequent remodeling is not fully understood. Congenital anomalies in pancreatic ductal morphology have been associated with symptoms such as recurrent abdominal pain, nausea and vomiting (Türkvatan et al. 2013). The inefficient drainage resulting from developmental defects in pancreatic ductal remodeling can ultimately cause inflammation and pancreatitis. In addition to this structural role, the remodeling pancreatic ducts provide a unique niche necessary for specification of endocrine cells during embryonic development (Löf-Öhlin et al. 2017; Kesavan et al. 2009; Bankaitis, Bechard, and Wright 2015; Azizoglu et al. 2017; Barlow et al. 2023). The duct begins as an epithelial evagination of the foregut endoderm at the center of which lies a central primary lumen (CPL). The CPL is thought to be formed from an outbudding of the foregut lumen. Unlike MDCK cells and other simpler branching organs, the pancreatic duct does not remain a simple, unilumenal structure. The secondary pancreatic multi-lumenal network is believed to be created ‘de novo’ (Villasenor et al. 2010; Kesavan et al. 2009; Hick et al. 2009; Flasse, Schewin, and Grapin-Botton 2020). These studies propose that the process has four phases, beginning with 1) ‘de novo’ acquisition of apical polarity in a few scattered cells, followed by 2) formation of a microlumen between neighboring polarized cells and 3) merging of microlumens to form a connected network that eventually 4) connects to the central gut lumen. In our study we set out to test this model using live imaging.

Studies in the mouse show that from E14.5, the ductal system found at the center of the pancreas consists of a network of interconnected ducts, coined the ductal “plexus” (Bankaitis, Bechard, and Wright 2015; Dahl-Jensen et al. 2018; Villasenor et al. 2010; Kesavan et al. 2009). At the periphery of and connected to this structure, are a system of ramifying branches. The plexus later resolves into an arborized structure by an unknown mechanism (Dahl-Jensen et al. 2018). The branches at the periphery of the pancreas have been demonstrated to form by lateral branching (Puri and Hebrok 2007), which has been proposed to be driven by proliferating progenitor cells at the ductal termini (Sznurkowska et al. 2018). Some epithelial branches contain several lumens at E11.75 (Villasenor et al. 2010), suggesting a disparity between the inner lumen topology and the outer branching of the epithelium. This observation is reinforced by the finding that outer branching morphogenesis, but not inner lumen formation, is perturbed by inhibiting extracellular matrix (ECM)-cell interactions (Shih et al. 2016). Thus, a combination of lumen remodeling and outer peripheral branching seems to underlie pancreatic ductal network morphogenesis. However, it remains unclear how these processes are coordinated to resolve the epithelium into an arborised structure consisting of single lumenal branches.

These unique phases of pancreatic tubulogenesis suggest that each transition is a highly dynamic process. Yet, with the exception of a few studies by ourselves and others (Azizoglu et al. 2017; Shih et al. 2016; Löf-Öhlin et al. 2017; Darrigrand et al. 2024), the majority of studies on murine pancreatic tubulogenesis have relied on static histochemical analyses. Similarly, most “real time” murine branching studies in other organs have been performed in either low temporal frequency or in 2D (Riccio et al. 2016; Mederacke et al. 2022; Metzger et al. 2008)(Araújo and Llimargas 2023; Mullapudi et al. 2019; Scheele et al. 2017; Packard et al. 2013)(Riccio et al. 2016; Mederacke et al. 2022; Metzger et al. 2008)Meanwhile, high frequency 4D (x, y, z, time) imaging of non-mammalian models of branching organs have also contributed to the advancement of the field (Araújo and Llimargas 2023; Mullapudi et al. 2019; Scheele et al. 2017; Packard et al. 2013). Here we bridge a gap and advance our understanding of mammalian tubulogenesis, by quantifying the dynamics of lumenal network development through analysis of high temporal frequency 4D data.

We introduce a new mouse strain for live imaging of apical proteins - the Muc1-mCherry fusion-protein reporter, which we believe will be of great benefit to cell, developmental biology and cancer research communities. Mucin 1 (MUC1) is a large transmembrane protein which is expressed at apical membranes and developing lumens of many branched organs (Lacunza et al. 2010; Sakurai et al. 2007; Chambers et al. 1994; Fanni et al. 2012), including the pancreas (Cano et al. 2004; Kesavan et al. 2009; Villasenor et al. 2010; Kopinke and Murtaugh 2010), but becomes mislocalized in cancer (Bafna, Kaur, and Batra 2010; Luan et al. 2022). MUC1 is heavily glycosylated on the extracellular N-terminal domain, which protrudes 200 nm above the cell surface and reduces cell-cell and cell-cell-extracellular matrix (ECM) adhesion (Wesseling, van der Valk, and Hilkens 1996). The MUC1 cytoplasmic domain interacts with multiple signaling pathways (Bafna, Kaur, and Batra 2010).

In this study we use Muc1-mcherry and other reporters to uncover two unique mechanisms driving pancreatic lumenogenesis. We document that the secondary pancreatic lumenal network is primarily generated from a highly dynamic transformation of the pre-existing CPL, rather than *de novo*. The resulting network of interconnected ducts contain multiple loop structures, which provide unique pancreatic progenitor niches and are resolved into an arborized structure by a novel mechanism, which we term “loop-closing”. By providing a new dynamic framework for lumen network formation and transition we hope future studies will be well placed to decipher how organ architecture and cell fate are interconnected at a cell and molecular level.

**Table 1:**
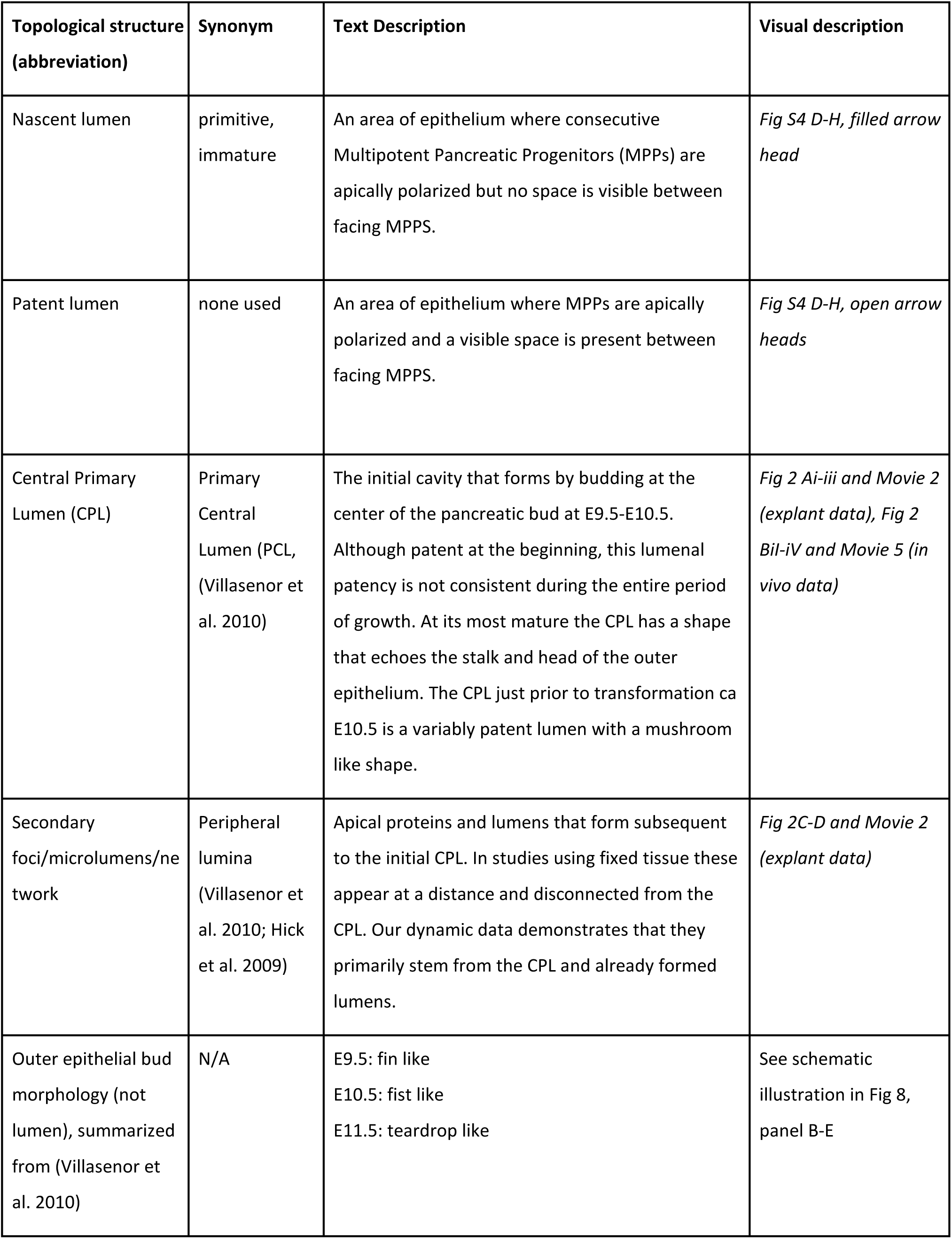
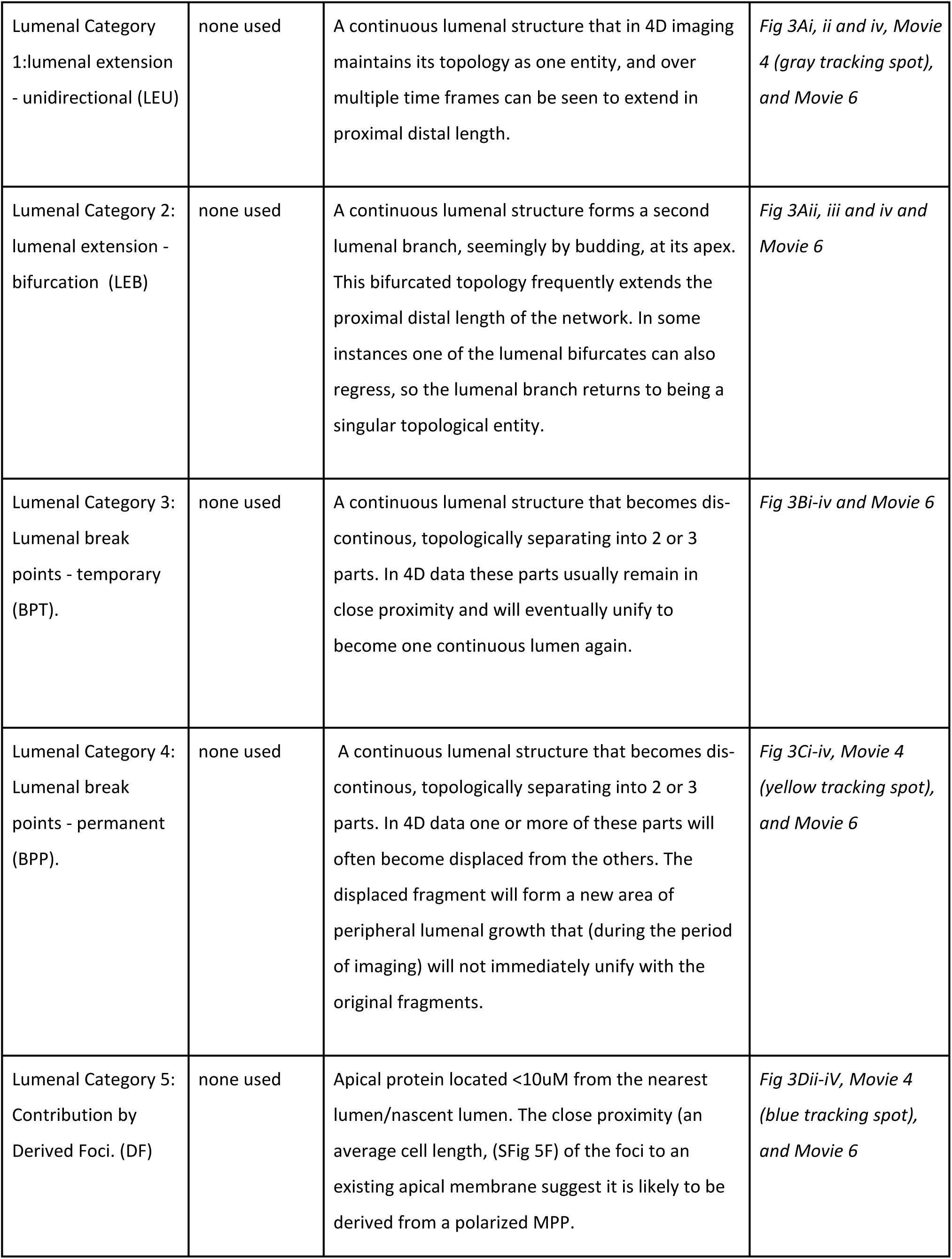
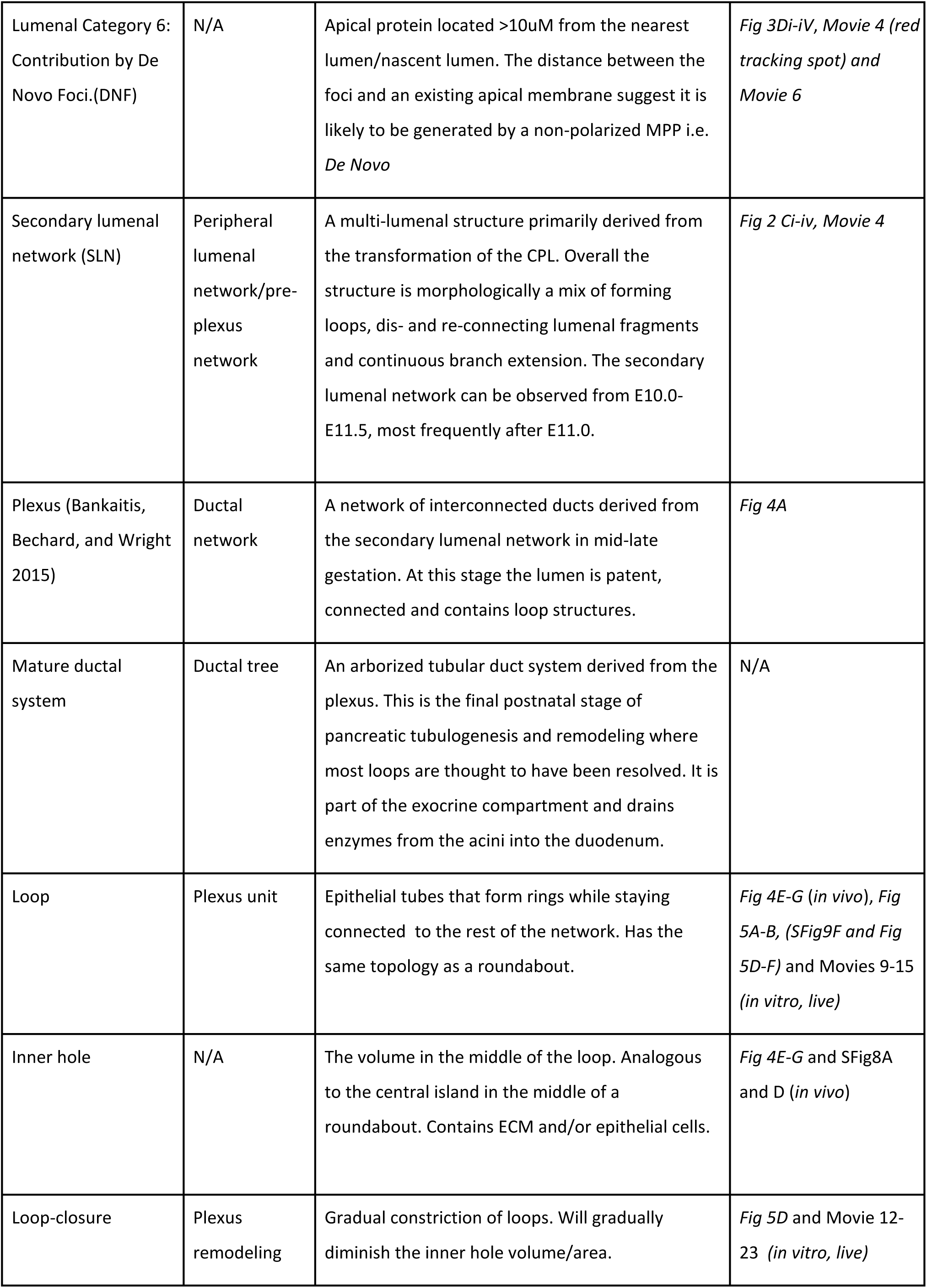
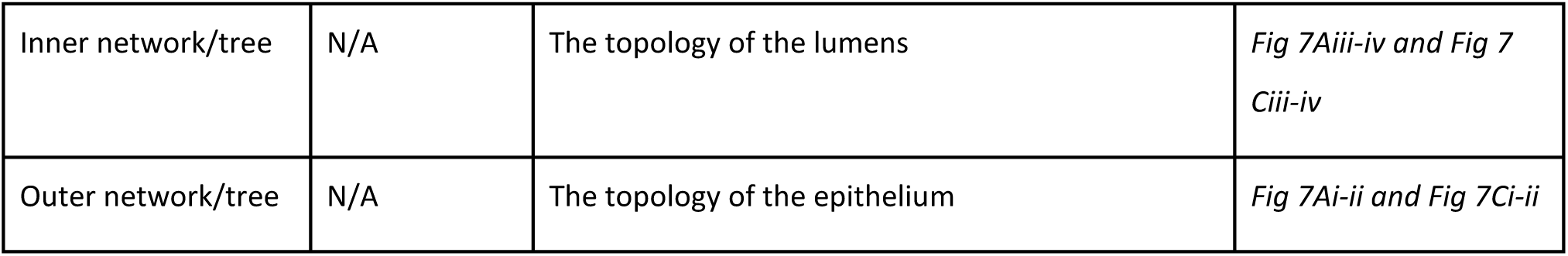
Definition of descriptive terminology.

## Results

### Generation of a Muc1-mCherry apical reporter mouse

To visualize apical polarity, we created a BAC (bacterial artificial chromosome) transgenic mouse expressing a Muc1-mCherry red fluorescent fusion protein (see Fig 1A for the targeting strategy). The fluorophore mCherry was chosen because of its fast maturation, high photostability (Shaner et al. 2004) and compatibility with GFP based reporters. We established three transgenic lines which were viable, fertile, and phenotypically normal (SFig 1-2).

**Figure 1:**
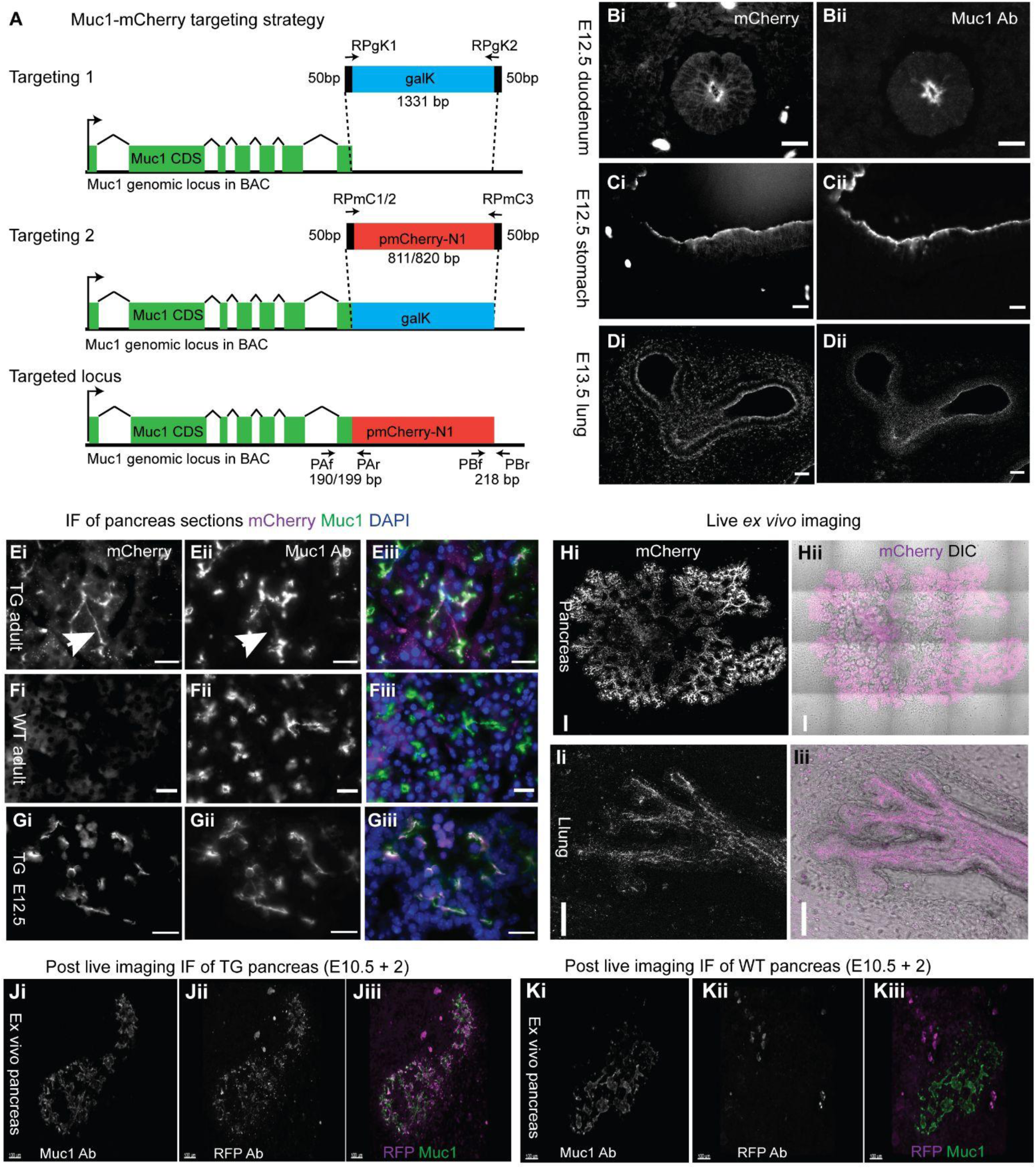
Generation of the Muc1-mCherry reporter line and validation of the reporter’s representation of endogenous Mucin expression. **A:** Schematic of the targeting strategy for the reporter construct which consists of the mCherry sequence fused to exon 7 of the *Mucin1* gene. Primer sequences are found in STable 5 **Bi-Gii:** Direct mCherry detection in TG embryonic tissue by fluorescence microscopy after fixation matches MUC1 antibody staining in E12.5 duodenum (Bi-ii), stomach (Ci-ii) and E13.5 lung (Di-ii). Ei-Giii: Direct mCherry detection by fluorescence microscopy after fixation in adult TG (Ei), but not wildtype (WT) (Fi) and embryonic TG (Gi) pancreatic epithelium also matches MUC1 antibody staining (Eii-Gii), with exception of a few duct-like structures which are not detected with the MUC1 antibody (arrowhead). **Hi-ii:** Stitched confocal mosaic tile scan from a whole E12.5 pancreatic explant after 3 days *ex vivo* culture. **Ii-ii:** Confocal scan from the tip of a E9.5 lung bud after 6 days *ex vivo* culture. **Ji-Kiii:** Maximum intensity 3D projection of confocal images of pancreatic TG (Ji-Jiii) and WT (Ki-Kiii) explants cultured for 2 days after explanting at E10.5 and fixation and staining for RFP and MUC1. Bar is 20 µm in B-G and 100 µm in H-K. DIC= Differential Interference Contrast. IF = Immunofluorescence. Ab = Antibody.

**Figure 2:**
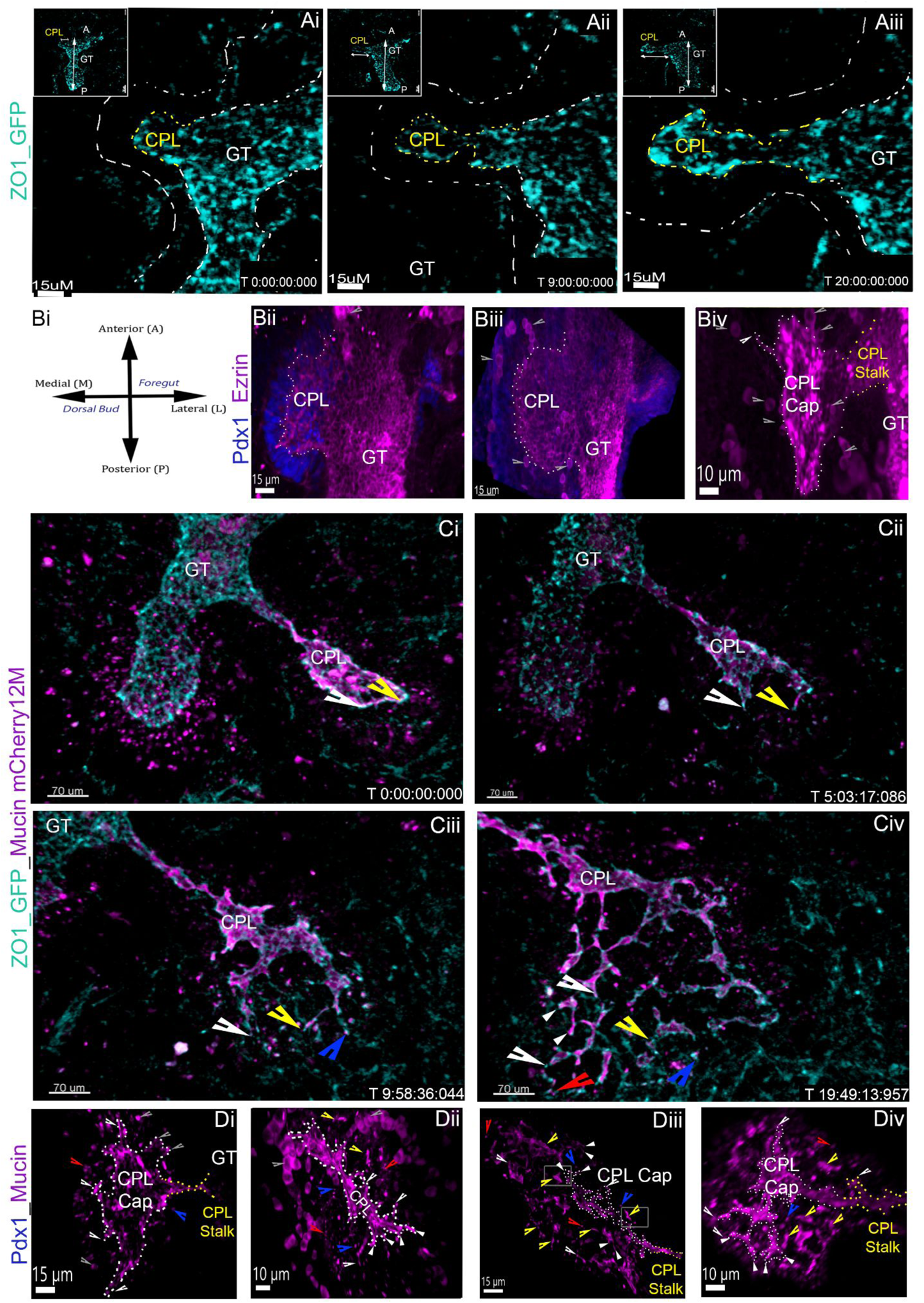
Lumenal network transformation is comparable in vivo and ex vivo. **Ai-iii:** Representative time frames from 4D imaging of a ventral pancreatic bud expressing ZO1_GFP reporter (Cyan). The TIFFs depict CPL budding out from and perpendicular to the foregut. White dashed outline estimated the outer epithelium, inner dashed lines mark the foregut, yellow dashed line marks the CPL. Inset shows an overview of the entire pancreatic explant with the direction of growth (A, anterior, P, posterior) marked by arrows. **Bi** Is a schematic to help orient the WMIF images in panels B and D, **Bii-iv:** MIP of embryonic foreguts (respectively at E9.0, E9.5 and E10.0) depicting equivalent lumenal topology observed in the 4D imaging of CPL budding (see A). **Ci-iv:** Representative time frames from 4D imaging of a ventral pancreatic bud expressing the Muc1-mCherry (magenta) and ZO1_GFP (cyan) double reporter. The TIFFs depict the transformation of the CPL into a secondary primitive plexus network. Arrows/spots highlight examples of different transformative topological events observed in the 4D imaging dataset (white large open = extension, white small closed = bifurcation, red = De Novo, Blue = BPP, Yellow = BPT). **Di-iv:** MIP of embryonic foreguts (i and ii E10.5, iii E11.0, iv E11.5) depicting equivalent lumenal topology observed in 4D imaging of a CPL transforming into a secondary network (C). Arrowheads denote the same structures detailed in Ci-iv. Grey boxed areas in Diii highlight the close proximity between the branching CPL and secondary lumenal site that are likely a static depiction of BPT. Gray arrows in panels B and D are used to point out autofluorescence from blood cells.

Muc1-mCherry was detectable in the pancreas, intestines, stomach and lung (Fig 1B-M), where it was expressed on the apical membrane of epithelial cells, closely resembling MUC1 antibody staining (Fig 1Fi-Kiii). We did occasionally observe structures (arrowhead in Fig 1I) that were Muc1-mCherry positive, but did not stain with the MUC1 antibody. As this fluorescence was found along ductal structures, we ascribe this difference to a higher sensitivity of the reporter.

To establish that introduction of the BAC had no effects on pancreatic architecture, differentiation, and function, we quantified whole mount immunofluorescence (WMIF) stainings for MUC1, Insulin (INS) and E-cadherin (CDH1) in E14.5 pancreata. We did not find significant changes in transgenic (TG) embryos as compared to wild type (WT) littermates (SFig 2). We further confirmed that the fasting blood glucose of TG mice was normal (SFig 1B, D, F, H).

We successfully imaged Muc1-mCherry in live explanted embryonic pancreas and lung by confocal microscopy (Fig 1Hi-Iii). Live-imaging of *ex vivo* cultured dorsal pancreas from each substrain revealed a lower Muc1-mCherry expression in the 0M substrain, and a delayed onset in the 5M substrain (results not shown). However, the 12M and 2F substrains had comparable, strong, Muc1-mCherry signals, detected in pancreatic explants that reflect the *in vivo* onset of endogenous MUC1 expression (Movie 4, Figure 2C and D). Pancreas ductal development and branching appeared normal in explant cultures from all substrains at all stages of culture compared to explanted pancreas from reporter-negative littermates (Fig 1Hi-ii and Ji-Kiii and SFig 2F-G). We henceforth proceeded using the 12M substrain, unless otherwise denoted.

### The pancreatic lumen is derived from the foregut lumen, but acquires distinct properties after budding

First, we aimed to define the spatiotemporal dynamics of CPL budding. To visualize lumens in early E9.75 to E10.0 pancreata we established an imaging protocol using foregut explants (see methods) from the Muc1-mcherry reporter and/or a reporter for zonula occludens-green fluorescent protein (ZO1-GFP) (Foote, Sumigray, and Lechler 2013). The ZO1-GFP reporter marks the apical domain in cells with immature apical polarity (Villasenor et al. 2010; Hick et al. 2009) and tight junctions in cells with mature apical polarity (Foote et al).

A hierarchy of apical protein acquisition in the CPL was evidenced by comparing ZO1-GFP and Muc1-mCherry during pancreatic evagination (Movie 2). ZO1-GFP was expressed throughout the CPL during its linear growth perpendicular to, but continuous with the foregut (Movie 1*, Fig2Ai-iii)*. This budding morphology of the CPL *ex vivo* is similar *in vivo* (Movie 3, *Fig 2Bii-iv*). In the ZO1-GFP live imaging a ‘fast moving, extra epithelial’ ZO1-GFP signal can also be observed (green arrows in Movie 1). This signal stems from endothelial cells, as verified by PECAM staining (red dashed outline in *SFig 5Ci* (ZO1-GFP) vs *SFig5Bii* (PECAM). In contrast to ZO1-GFP, Muc1-mcherry expression does not appear until after outbudding (Movie 2 and *SFig3Ai-ii*), when it is first detected in the distal CPL of a dorsal bud (*SFig 3Bi & iii*), overlapping with Zo1-GFP (*SFig 3Bii & iii*). In the newly evaginating ventral bud (*SFig 3Ci-iii*), there is minimal or no detectable co-expression. This acquisition pattern was verified *in vivo* by WMIF staining at E10.0 where MUC1 was detected in polarized cells most distal to the foregut (i.e the cap of the CPL) (*SFig 3Dii pink dashed line*). Conversely, ZO1-GFP (*SFig 3Dii* (*green dashed line*) & *SFig 4Ai*) and the apical proteins Ezrin (EZR) (*Fig 2Bii-iv,* Movie 5 & *SFig4Aiii*), and protein kinase C iota (PRKCi) (*SFig4Di)* in fixed pancreata also showed continuous apical expression from CPL cap (white dashes), to stalk (yellow dashes) to foregut. Interestingly, the CPL’s patency is variable across all morphological stages (S*Fig 4D-G*).

**Figure 3:**
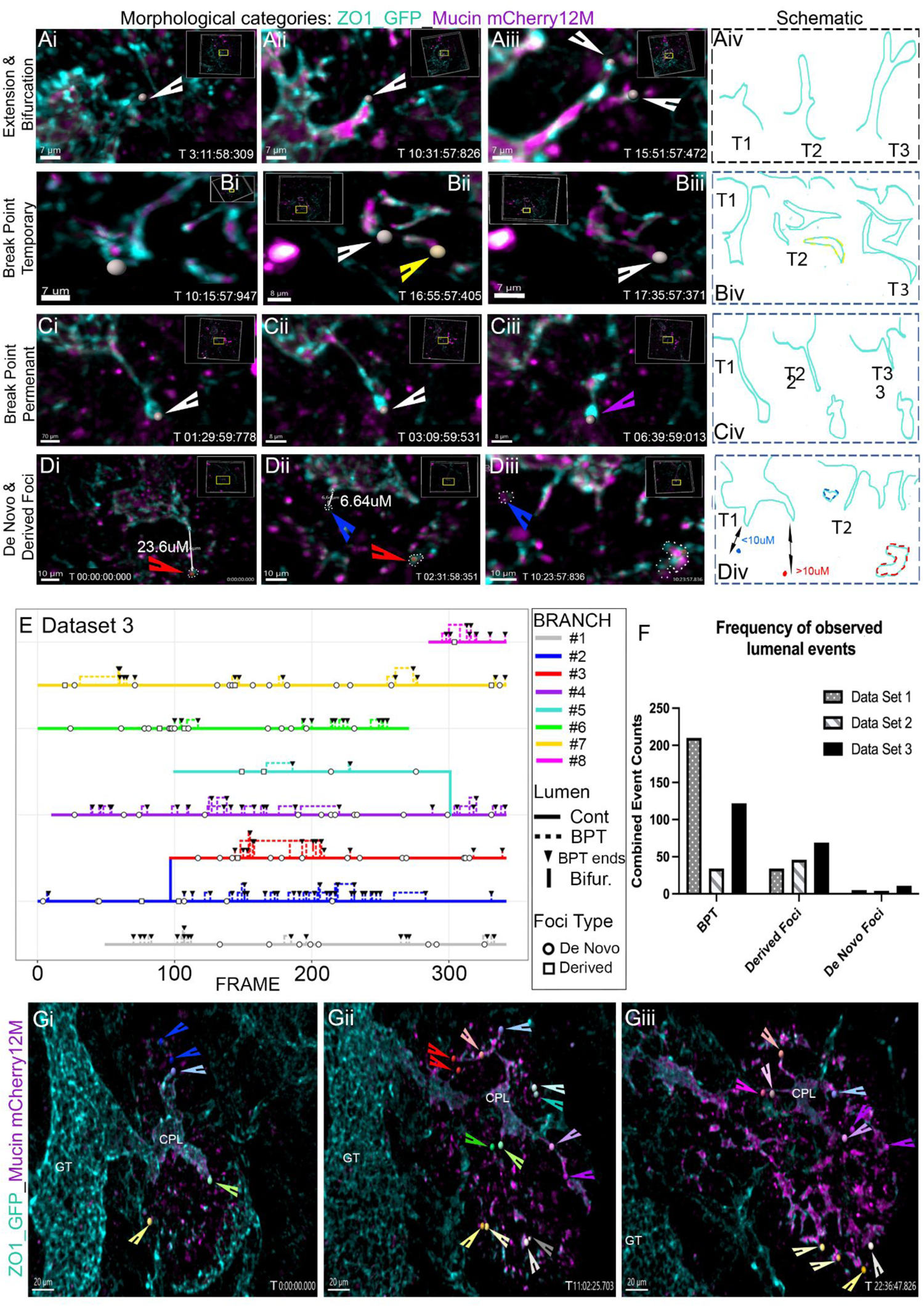
The pancreatic lumenal network is largely derived from the central primary lumen through expansion and rearrangement of apical membranes. **A-Di-iii:** Representative time frames from 4D imaging of the remodeling lumenal pancreatic network expressing ZO1_GFP (Cyan) and Muc1-mCherry (magenta). The TIFFs depict the different lumenal events occurring during network growth. (A) extension (B) temporary break points (C) permanent break points (D) derived and de novo foci. A-Div are skeletonised lumenal schematics summarizing the lumenal events. Arrows and spot colours highlight different lumenal events: white = the original lumenal branch, or fragment, yellow is a BPT, purple is a BPP, red is a de novo foci and blue is a derived foci. In panels D the tracked foci are further highlighted by a dashed outline. **E:** A tracking plot from dataset 3 (for dataset 1-2 see SFig 6) displaying the frequency of tracked lumenal events, occurring over time, to contribute to the proximal distal growth and transformation of discrete lumenal branches). The plot visualizes the high frequency of lumenal breaking events (dashed line, downward arrows define the point at which the lumen unifies) and derived foci (unfilled circles) are observed during lumen growth. In contrast de novo foci (unfilled square) occur relatively infrequently. **F:** Bar chart displaying the quantified frequency of each lumenal event observed in the 3 tracked live imaging datasets (BPP 35/570). G**i-iii:** Representative time frames from 4D imaging of the remodeling lumenal pancreatic network expressing ZO1_GFP (Cyan) and Muc1-mCherry (magenta). The TIFFs shown are a visual representation of the quantitative tracking data shown E and data set 3 in F. Each spot colour represents a different lumenal branch, light spots mark lumenal extension, dark spots mark breaking events and foci (Imaris tracking spots are further highlighted with an arrow of the same color).

**Figure 4:**
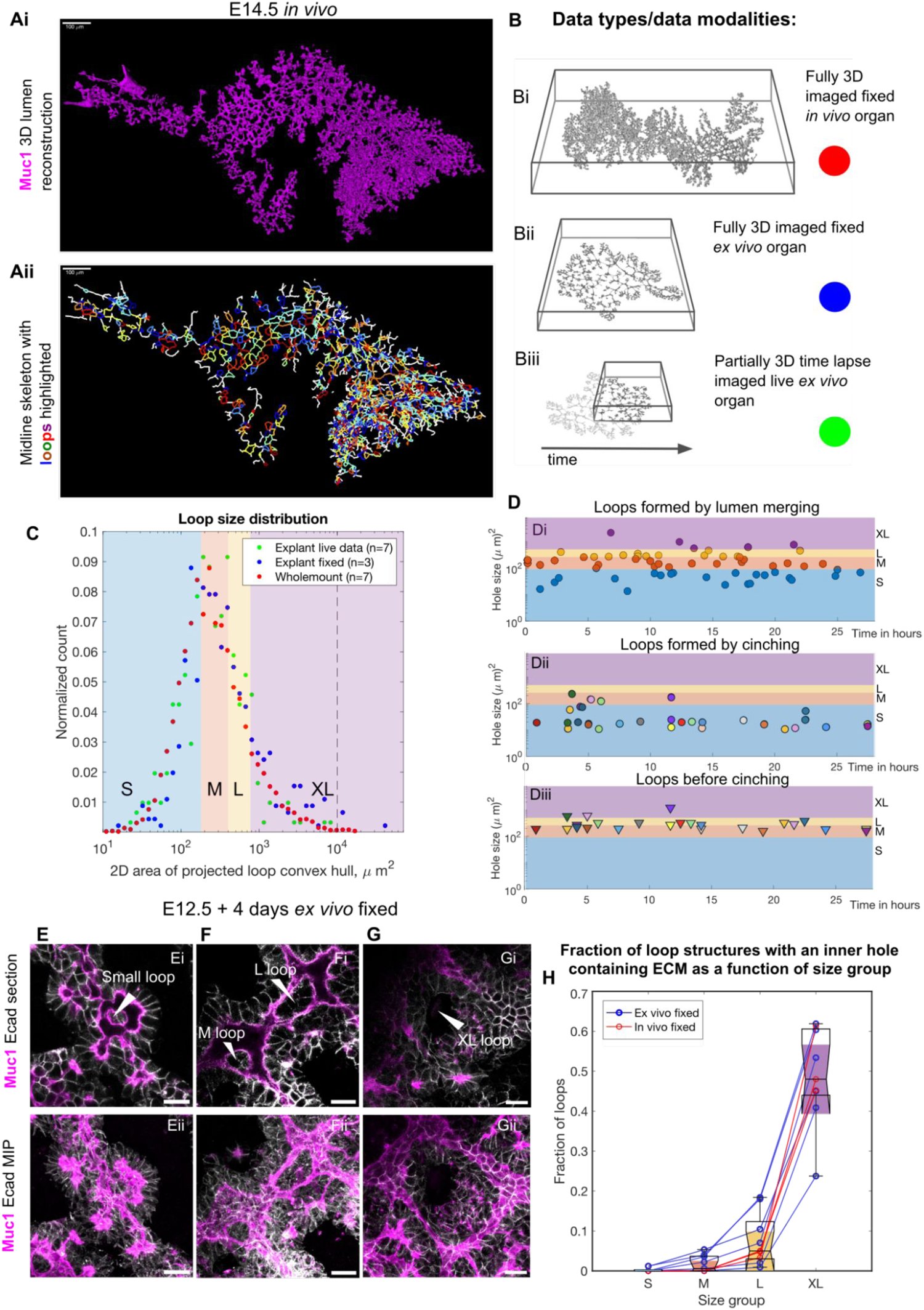
Characterization of loops and loop formation. **A:** Random forest based segmentation and 3D reconstruction of Muc1 stained E14.5 *in vivo* pancreas (upper) and corresponding midline skeleton with loops highlighted in different colors (lower). White bar is 100 µm. **B:** Cartoon illustrating the three different samples types used for analysis of loops: Bi) Muc1 and Ecad stained E14.5 WT fixed *in vivo* pancreas Bii) Muc1 and Ecad stained fixed explants (*ex vivo*) from E11.5 WT embryos cultured for 3 days, Biii) Muc1-mCherry pancreas explanted at E12.5 and grown *ex vivo* for 48 hs, followed by live time lapse imaging for up to 48 hs. For the latter, only approximately one quarter of the total pancreas was kept in the field of view and tracked. We used a random forest based method for segmenting lumens in 3D images of fixed and immunofluorescence stained samples and a semi-automated feature based method for segmenting lumens in 3D live imaging data. **C:** Loop size distribution for the three sample types, measured as the projected area onto the 2D xy plane of the loop formed by lumen midline. The size classes Small (S, blue), Medium (M, orange), Large (L, yellow) and Extra Large (XL, purple) are color coded in subsequent figures. The count is normalized by the total number of loops observed in each size bin. n=7/3/7 in vivo/ex vivo fixed/ex vivo live samples. Black dotted line signifies approx. upper size limit for loops in *ex vivo* live due to field of view limitation. **D:** Quantification of cinching and loop neoformation in n=7 movies of *ex vivo* live samples. (Di and Dii) hole size is here the size of the loop right after formation. Loops before cinching and loops formed by cinching have the same color when they correspond to the same cinching event. (Diii) hole size is here the size of the loop right before cinching occurs. **E-G:** Examples of loops from different size categories from ex vivo samples explanted at E12.5, grown for 4 days, and stained for Muc1 and CHD1. The upper row shows a single z-plane from the middle of the loops. The lower row shows a maximum intensity projection. Arrowheads point to the inner holes in loops, except for the XL loops which have epithelial cells and thus no inner holes. The bars are 20 µm. **H:** Quantification of loops without epithelial cells in the central hole of the loop as a function of size group in *ex vivo* fixed (n=3) and *in vivo* fixed (n=7) samples. Size group S is significantly different from group L and XL, and size group M is significantly different from XL (Krusksal Wallis test P<0,05).

**Figure 5:**
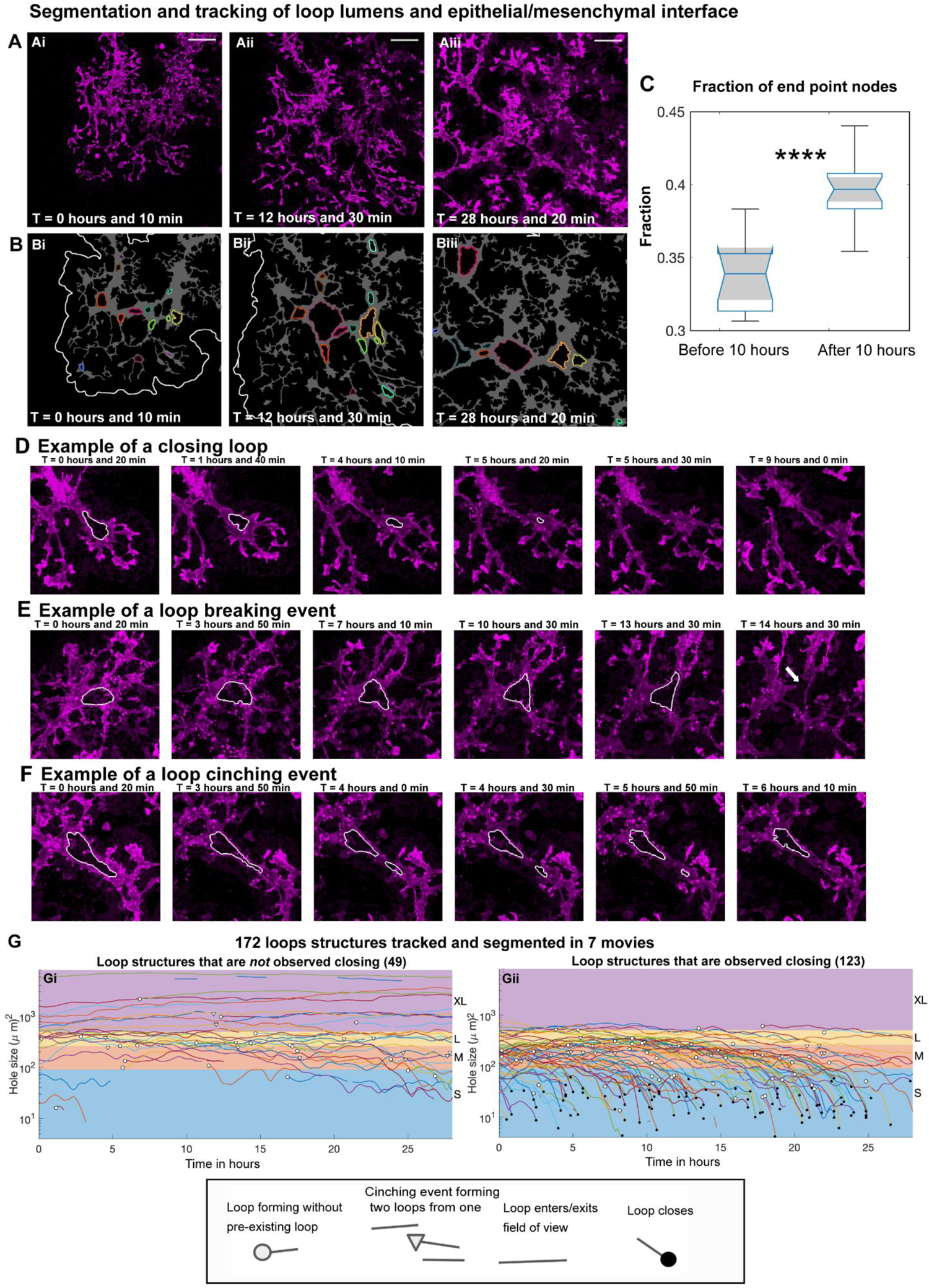
Tracking of ductal loop structures in mid-gestation pancreas. **A-B :** Tracking loop structures and segmenting lumen plus epithelial/mesenchymal border in live imaging of pancreatic explant culture imaged every 10 min, (MIP from one out of 6 experiments/movies shown in C and D, (corresponding to light blue dots/Exp. 1) at (Ai) T= 0 hours 10 min, (Aii) T = 12 hours 30 min and (AiiI) T = 28 hours 20 min). **B:** Same data as A but showing lumen segmentation in gray, tracked loop structures with unique color ID (same color across time means same loop). Note that there may be loop structures besides those highlighted, the ones highlighted have all been inspected and verified as bonafide loop structures in 3D. For some parts of the tissue, the thickness was too high and signal thus too low to properly distinguish loop structures. White lines are approximate placement of epithelial mesenchymal interface. **C:** Analysis of the ratio between number of end points and branch points before and after 10 hours of imaging, showing that the fraction of nodes that are end points go up with time. n=6 experiments. p= 9.23e-6 with Wilcoxon rank sum test. **D:** Example of a gradually constricting loop structure that ends up closing. Magenta: MUC1. White line: segmentation of loop. **E:** Example of a loop breaking event. Magenta: MUC1. White line: segmentation of loop. **F:** Example of a loop ‘cinching’ event’. Magenta: MUC1. White line: segmentation of loop. **G:** 172 loop structures segmented and tracked in 7 movies (of 5 different explants). Of these, two movies have duration 50 hours, five have duration 28 hours and 30 min, imaging was done every 10 min. A range of features of the loops projected onto the XY plane, where extracted over time: *’inner hole area’, ‘inner hole diameter’, ‘convex area’, ‘perimeter of convex hull’, ‘minor / major axis length of an ellipse fitted to the inner loop hole segmentation’, ‘lumen loop and hole total area’, ‘number of branch points/outgoing lumens’, ‘distance to epithelial/mesenchymal interface’, ‘mean lumen width’, ‘eccentricity of an ellipse fitted to the inner loop hole segmentation’.* Here we show loop trajectories of the size feature ‘*inner hole area*’ measured in (μ*m*)^2^over time, separated between trajectories where we observed loop closure (Ei) and trajectories where we did not (Eii). Hollow circle marks a loop forming event. Triangle marks a loop cinching event. A line ending/starting with no symbol means that the loop exits/enters the field of view. Full circle marks a loop closure event.

From these live imaging and *in vivo* data we conclude that the CPL forms from an apically polarized, patent lumenal tissue (the foregut) via budding (Schematic 8 A-B). During budding, the CPL acquires distinct properties as evidenced by apical domain maturation and reduced lumenal patency.

### The secondary lumenal network is established primarily through remodeling of existing apical domains

Our next aim was to define and quantify how the multi-lumenal secondary network, which forms the basis of the pancreatic ductal system, is generated. Specifically, we wanted to test the current model (Hick et al. 2009; Villasenor et al. 2010; Kesavan et al. 2009), which states that the secondary lumenal network forms by fusion of peripheral, de novo generated foci.

The first event consistently observed in all datasets after the completion of budding was a remodeling of the CPL (Movie 4). Over time, the ellipsoid CPL (*Fig 2Ci* & *S Fig6Ii*) forms unilateral and bilateral i.e. bifurcation (white and yellow arrowheads *Fig 2Di-iv*) extensions to create a branched structure (*Fig 2Civ & S Fig6Iiii*). These extensions form multiple lumenal branches some of which remain in contact with the foregut lumen via the CPL stalk (Movie 4 & 6). WMIF staining confirmed that transformation and extension of the CPL was also evident *in vivo* (*Fig 2Di* & *S Fig4Bi-iv*). We therefore conclude that CPL extension is the first event in the establishment of a secondary lumenal network (Schematic 8 C).

The next dynamic transition observed was a change in the connectivity of the lumenal network. We observed CPL extensions transitioning between unified and disconnected (yellow arrow in Movie 4 & *Fig 2Cii-iv*), with lumenal break points which were either temporary (break point temporary, BPT - *Fig 3Bi-iv* & Movie 6) or permanent (break point permanent, BPP - *Fig 3Ci-iv* & Movie 6). In concurrence with previous studies, we also detected apical foci at distance and proximal to the CPL. However, our quantifications show that most foci appear proximal to a lumen and are unlikely to be formed through *de novo* cell polarization. Foci that first appear within a distance of one average cell length (<10 µm, see materials and methods and *SFig5D-E*) from a lumen we categorize as derived, whereas those at a longer distance, which may not be derived, we categorize as “*de novo”* (*Fig 3Di-iv* & Movie 6). The foci also displayed varied behaviors and did not simply coalesce to establish a secondary network (Movie 4). BPTs, BPPs and derived foci are all examples of apical domain remodeling events that transform the growing lumenal network.

To determine how frequently remodeled and *de novo* apical domains occur during network growth we tracked these events with 4 minute intervals (Movie 7), and integrated the dynamic data into a tracking plot (*Fig 3E (Dataset 3), SFig 6F & G (Datasets 1 & 2)*). Overall, there was no consistent pattern in the frequency of events between branches (*Fig 3E SFig 6F & G*). Importantly, we found that events remodeling existing apical polarity i.e. BPTs (dashed lines) and *derived* foci (circles) were much more frequent than contributions from *de novo* foci (squares) (*Fig 3E, SFig 6F & G*). Of the total events tracked and counted, 64% were BPTs, 6% were BPPs, 26% were derived foci and only 4% were *de novo* foci (chi squared test df 67.26, P value <0.0001). While permanent lumenal break points (BPP) were limited, each initiated a new unique site of network growth (*Fig3Ci-iii*, Movie 6). While rare, *de novo* foci had a much longer (10 fold) duration (*SFig 6C*) than BPTs (*SFig 6A)* and derived foci (*S Fig6B*), emphasizing the need for high frequency imaging. WMIF *in vivo* stainings reflected the transitions observed in 3D timelapse data i.e from single CPL (Movie 5, *Fig2Di-ii* & *S Fig4Ai-iv*), to multiple disconnected lumenal sites (Movie 5, *Fig2Diii* & *S Fig4Ci-iv*), to a more unified structure (Movie 5 & *Fig2Div*). We therefore find it credible that many of the ‘disconnected’ lumenal sites (yellow arrowheads *Fig 2Dii-iv*) and foci (blue and red arrowheads *Fig 2Di-iv*) observed in fixed tissue are static depictions of dynamic remodeling events, rather than *de novo* generated lumens. We thus conclude that the secondary lumenal network is primarily established through remodeling of existing apical domains.

**Figure 6:**
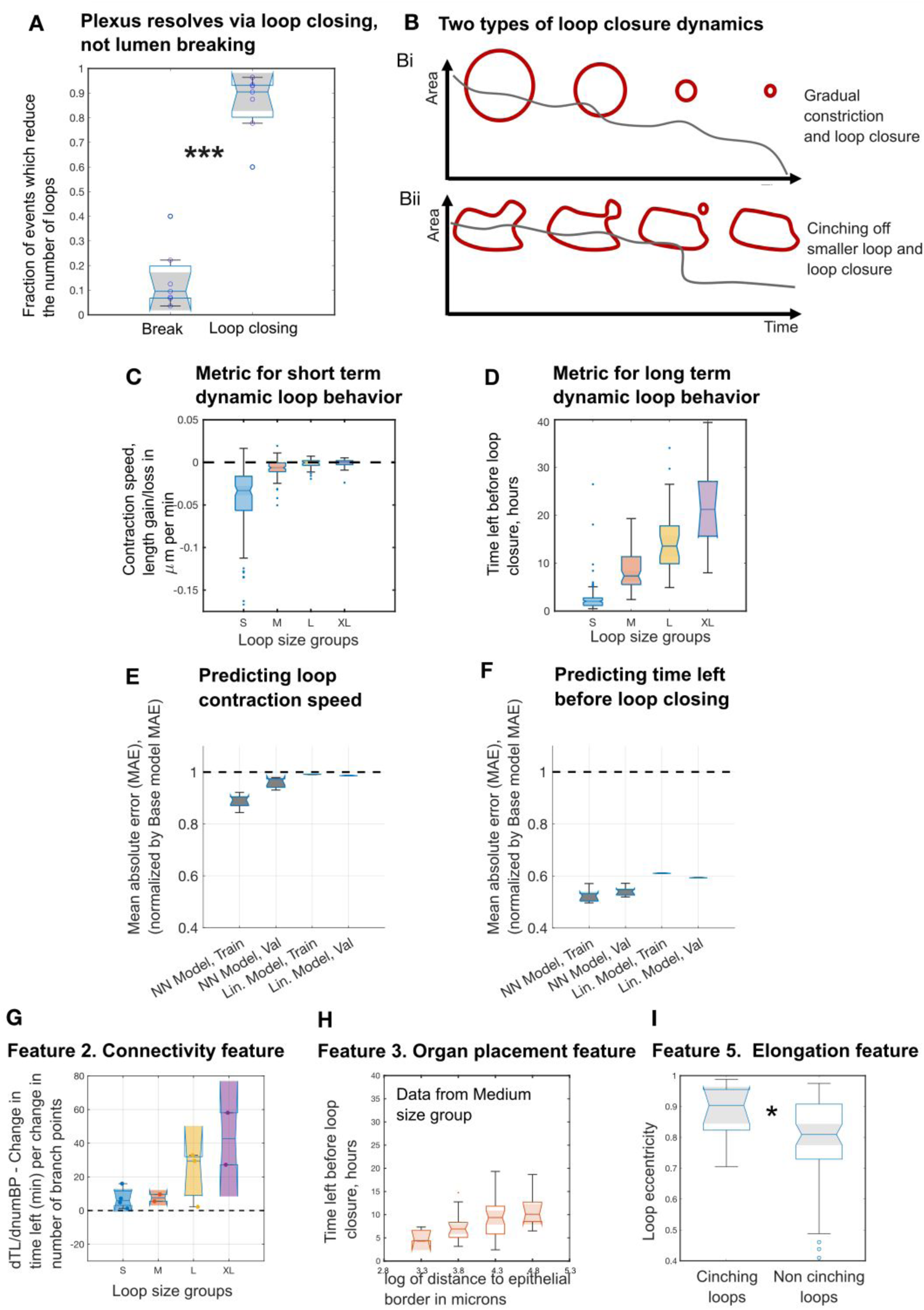
Loop closing dynamics in mid-gestation pancreas. **A:** Events that reduce the number of loops in the network are lumen breaking events (if the lumen breaking is participating in a loop structure) and loop closing events. These two types of events were counted for n = 7 different explants (data gathered from 14 movies. 5 explants have 2 positions/movies, 1 explant 3 positions/movies and 1 explant have 1 position/movies). Distributions pass Anderson-Darling normality test so p value = 2e-7 is found via two sample T test. **B:** Cartoon of the two types of loop closure dynamics. (Fi) Gradual constriction followed by loop closure (Fii) cinching off smaller loop that then undergoes loop closure. **C:** Y axis: Metric for short term dynamic loop behavior - *‘Mean loop contraction/expansion rate in cell loss/gain per day’* (contraction speed of loop perimeter per 6 microns, which is the average length of one cell junction contributing to the loop circumference), X axis: Loop size groups. **D:** Y axis: Metric for long term dynamic loop behavior - *‘Time left before loop closure in hours’*. X axis: Loop size groups. **E-F:** Mean absolute error (MAE) on training/validation data for linear and neural network models predicting ‘Time left before loop closure’ and ‘loop contraction speed’ normalized by base model MAE. **G:** We estimated the change in time left when the number of branch points is changed (i.e. finding the slope dTL/dnumBRP of a linear fit) within even smaller loop size groups, than S, M, L and XL, in order to make sure that the correlation we observe between time left and number of branch points is not caused by the strong correlation between number of branch points and loop size. We see that these slopes were consistently positive as expected, if the effect still holds even when loops of the same size are compared. The colored ‘notch’ areas show the 95% confidence intervals of the median slope for each size group (S, M, L and XL). None of these confidence intervals cross zero which shows that a higher number of branch points is associated with slower closure. Note that the apparent trend that L and XL size groups have a steeper slope is not significant. **I:** Mean eccentricity of loops in size groups L and XL (mean is taken over time) is higher in loops that go through one or more cinching events during the time course, P value is 0.0262 (Wilcoxon Ranksum test). **H:** We found a positive correlation between *time left* and *mean distance to epithelial/mesenchymal interface*. Since there is also a slight positive correlation between category and distance to the organ edge we look at correlation in our size categories separately (here is shown group M, see SFig 10H for S, L and XL). The strongest effect of placement in the organ on *time left* seems to be in the M and L groups, where Spearman’s rho is >0.4. Number of time courses in each category is, Small: n=114. Medium: n=72. Large: n=33. XL: n=9.

### Epithelial loops of varying sizes provide unique pancreatic progenitor niches

Once the naive primitive network is generated we observe a gradual transition to a more unified structure. Examples of this re-connectivity can be seen in the tracking plot *Fig 3F*. The next lumenal network stage is termed the plexus; an interconnected multi-lumenal structure containing many loops in the trunk region. Here, we analyze the transformation and cellular architecture of the lumenal plexus (as illustrated in Fig 8 F-H). Loop formation by lumen merging was observed at all stages, post CPL transformation (e.g from ca E10.75 onwards). Examples of loop formation at early stages can be seen in *SFig 7A-C* and movie 8 and at later stages in *SFig 7D* and Movie 9. Thus, we conclude that loop formation is not specific to a particular stage of secondary network development.

**Figure 7:**
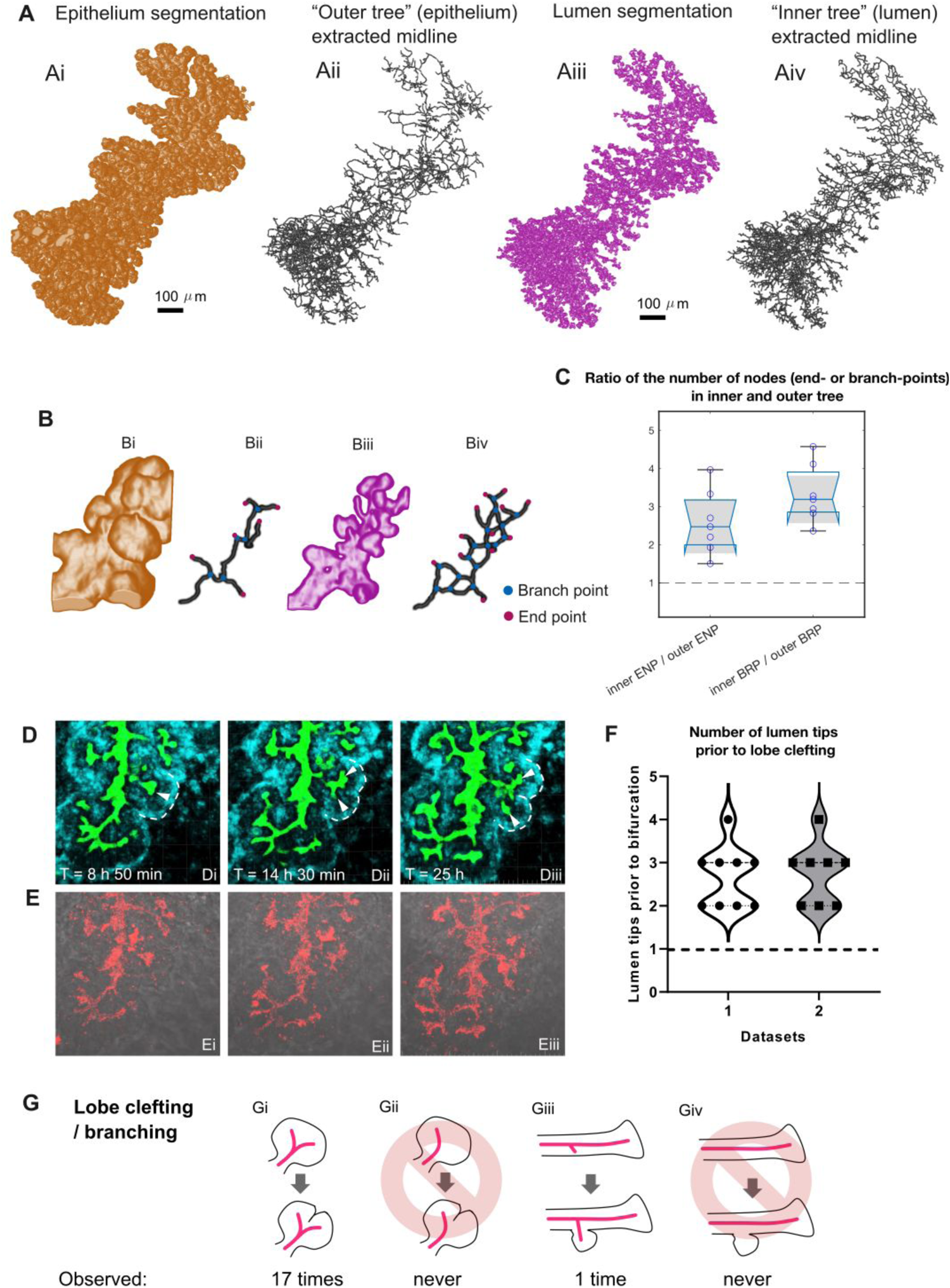
Comparative analysis of epithelial and lumenal networks. **A:** (Ai) - Segmented and 3D reconstructed epithelium E14.5 *in vivo* pancreas. (Aii) - ‘Outer tree’, corresponding midline skeleton of epithelium segmentation. (Aiii) - Segmented and 3D reconstructed lumenal network E14.5 *in vivo* pancreas. (Aiv) - ‘Inner tree’, corresponding midline skeleton of lumen segmentation. **B:** Bi - detail of epithelial branch, Bii - epithelium midline with branch points (BRP) and end points (ENP) highlighted in blue and pink, Biii - detail of lumen in same branch, Biv - lumenal midline with branch points (BRP) and end points (ENP) highlighted in blue and pink. **C:** The number of endpoint point (ENP) nodes in ‘inner tree’ per ‘outer tree’ (ENP) (median ∼2.5) and the number of branch point (BRP) nodes in ‘inner tree’ per ‘outer tree’ (BRP) (median ∼3.2). In vivo fixed. N=7. **D-E:** Example of lobe clefting event. Di-Ei: T = 8h 50 min. Dii-Eii: T = 14h 30 min. Diii-Eiii: T = 25. **D:** Green: MIP of probability of pixel lumen class predicted by Random Forest. Blue: MIP of probability of pixel epithelial/mesenchymal interface class predicted by Random Forest. White dashed line: highlighting a clefting lobe. White arrows: lumenal tips. **E:** Red: Raw Muc1 signal (appears . Gray: Raw DIC channel. **F:** Count of the number of lumenal tips present in lobes before clefting. **G:** We tracked branching events in the outer tree and backtracked the inner tree to the time point immediately preceding an outer three branching event and recorded the number of lumens in two live imaging datasets of Muc1-mCherry E14.5 +2 days culture dorsal pancreata. We here define clefting as the bifurcation (type Gi and Gii event) of a pre-existing epithelial lobe, and lateral branching (type Giii and Giv). In the first dataset, 9 type Gi/Gii clefting events and no type Gii/Giv events were observed in the outer tree consisting of 5 epithelial main branches. In the second dataset, 8 Gi/Gii clefting events and one lateral branching type Giii/Giv event was observed in the outer tree consisting of 2 epithelial main branches. All type Gi events were preceded by at least two lumenal tips inside the lobe. We observed only one type Giii event, which was preceded by one lumen protruding at the branching site. For total 17 type Gi events, the number of lumen tips observed inside lobes before clefting was [3 2 2 3 4 3 3 2 2 2 3 3 4 3 2 3 2], mean-+std = 2.7059+-0.686, median = 3, min = 2, max = 4.

**Figure 8:**
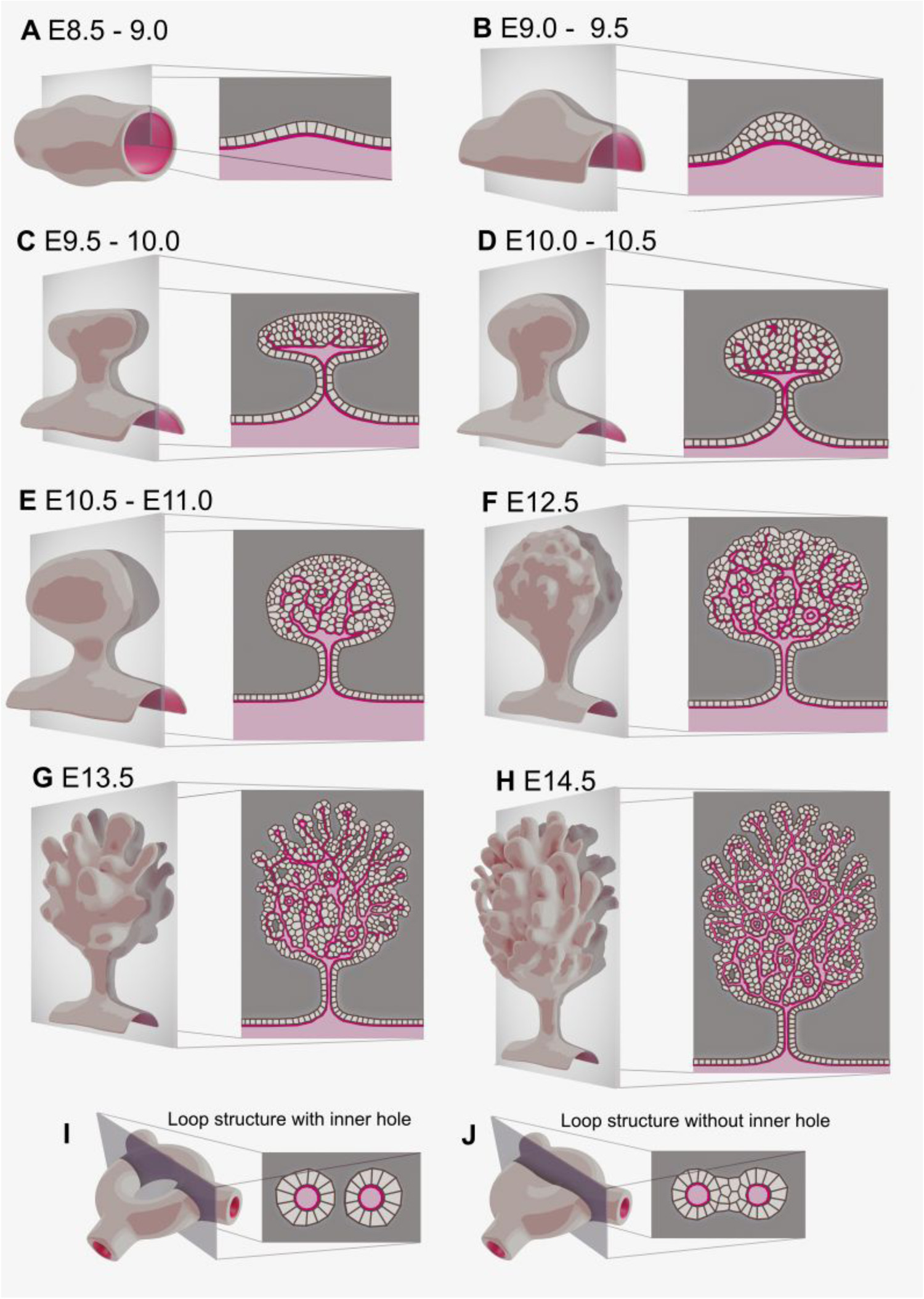
Schematic illustration of pancreas morphogenesis. Each embryonic stage is represented by a 3D illustration showing the general structure of the pancreas (left side) and a 2D section showing cellular detail (right side) through the indicated optical plane. Dark red color denotes apical membrane, light red color marks the inside of the lumens/tubes. The light gray cells are epithelial/pancreatic cells, while the dark gray background color denotes non epithelial tissue. A-B : The pancreas starts as an outbudding of the endodermal foregut tube at E8.5-9.0, causing the apical membrane of multipotent pancreatic progenitor cells to form a central primary lumen (CPL), as a perpendicular extension of the foregut lumen. C: The CPL does not remain uniformly open (patent) at the stalk or cap. The secondary lumenal network starts as extensions of non-patent apical polarity domains extending from the CPL. D: The apical polarity extensions start to break apart and form secondary lumenal sites (foci and small plaque like lumens). E: The disconnected lumens and foci begin merging with each other and with the CPL extensions forming loops and branches. F: The lumenal network is fully connected and contains loops. G-H: Growth of the lumenal network, loop closing, and branching of the epithelium leads to a stage (around E14.5) with a branched periphery (tip domain) and a plexus containing center (trunk domain). I: An example of the type of ductal loop with non epithelial tissue present in the center of the hole, which tends to be larger and persist for a longer time. J: An example of the type of ductal loop with only epithelial cells present in the hole, which tends to be smaller and close faster.

To define the size and spatiotemporal distribution of loops within the plexus, we focused our analysis on midgestation pancreata where loops are frequently observed (*Fig 4A*). We segmented the lumenal network in three different sample types (illustrated in *Fig 4B*): Fixed *in vivo*, live *ex vivo,* and fixed *ex vivo* pancreata. Segmentations of the MUC1 IF signal (*Fig 4A*) or Muc1-mCherry signal (*Fig 5A-B*) were used for identifying and manually tracking loops. Loops sizes range over four orders of magnitude (inner hole area approx. 10 - 100.000 µm^2^ (*Fig 4C*)) in all sample types. We categorized loops based on a combined size feature (see materials and methods) as Small (S, blue, 0-90 µm^2^ inner hole area), Medium (M, orange, 90-260 µm^2^), Large (L, yellow, 260-505 µm^2^), and extra large (XL, purple, from 505 µm^2^ and above) (*Fig 4E-G*). Formation of new loops occurs in all four size categories but is more common in thesmaller size groups: S (56,4%), M (30.4%), L (8.3%) and XL (4.9%). Loop structures form in two topologically different ways: through merging of two lumenal extensions (Fig 4Di, *SFig 7D* and movie 9) or through ‘cinching’ at the center of a preexisting loop, to generate two smaller loops (*Fig 4Dii-iii, 6D,* and movie 11). Lumen merging generates new loops in all size categories (*Fig 4Di*). Cinching however only generates new S-M loops (Fig 4Dii) while simultaneously leading to loss of larger loops (*Fig 4Diii*).

Since it has been hypothesized that loops provide a unique niche within which endocrine progenitors are specified (Bankaitis, Bechard, and Wright 2015), we next analyzed the expression of epithelial, mesenchymal, and ECM markers in loops by WMIF. The inner hole of S-L loops contained CDH positive epithelial cells, in an amount scaling with loop size (*Fig 4E-F and H, SFig 8B,C,E,F, and illustrated in Fig 8J*). In contrast, the inner hole of half of the XL loops was devoid of epithelial cells (*Fig 4G and H, SFig 8A and D, and illustrated in Fig 8I*). Optical sectioning of WMIF revealed that DAPI and fibronectin receptor integrin ɑ5 (ITA5) positive, CDH1 negative, mesenchymal cells were rarely found within the inner hole of XL loops (arrow in SFig 8Ai-ii), but were instead found on the outside of the pancreas (SFig 8Aiii-iv). ITA5 was also detected on the basal side of epithelial cells facing the basement membrane (arrowhead in *SFig 8Aii, Bii, Cii*). Instead, the cell-free part of the inner holes contains ECM. Fibronectin (FN) was consistently present in XL loops (arrowhead in *SFig 8Di-ii*), and in many M-L loops (arrowheads in *SFig 8Ei-ii*), but not observed in inner holes of S loops (arrowhead in *SFig 8Fi-ii*). FN was also deposited as fibrils in close proximity to the basement membrane of epithelial cells, and in the interstitial matrix (*SFig 8D-F*). Interestingly, some large ducts also contained FN deposits inside the lumen (arrowhead in *SFig 8Gi-ii*). Laminin (LN) was not detected in any loops in the trunk region (results not shown), but only in the basement membrane at the periphery of the pancreas (*SFig 8Hi-ii*). In summary, the inner hole of small loops provide an epithelial niche with little contact to FN, LN or mesenchymal cells. On the other hand, epithelial cells forming the inside layer of larger loops are exposed to fibronectin. Cells in the loop-sparse environment at the periphery of the pancreas are exposed to mesenchymal cells, fibronectin, and high levels of laminin.

### Loop closing resolves the ductal network

To address how the plexus resolves, we analyzed lumen remodeling in time-lapse data from Muc1-mCherry expressing mid-gestation pancreata (172 loops from 7 explants). An example of loop tracking and segmentation can be seen in *Fig 5A-B* and movie 10, and a graphical representation of all tracks in *Fig 5G*. The rate of loop removal was slightly higher than the rate of loop formation, resulting in a gradual net reduction in the number of loops over time (*SFig 9C*). Interestingly, while lumenal branch points decreased, end points increased (*Fig 5C* and *SFig 9A-E)*. This topological change indicates a non-branching type of network remodeling, as classical branching with bifurcation alone should not increase the relative number of end-points.

We next analyzed our data to identify the mechanism driving plexus to arborised network remodeling. We found that loop structures are primarily removed from the lumenal network by a new mechanism where loops are gradually constricted (*Fig 5D* and movies 12-13). We coin this mechanism “loop closure”. Loop removal through pruning or breaking of lumens was rarely observed (*Fig 5E* and movie 14) and occurred significantly less frequently than loop closure (*Fig. 5C*).

Out of the 172 tracked loops 72% closed during imaging. Even though the sizes of the tracked loop structures tended to fluctuate, there were clear trends in loop behavior for different size classes (*Fig 5G*). S loops mostly contract, and close fast after reaching 8-12µm. M-L loops fluctuate in size but overall tend towards closing. XL loops tend to maintain or even increase in size (*Fig 5G*, *SFig 9F* and movie 15). Only 7 (30%) of the loops that start off as XL are observed closing in the time span of our imaging; 3 of these have one or more cinching events on the way to closing (*Fig 5F*). Cinching produces two loops of smaller size than the original loop and thus both loop closing and cinching events contribute to the overall reduction of loop sizes over time (illustration in *Fig 6B*).

### Long term dynamic behaviors of loops can be predicted from static features

Based on the recorded loop closure dynamics, we hypothesize that the closure mechanism depends on physical forces acting on loop epithelial cells. In order to address this hypothesis, we tested if static loop features related to the local tissue forces experienced (Sugimura, Lenne, and Graner 2016), could be used to predict dynamic loop behaviors. We chose to use static features, as this would allow us to measure loop characteristics using *in vivo* samples. First we defined two dynamic target parameters: mean loop contraction (’Contraction rate’) as a metric for short term loop behavior and time left in hours before closure (’Time left’) as a metric for long term loop behavior (*Fig 7C-D*). Next, we extracted static loop features relating to size, shape, connectivity and placement in the organ, from individual time-lapse frames, in movies where the dynamic behavior of the loop is known (*SFig 11C*) and trained feedforward, fully connected neural networks (NN) for the regression of ‘Time left’ and ‘Contraction rate’. To avoid overfitting, we used a validation dataset for early stopping. We also fitted linear regression models, which due to their simplicity do not overfit. As a control benchmark for baseline performance, we include base models; constant models that always return the median target value of the training dataset, regardless of the input. We use the base model mean absolute error (MAE) to normalize MAE for the NN and linear models, (*Fig 6E and F*).

For the prediction of the short term metric ‘Contraction rate’, both NN and linear models have similar errors to the control base models (their NMAE is close to 1, Fig 6E). Thus, neither model is able to predict short term dynamic loop behavior. In contrast, for the long-term metric ‘Time left’, both NN and linear models are significantly better than the control base model, (their NMAE is 0.5-0.6). In conclusion, whilst loop behavior is highly stochastic, we were able to model and predict long term trends, using static loop features.

The models confirm our finding that loop size is important, but also identify that placement in organ and connectivity matter: The most important feature for predicting ‘Time left’ was the combined loop size feature (Fig 6D-C) and SFig 11C). The next most important feature for predicting ‘Time left’ was placement in the organ. We found that proximity to the epithelial-mesenchymal interface is correlated with faster loop closure (*Fig 6H*). The third most relevant feature for predicting loop closure was loop connectivity within the lumenal network. A higher number of branch points (indicating that the loop has more outgoing lumen connections) is associated with longer time left before loop closure (*Fig 6G*). Of note, mean lumen width within a loop did not correlate significantly with dynamic behavior (*SFig 10F*). From our extracted static feature data, we were also able to determine that loop shape is associated with a specific mode of loop closure: We found that elongated (high eccentricity) L and XL loops are associated with cinching, (*Fig 6I*). Since L and XL loops that do not contain epithelial cells in their inner hole, are on average less elongated (SFig 10D-E), these data suggest that cinching is less likely to occur when ECM is present in a loop. In conclusion, we find that static loop features related to size, organ placement, connectivity and geometry can predict long term dynamic loop behavior.

### Formation of the inner lumenal network precedes the outer epithelial branching

In our live imaging analysis of pancreatic lumenal network establishment and remodeling we have focused on imaging apical polarity, and therefore the “inner” architecture of the ducts (the “inner tree”, see *Fig 7A and B*). In order to compare our results with previous fixed imaging and modeling of epithelial branching, which are mostly based on epithelial outlines of ducts, we next compare this “inner” network topology, with the “outer” network topology, (i.e. the epithelial tissue architecture or the “outer tree”), (*Fig 7A and B*).

E14.5 fixed *in vivo* pancreata (n=7) were used to extract an inner and outer mid-line map from segmentations of MUC1 and CDH1 IF and differential interference contrast (DIC) images, (see methods). A comparison of the maps showed that the inner tree had on average 3.1 nodes per outer tree node. Of these, the ratio is 3.2 for branch points and 2.5 for endpoints (*Fig 7C*). Loops with holes inside also exist as loops in the outer tree, while loops with only epithelium inside are not detected as loops in the outer tree (*Fig7A and B*). These fixed data suggest that the inside tree is more branched than the outside tree.

To determine the hierarchy of inner and outer tree clefting, we tracked the changes in inner and outer tree topology in real time. We focused on the formation of new peripheral ductal branches/lobes in E14.5 +2 days cultured dorsal pancreata (n=2 datasets). Muc1-mCherry was used for detection of the inner topology and a combination of autofluorescence and DIC for detection of the outer epithelial borders (*Fig 7D-E*). We tracked the growth of the outer tree and preceding every outer clefting event we recorded the number of lumenal end points present in the inner tree. We define clefting as the bifurcation (type Gi/Gii event in Fig 7G) of a pre-existing epithelial lobe and lateral branching (type Giii/Giv event) as the generation of a new epithelial lobe from a straight branch (See *Fig 7G* for illustrations).

In the first dataset, consisting of 5 epithelial branches we detected nine clefting events and no lateral branching events in the outer tree (Fig 7D-G). In the second dataset, consisting of 2 epithelial branches, eight clefting events and one lateral branching event were observed in the outer tree (*Fig 7D-G*). All clefting events were preceded by at least two lumenal tips inside the lobe. We observed only one branching event, which was preceded by one lumen protruding at the branching site. For clefting events, the mean number of lumen tips observed inside lobes before clefting was 2.7 (Fig. 7F). We never observed branching of the outer network without a preceding change in number of lumenal tips.

However, we often observed the presence of multiple lumen tips in one lobe without a subsequent lobe splitting event. These data signify that preceding lumenal changes are necessary but not sufficient to trigger epithelial branching.

In conclusion, the lumenal inner tree is more complex than the outer epithelial tree. Branching events in the outer epithelial structure are always preceded by the formation of an inner lumen, suggesting that lumenal remodeling drives epithelial branching.

## Discussion

Our study documents lumenogenesis live across multiple overlapping stages of pancreatic development. We identify a number of new lumenal mechanisms that drive the transformation of the pancreatic ductal system from a simple tube to an interconnected arborized network. Our findings are summarized in a schematic illustration in Figure 8.

The generation of a new reporter mouse for live imaging of apical polarity was imperative for visualizing these novel mechanisms in lumen transformation. The Muc1-mCherry reporter faithfully recapitulates endogenous MUC1 expression in multiple epithelial organs, serving as a valuable tool for apical polarity research. As opposed to the Cre-inducible Crb3-EGFP apical reporter (Pan et al. 2015), we found no evidence of gain of function phenotype in Muc1-mCherry heterozygous carriers. This is likely due to the use of the endogenous *muc1* promoter and regulatory regions in the BAC construct. To capture the fluorescent reporter’s expression in real time we utilized an *ex vivo* pancreas explant culture protocol (Percival and Slack 1999). The explant’s growth, though slightly compressed in depth, supports cell differentiation and morphogenesis that is relative to an *in vivo* dorsal bud (Löf-Öhlin et al. 2017; Puri and Hebrok 2007). We further validated the pancreas explant model by verifying key findings in stage matched *in vivo* pancreata.

To our knowledge, our study is the first to analyze the early stages of CPL formation and transformation using lumenal markers in high resolution 3D and 4D detail. Utilizing ZO1-GFP and Muc1-mCherry reporters we find a hierarchy, respectively early and late, in apical expression during CPL formation. We confirm that the CPL forms through a process of budding, during which the CPL stalk remains continuous with and extends perpendicularly from the foregut lumen (see illustration in Fig 8A-C). Thus, we conclude that the CPL stems from an already polarized site. The process of budding is thought to contribute to the formation of other foregut endoderm primordia such as lung (Metzger et al. 2008) and thyroid (Pierreux 2021). However, while other budding primordia, e.g. the lung buds, retain a patent lumen we find that the CPL and the lumens it derives do not. We speculate that a lack of classical polarity hallmarks, such as tight junction localized ZO1 and non-patent lumens, could permit a more dynamic cellular behavior enabling the observed variety of lumenal transformations.

Current models of pancreas duct formation depict the immature pancreas network initiating *de novo* by microlumen formation at the periphery of the dorsal bud (Hick et al. 2009; Villasenor et al. 2010; Kesavan et al. 2009). These microlumens are purported to coalesce and unify into a lumenal plexus. However, only one other study (Azizoglu et al. 2017) to date has provided live data documenting apical foci contribution to the lumenal plexus. This study included imaging of later stage Crb3-EGFP explants (E11.5+1 day) and found that apical foci only contributed to network growth at the tips/periphery of the network at this time. In contrast, we observed foci contribution across the whole network during secondary network establishment.

Using live imaging, we detected isolated lumenal foci distant to the CPL (Fig 8D-E), which would appear “de novo” if visualized statically. However, by tracking network transformation in 4D, we demonstrate for the first time that contributing foci stem from a pre-existing lumen. We document and quantify six dynamic mechanisms of lumen transformation, of which *de novo* lumen formation contributed least (3.2%) to network growth and transformation. We conclude that whilst secondary network growth is complex, it is largely driven by the reshaping of already established lumenal structures. The founding transformative structure is the CPL itself which, as Hick et al observed in their static study, first forms naive secondary lumenal branches (Hick et al. 2009). In contrast to previous studies (Shih et al. 2016; Hick et al. 2009), we find that lumenal branch extension from the CPL and the formation of peripheral foci are connected processes. We observe apical foci that both stem from and then later re-contribute to forming lumenal branches. Moreover, continuous lumens can fragment, become disjoined, and reunify. To our knowledge, this is the first study to report a tubulogenesis process which combines budding, dynamic remodeling and foci contribution. Follow-up studies utilizing cell membrane reporters will be necessary to understand the cellular behaviors that contribute to each category of lumen transformation. We speculate that cell proliferation could drive network growth, but how this could contribute to the temporary breaking and reunification of lumenal sites remains to be defined.

Multipotent pancreatic progenitors (MPPs) that form the pancreatic buds are uniquely specified within the foregut to express a pancreas transcriptional profile (Jørgensen et al. 2007). We and others in the field (Azizoglu et al. 2017; Shih et al. 2016; Nyeng et al. 2019; Mamidi et al. 2018) have shown that MPPs also have unique cellular behaviors and properties. The primitive bud provides a MPP niche within which complex inner lumenal networks and outer epithelial branches are initiated. These structures establish an intermediate plexus stage within which endocrine cells are specified. Eventually, the plexus is remodeled to an arborised architecture within which MPPs acquire their mature exocrine or endocrine function. We show that the loops found in the plexus provide unique size dependent niches that differ in their closing dynamics and cell and ECM composition. This observation is especially interesting, given earlier studies showing that endocrine progenitors are preferentially specified in and depend on a plexus environment: Genetic perturbation of plexus formation decreases endocrine differentiation (Kesavan et al. 2009; Barlow et al. 2023), while prolongation of the plexus stage enhances endocrine cell numbers (Azizoglu et al. 2017). We speculate that a progenitor’s exposure to unique ECM profiles during loop remodeling could regulate endocrine differentiation. We aim to address this hypothesis using live imaging reporters of endocrine differentiation in future studies.

Our tracking analysis of mid-gestation ductal remodeling led to the discovery of a novel mechanism, which we term “loop closure”, that leads to a net reduction of loops. Although our data is influenced by growth of the explant, which pushes some loops out of the imaged frame of view, this loop reduction replicates what has been documented *in vivo* (Dahl-Jensen et al. 2018). We show that the ductal network is remodeled into an arborized structure by gradual closing rather than by pruning of the ductal loops, as has previously been suggested (Flasse, Schewin, and Grapin-Botton 2020)

One study has suggested that the network is remodeled based on eliminating lumens with the least liquid flux (Dahl-Jensen et al. 2018). If we assume that liquid flux is related to lumen width, we do not find any evidence in our live imaging data to support this hypothesis, as we do not observe any correlation between lumen width and loop closure. We instead hypothesize that loop closure depends on physical forces acting on the loop epithelial cells’ cytoskeleton depending on the size, geometry, and topology of the loop. In support of this hypothesis, we were able to predict a metric for long term loop behavior using models trained with features associated with physical force. We also observed that these features correlated with loop closing speed. Supporting this data, it has been shown that the plexus fails to remodel in models of perturbed actomyosin contractility (Azizoglu et al. 2017). We speculate that forces transmitted through the lumenal network connections - likely via actomyosin contractility - hinder loop closure and/or cause loop widening. Noticeably, the ability to infer dynamic behavior based on static features also allows us to bridge a bioimaging gap between low resolution dynamic *in vitro* data and high resolution, multiple-marker, *in vivo* data.

Branching morphogenesis studies of the pancreas tend to focus either on epithelial branching morphogenesis or internal lumen remodeling. We aimed to reconcile these studies by tracking lumen and epithelium in parallel. Early live imaging studies using the Pdx1-GFP reporter concluded that the predominant mode of early stage pancreas growth is lateral branching (85%), with some bifurcation (15%) (Puri and Hebrok 2007). We tracked epithelial branching at a slightly later stage and predominantly found bifurcations, which is in agreement with another recent live imaging study (Darrigrand et al. 2024). At the cellular level, epithelial branching has been explained by localized proliferation, coordinated cell shape changes, and/or cell rearrangements (Varner and Nelson 2014). In the pancreas, proliferation happens preferentially at the tip during branching (Sznurkowska et al. 2018; Zhou et al. 2007; Larsen et al. 2017), but a direct coupling between proliferation and branching has not been reported. Studies manipulating cytoskeletal components (Azizoglu et al. 2017; Shih et al. 2016), ECM and cadherins (Shih et al. 2016; Nyeng et al. 2019; Marty-Santos and Cleaver 2016) suggest that cell shape changes and cell rearrangements are driving pancreatic branching morphogenesis, but no clear inductive mechanisms for the localization of a new branch-site have been demonstrated. Our finding that a new epithelial branch is always preceded by a lumen, combined with the recent findings of Darrigrand et al (Darrigrand et al. 2024) that epithelial bifurcation is always associated with ductal cell rearrangements, suggests that ductal apical polarity could be an inductive signal for branching. Although our data also indicates that lumen formation is not sufficient for branching, suggesting that additional input, perhaps from the ECM, is required.

Several parsimonious mathematical models of branching morphogenesis have been proposed and compared to fixed tissue branching statistics (Hannezo et al. 2017; Dahl-Jensen et al. 2018; Metzger et al. 2008; Sznurkowska et al. 2018). Whilst it is fascinating that a few simple rules can govern the generation of some developmental patterns, we would argue that live imaging data such as ours demonstrate that the formation of complex patterns during organogenesis require more complex models. It would be interesting to test whether models incorporating features identified in our study better replicate pancreatic morphogenesis.

Our findings highlight the importance of utilizing live imaging to define dynamic processes. Similar studies are beginning to uncover the underlying mechanisms driving topological change during the growth and development of other organs (Varner and Nelson 2014). Whether the specific dynamic events we report in the pancreas, such as lumen transformation and loop closing, can be found in other branched organs deserves more detailed studies in the future.

## Materials and Methods

Key reagents and equipment can be found in the supplementary tables.

### Generating Muc1-mCherry reporter line

The Muc1-mCherry fusion protein construct was created by BAC recombineering using galK selection (Warming et al. 2005). To maintain glycosylation at the N-terminal, the *Muc1* gene was targeted in frame after exon 7 in order to create a fusion protein with mCherry at the intracellular c-terminal. Upon translation, this apically targeted fusion protein absorbs light in the range of 540-590 nm and emits light in the range of 550-650 nm (Shu et al. 2006).

The 171 kb BAC RP24-365K20 spanning the *muc1* gene was ordered from BACPAC Resources Center at Oakland Research Institute). After verification of the BAC by PCR for *Muc1* exon1 and 7, it was transformed into DH10B derived SW102 bacterial cells (National Cancer Institute Biological Resources Branch). These bacteria contain a fully functional gal operon, except the *galK* gene is missing. The first targeting step introduced a *galK* construct flanked by 50bp homology arms. After selection on minimal medium with galactose, a second targeting step replaced the *galK* cassette with an amplified pmCherry-N1 plasmid (kind gift Roger Tsien) PCR fragment flanked by the same homology arms used in the first selection step and with or without a Gly-Gly-Ser linker. This is achieved by negative selection on minimal medium containing 2-deoxy-galactase (DOG) with glycerol as the sole carbon source. Amplification primer sequences with homology arms are included in STable5. 6 out of 50 recombinant clones were verified with PCR of insertion sites and sequenced. 5 of these were correct. The targeted BAC (with and without linker) was purified (using the BAC100 kit from Nucleobond), but not linearized before oocyte injection at 1 ng/µl in NFR x C57bl/6 (Lund University transgenic core) or C57Bl/6N (University of Copenhagen core facility for transgenic mice) oocytes. Purified BACS were verified by SpeI digest. Three rounds of oocyte injection with the entire purified targeted BAC (Two at Lund University transgenic core and one at University of Copenhagen core facility for transgenic mice) gave rise to five founders. Only the BAC with targeted construct without the linker sequence gave rise to positive offspring.

One female founder was infertile. The four remaining founders were named Muc1-mCherry 0M, 5M, 12M (male founders) and 2F (female founder). They were all maintained on a C57BL/6BomTac genetic background. Transgenic offspring from the 0M lineage were smaller at birth and born at a 25% sub-Mendelian ratio. The other lineages were viable, fertile, phenotypically normal and born at expected Mendelian ratios. At 8-9 weeks of age the weight of most transgenic mice was not significantly different from their non-transgenic littermates. A few exceptions were females from the 0M and 5M sub-strains and males from the 2F sub-strain that had a slightly reduced weight compared to non-transgenic littermates (SFig1 A, C, E, G). In this study we henceforth used the 12M sub-strain, unless otherwise denoted in the figures.

### Animal studies

All mice were housed at the Department for Experimental Medicine at Copenhagen University and their use in experiments approved by “Miljøog fødevareministeriet – Dyrforsøgstilsynet”. Heterozygous Muc1-mCherry reporter mice of breeding age were mated with C57BL6 mice or ZO1_GFP mice (Foote, Sumigray, and Lechler 2013) to generate embryos for static and live analyses. Breeding males (12 weeks-2 years of age) were single housed, while females (8 weeks-20 weeks of age) were group housed. For timed mating, 1-2 females were added to the male cage in the afternoon and presence of a vaginal plug was checked every morning for up to 3 days. Littermates were used as controls in all experiments. Embryonic day 0.5 (E0.5) was designated as noon on the day a vaginal plug was detected. Somite pair numbers were used to accurately stage embryos between the ages of E9.0-11.0. After this stage, anatomical features such as limb and craniofacial morphology were used to verify the developmental stage. At least three embryos were analyzed per experiment.

### In vivo studies

Male and female embryos at gestational age E8.5-E18.5 were used for experiments as indicated in the results section and below. Organs were dissected from embryos at noon on the indicated day and immediately fixed in 4% paraformaldehyde (PFA) at room temperature (RT) for 30 minutes to 16 hours depending on size of the tissue and either transferred through an ascending gradient to 100% methanol for storage at −20°C or used immediately for immunofluorescence staining. For a more detailed description of embryonic mouse pancreas dissection and processing see (Heilmann, Semb, and Nyeng 2021).

### Genotyping

Tissue from one limb bud (older embryos), yolk sacs (young embryos) or ear biopsies (3-week-old pups) was used for PCR based genotyping using the REDExtract-N-Amp kit (Sigma-Aldrich) with the primers listed in STable5. PCRs were performed using the following programs, and analyzed by gel-electrophoresis:

Muc1-mCherry:

1. 94°C x 3 minutes, 2) 94°C x 45 seconds, 3) 61°C x 1 minutes, 4) 72°C x 1.30 minutes, 5) GO TO STEP 2 AND REPEAT 34 X, 6) 72°C x 8 minutes

ZO1-GFP:

1. 94°C x 3 minutes, 2) 94°C x 30 seconds, 3) 61°C x 1 minutes, 4) 72°C x 1 minute, 5) GO TO STEP 2 AND REPEAT 34 X, 6) 72°C x 2 minutes

### Fasting blood glucose and weight

Muc1-mCherry 0M, 5M, 12M and 2F heterozygous males were crossed to C57bl/6 females and the offspring was genotyped. Muc1-mCherry heterozygous and negative pups were kept until 8 weeks of age, when they were fasted overnight. The next morning, the mice were weighed, and a blood drop was collected from the tip of the tail and blood glucose was measured directly using a glucose meter. 2-4 litters were analyzed per sub-strain.

### *Ex vivo* explant cultures

35mm dishes or 8-well µ-Slides (Ibidi) with #1.5 coverglass bottoms were used for culturing and imaging ex *vivo* whole embryonic pancreata and lung. Silicon inserts (Ibidi) were applied to 35mm dishes to ensure explants could be identified for genotyping after imaging and for increasing the explant attachment rate. All culture vessels were pre-coated for 45 minutes at RT with 100 µg/ul of Fibronectin (Invitrogen) to allow explant attachment, after which explant media (Media 199 ^with^ phenol red supplemented with 10%FBS, 50 U/mL of penicillin and Streptomycin and 0,25 µg/ml of Fungizone) was added to culture vessels. Explants were cultured at 37°C at 5% CO_2_ for 6 to 48 hours before imaging (see below for experiment-specific pre-imaging and imaging culture times). Shortly before imaging, the explant medium was replaced with imaging medium (Media 199 without phenol red supplemented with 10%FBS, 50 U/mL of penicillin and Streptomycin and 0,25 µg/ml of Fungizone).

### Pre-midgestation embryos (E9.0-E11.0)

Embryos were collected from their mother’s dam and the posterior half of their gut tubes or lung buds were isolated by fine dissection under a stereoscope in sterile PBS. The lung buds were directly transferred to a coated culture vessel for evaluation of Muc1-mCherry expression by live imaging. To determine which mechanism is responsible for establishing the pancreatic CPL we established a protocol for live imaging of whole E9.75 to E10.0 posterior foregut explants. This method includes dissecting out and explanting the dorsal and ventral pancreatic buds, the biliary bud, foregut, and some of the prospective stomach. Keeping so much of the posterior foregut intact aids identification of the early anlage, increases explant survival and attachment and maintains tissue physiology. To minimize the thickness of tissue to be visualized and to aid explant attachment, epithelium and a thin layer of mesenchyme was removed from the ventral side of the gut tube.

### Midgestation embryos (E11.5-E12.5)

E12.5 Embryos were collected from their mothers and the dorsal pancreas was isolated by fine dissection under a stereoscope in sterile PBS and explanted onto fibronectin coated culture vessels. The medium was changed to fresh explant media the following day. Explants were cultured for a total of 48 hs before live imaging or fixation. For the ECM expression studies, dorsal pancreas was explanted from E11.5 embryos and cultured for 3 days before fixation and immunofluorescence staining.

### Immunofluorescence staining of cryo-sectioned tissue

Embryonic tissues were processed for cryo-sectioning after fixation in 4% PFA and washing in PBS. Samples were incubated overnight in 30% sucrose diluted in 1x PBS, equilibrated through 30% sucrose:OCT 1:1 for one hour into 100% OCT for one hour and finally transferred into cryo-molds filled with fresh OCT for freezing on blocks of dry ice. Cryo-sectioning at 6μm was performed using a Leica cryostat.

For immunofluorescence staining, tissue was first cleared of OCT and permeabilized by washing three times for 5 minutes in PBST (1x PBS with 0.5% or 0,01% Triton-X) and then incubated in a blocking solution of 5% neonatal donkey serum (NDS) in PBST for a minimum of 1 hour. Primary antibodies were diluted in a blocking solution and applied to the sections and incubated at 4°C overnight (16 hs) in a humidified chamber. For a list of antibodies and dilutions see STable 2.

Subsequently, the primary antibodies were removed, and the slides washed in PBST three times before application of diluted fluorophore-conjugated secondary antibodies, (STable 3), in a humidified chamber for 1 hour. The secondary antibody solution was removed, and the slides washed in PBST. Slides were mounted in DAKO mounting solution with or without Hoechst 33342 (Invitrogen, H3570/23363w, 1:1000) using #1.5 cover glasses.

### Whole mount immunofluorescence staining (WMIF)

Embryos were collected from their mother’s dam and the posterior half of their gut tubes (pre-midgestation embryos) or the stomach, pancreas and duodenum (mid-late gestation embryos) were isolated by fine dissection in sterile PBS. 4% PFA was used for fixation of all samples; pre-midgestation gut tubes were fixed 30 mins −1 hour at RT, mid-late gestation organs were fixed overnight at 4°C. After fixation, samples were washed in PBS and dehydrated through methanol:PBST gradations into 100% MeOH. mid-late gestation samples were bleached in a 1:1:4 dilution of 30%H2O2: DMSO:MeOH (Dent’s Bleach) for up to 2 hours at RT. Samples were then rehydrated through methanol:PBT2 gradations into PBT2 and blocked in 5%NDS;0.5%PBST or blocking buffer for 2 hours at RT, or for 16-24 hours at 4°C. Primary antibody was then applied (see STable 2) and samples left to incubate, shaking, for 16-24 hours at 4°C. The following day primary antibody and samples were washed in 0.5%PBST once an hour for 5–8 hours at RT to remove any unbound antibody from the tissue. Subsequently, secondary antibody in 0.1%PBST was applied to samples and incubated overnight at 4°C (see STable 3). The following day, unbound secondary antibody was removed by washing samples in 0.1%PBST once an hour for 5–8 hours at RT. To prepare for clearing, the tissues were dehydrated in methanol: 0.1%PBST gradations to 100% MeOH. Finally, samples were transferred to a clearing solution i.e. BABB (1:2 benzyl alcohol and benzyl benzoate) for late gestation samples, methyl salicylate for pre-midgestation gut tubes and explants. For immunofluorescence staining of *ex vivo* cultured explants, medium was removed, and explants fixed in the culture vessel for 30 mins in 4% PFA at RT. After fixation, explants were washed in PBST and incubated with blocking buffer overnight at 4°C. Explants were then incubated with primary antibodies (see STable 2) in blocking buffer over 3 nights at 4°C for optimal antibody penetration. Subsequently, explants were washed with PBST for 5-6 hours, followed by secondary antibody (see STable 3) incubation overnight at 4°C. Explants were directly imaged in PBS after washing. No clearing agent was used as it degrades the plastic surrounding the glass bottom surface.

### Image acquisition

A subset of images in figure 1 was recorded on a Zeiss axioplan2 widefield fluorescence microscope. All other image acquisition was conducted with inverted laser scanning confocal microscopes. A Leica SP8 system with 20x and 63x glycerol immersion objectives was used for a subset of fixed imaging. Zeiss LSM780 or LSM880 systems with 40x water immersion objectives were used for live imaging, and 40x and 63x oil immersion objectives for fixed imaging. See specifications of systems and objectives in STable 7.

### Live imaging

STable 1 contains individual imaging settings for each live imaging experiment.

Before imaging, explant media was removed from the sample dish and pre-heated imaging media added. The on-stage incubation chambers were pre-heated to 37°C with 5% CO2 supply for a minimum of 2 hours prior to image acquisition. Image acquisition was performed using Zeiss LSM780 or LSM880 Airyscan (in regular scanning confocal mode) confocal systems. Multi-position images were acquired at an interval of either 4, 8, 10 or 12 minutes (as indicated in figure legends and STable 1). Pixel resolution was as standard at 512 × 512 for pre-midgestation embryos and 1024 × 1024 for midgestation embryos unless stated otherwise. Laser lines were kept to a minimum in all imaging sessions to reduce phototoxicity and DIC light was collected with the PMT detector. The imaging session was carried out for a minimum of 20 hours and a maximum of 72 hours for one dish of samples. For a subset of imaging sessions, the explants were fixed and stained post-imaging by immunofluorescence staining to assess epithelial integrity, lumenal network morphology and, where relevant, cell differentiation. Assessments were made within the same dish for explants that had been exposed to the confocal laser and those that had not (i.e. were not imaged). For experiments where enough embryonic material was available a non-imaged control set of explants were also kept in a standard incubator and used as a second control for any effect of the microscope on-stage incubator.

### Image processing, rendering and analysis

Image processing, rendering and analysis was performed using Imaris, Fiji/ImageJ or Matlab. All confocal images are shown as either 2D single Z-slices or 3D maximum intensity projections (MIP). Image intensity and contrast has been adjusted across the entire image/stack to aid in visualization. Any further processing is mentioned in the legends or below for each experiment.

### Processing and rendering of live imaging movies of pre-midgestation (E9.5-E10.5) pancreata

For analysis of lumenal network morphogenesis the following pre-processing steps were applied in Imaris:

1. Files were split into single positions
2. A median (3×3×3) smoothing filter was applied to ZO1-GFP and Muc1-mCherry channels.
3. Where necessary to improve visualization Z-slices at either end were deleted from the image stack to remove non-epithelial signal (e.g. ZO1-GFP in endothelial cells, see S Figure 5 and main text for explanation of how to identify this signal) or to reduce noise. Raw data files are provided for comparison. No data was removed during quantifications.
4. Because of the unstereotypical/heterogeneous expression of lumen and foci marked by the live imaging reporters, Imaris could not faithfully segment all lumenal structures automatically. Hence all tracking was undertaken manually. For lumenal network tracking the Imaris spot function was used as a rendering method (see below).

### Pre-midgestation lumenal network tracking and quantification in Imaris

1. After rendering (see above) movies were cropped in time to a point where most lumenal branches could be tracked backward in time i.e. out of frame data was deleted.
2. Each unique lumenal branch during the imaging interval was defined in a new folder. The Imaris spot function was used in each 4-minute time frame to mark - independently - whether a branch was contributed to by: i) extension of an existing lumen, ii) a temporary or iii) permanent break point, or, iv) a derived or v) de novo foci. We also observed lumenal merging events but did not track these. Events were tracked back and forth in time to ensure the tracking was accurate.
3. To simplify the tracking, we only tracked events that contributed to permanent proximal-distal growth of a lumen. In other words, foci or breaking events that may contribute to media-lateral growth of the lumen, or to temporary structures such as lumenal spikes, were not quantified. This means that the number of total events were underrepresented in this dataset.
4. When a foci was tracked, a measurement was also taken of the distance of this foci to surrounding lumenal structures. We presume that foci less than the average epithelial cell length away (<10 µm) are foci derived from already polarized MPPs and those more than a cell length away (>10 µm) could be generated ‘de novo’ from unpolarized MPPs. We defined an average cell length from measurements of elongated and rounded pancreatic cells in WMIF stainings, (quantified in graph SFig5F from E10.0-10.5 WMIF data shown in SFig 5D&EF).
5. At the end of the Imaris tracking we exported the tracking count data in excel table and input the data into Prism for analysis and visualization as scatter graphs.

### Processing and rendering of fixed in vivo pre-mid-gestation (E9.5-E10.5) pancreata

All fixed data was processed by applying the median (3×3×3) smoothing filter in Imaris, any channels lacking signal were also deleted. In some samples there was a strong autofluorescence, derived from red blood cells covering the pancreatic bud, or from debris in mesenchyme. In these cases, to be able to view the lumenal signal in the 3D MIP we segmented the CDH1 or PDX1 channel. Once we had a faithful pancreatic epithelium segmentation, we could view the lumenal channel within the segmentation as a unique ‘masked lumenal channel’. For an example see SFig4Gi.

### Processing, rendering and analysis of live imaging movies of mid-late gestation (E14.5-E16.5) pancreata

#### Movie preprocessing

##### Noise filtering

First we performed 3D gauss filtering (sigma=0.5) then 3D median filtering (3×3×3 neighborhood) of original Muc1 channel images, ORG_IM, producing IM.

##### Filtering out small bright artifact vesicles removed from lumenal network

The Muc1-mCherry channel contains small and very bright spots that are most likely Muc1-mCherry concentrated in intracellular vesicles. We removed these by constructing a REMOVE_MASK, as they interfere with the analysis of ducts.

1. We determine the 99-intensity percentile threshold, T_99, using as input 10 consecutive timepoints at a time. We use T_99 to threshold IM and obtain a mask (MASK1) containing only the very brightest objects (bright vesicles inside epithelial cells and lumenal tips). We then produce a second mask, MASK2 by taking the 97-percentile threshold (T_97) of IM. From MASK2 we keep only connected components larger than 3000 voxels, by using MASK2=bwareaopen(MASK2,3000). Our REMOVE_MASK is then all connected components (cc’s) in MASK1 that do not have an overlap with connected components in MASK2.
2. We then produce a third mask, MASK3 by taking the 95-percentile threshold (T_95) of IM. We produce MASK4 by keeping only cc’s larger than 3000 voxels from MASK3 and dilating the remaining component with a spherical structuring element of radius 1 pixel. REMOVE_MASK is then set to include objects in MASK3 which do not have an overlap with objects in MASK4.
3. We dilate the REMOVE_MASK with a spherical structuring element of radius 1 pixel. Intensity values or IM inside REMOVE_MASK are replaced with those of voxels from 3D medfilt(ORG_IM) (15×15×3 neighborhood), producing the ‘cleaned/filtered’ IM_CLEAN result. See examples of raw image and filtered image in SFig11A-B

##### Loop tracking

Maximum intensity projections (MIPs) of the prepossessed/cleaned IM_CLEAN (see steps above) Muc1-mCherry channel movies were produced. Using a custom designed GUI we tracked/detected all loop structures and detected lumen breaking events in 2D. When a loop cinches in two, we record the largest as a continuation of the old loop, and the smallest as the new loop. The human annotator determined beginning and end times for a given loop event and clicked on the approximate x,y coordinates of the loop center at every 5 time points, (the loop position in the 4 unannotated timepoints between two annotated timepoints were then determined by linear interpolation). All such manually detected loop events were then extracted from the data as 4D (3D plus time) cropped movies and inspected visually in Imaris to confirm that they were indeed bonafide loop structures when examined in 3D, and not just lumen structures crossing each other in different image planes. The dark area (without Muc1-mCherry signal) inside each loop was then segmented in MIP 2D movies in a semi-automated fashion with final hand corrections done by a human in itk-snap, when needed. The rest of the lumenal network (i.e. areas outside the cropped/extracted loop movies) was segmented in a fully automated manner.

### Processing, rendering and analysis of fixed *in vivo* pancreas and fixed *ex vivo* pancreas from mid gestation (E14.5)

#### Image preprocessing

##### Intensity rescaling

Original 2 color images (Ecad and Muc1) were stored as 16 bit but in reality less than 14 bit was used (Zeiss czi files have a true bit depth of 14). The images all had their intensities rescaled to fit 8 bit by recasting to double type, normalizing/dividing with 99.9 percentile intensity value of the respective channel multiplying with 255 and recasting to uint8.

##### Spatial rescaling

All 3D images where rescaled spatially to become isotropic with xyz resolution [1 micron x 1 micron x 1 micron] per pixel, interpolation to new grid was bicubic, (the output pixel value is a weighted average of pixels in the nearest 4-by-4 neighborhood). We used the MATLAB imresize() function.

#### Semantic segmentation

##### Random forest prediction

A random forest (RF) model was trained using ilastik (https://www.ilastik.org/), based on sparse annotations performed by hand on select images in itk-snap. In ilastik, all possible image features were selected at all possible spatial scales. Using the trained ilastik RF model, pixels/voxels of all images were segmented into four classes: 1) lumen, 2) epithelial cell 3) mesenchymal cell and 4) medium/glass/nothing, with simple segmentation output.

#### Post prediction processing

##### Morphological operations performed on RF output

*Step by step operations performed on pixel class 1 / Lumen:*

● Close of small ‘holes’ in *Lumen* by performing morphological closing with a spherical structure element with radius=1. MATLAB function imclose(*Lumen*,strel(’sphere’,radius)).
● Fill holes in *Lumen*. MATLAB function imfill(Lumen,’holes’). Based on 6-connectivity.
● Remove isolated pixels, (isolated based on 26-connectivity), with MATLAB function bwmorph3(Lumen,’clean’).
● Fill all holes in *Lumen* again.

○ Make a mask, *LumenConnMask*, which includes the entire lumenal network by performing morphological closing with a cubic structure element with radius=20. MATLAB function imclose(im,strel(’cube’,radius)).
○ From *LumenConnMask*, keep only connected objects larger than 1e4 pixels.
● Remove pixels in *Lumen* that are outside *LumenConnMask.* This gets rid of small objects further away than 20 pixels from the main lumenal network

Label different connected components in *Lumen* (connectivity 26) with different indices and inspect visually in itk-snap. Join up disjoined elements if needed by hand (sometimes classification fails for very small/thin lumens which can cause the lumenal network to break).

*Step by step operations performed on pixel class 2 + Lumen (from above):*

● Join pixel class 2 with *Lumen* as *Epi* (epithelium mask). (MATLAB: Epi = P==2 | Lumen);.
● Fill holes in *Epi*.
● Remove isolated pixels.
● Keep a pixel set to 1 if 14 or more voxels (the majority) in its 26-connected neighborhood are 1; otherwise, set the pixel to 0. with MATLAB function bwmorph3(Epi,’majority’).

○ Make a mask, *EpiConnMask*, which includes the entire epithelium by performing morphological closing with a cubic structure element with radius=15.
○ From *EpiConnMask*, keep only connected objects larger than 2e6 pixels.
Remove pixels in *Epi* that are outside *EpiConnMask.* This gets rid of small objects further away than 15 pixels from the main epithelium.

*Step by step operations performed on pixel class 3 / Mesenchyme:*

● Small ‘holes’ in class 3/mesenchyme (*Mesen*) were closed by performing morphological closing with a spherical structure element with radius=4.

Lumen overrules Epi which overrules Mesen in terms of pixel identity.

#### Network feature extraction

##### lumenal/epithelial graph extraction and loop count/detection

From Lumen and Epi we made a 1 pixel width skeleton representation (LumenSkel and EpiSkel) of the lumenal network using the MATLAB bwskel() function. E.g. bwskel(Lumen,’MinBranchLength’,15) (which uses the medial axis transform). We then made the skeleton images into a MATLAB graph object where each pixel=1 in the skeleton image was considered a node and edges were introduced between nodes closer than √3 pixels apart, (the distance of the internal diagonal of a 1×1×1 cube). Forming a graph in this way has the unfortunate side effect of forming some small ‘false’ loops/cycles (with 3-6 nodes) near branch points. Using a custom written function, we removed select edges in order to remove/break such loops of size less than 15 nodes. Using the MATLAB function cyclebasis() we found all loops/cycles in the graph. The cyclebasis() function returns a cycle basis (a set of cycles with which one can construct all possible cycle paths in the graph) for the graph but not necessarily the basis with minimal overlap between node sets of each cycle. Using a custom written function, we rearranged cycles to get the basis with the least amount of overlap possible between node sets in each cycle.

##### Locating endpoints, branch points and mid-length points in skeleton images

Using the MATLAB function bwmorph3(skelIM,’endpoints’) and bwmorph3(skelIM,’branchpoints’) we located endpoints and branch points in the lumenal network. We found the points on the skeleton at mid length between branch points in two steps: First we determined the distance transform (along the skeleton) to the nearest branch point with D=bwdistgeodesic(skelIM,branchPointIM). We then found the ‘mid length points’ by locating the regional maxima in D with imregionalmin(D).

##### Determining lumen and epithelial width

We used the MATLAB distance transform function (bwdist()) to make Dlum=bwdist(imcomplement(Lumen)) and Depi=bwdist(imcomplement(Epi)), the approximate short axis x 1/2 of the lumen/epithelium (in microns) could then be found by extracting the pixel value of distance image D along the skeleton image pixels.

### Processing and rendering of images from fixed *in vivo* late gestation (E14.5) pancreata from Muc1-mCherry embryos

For analysis of any phenotype of the Muc1-mCherry reporter mouse, whole dorsal pancreata, duodenum and stomach from reporter positive and negative littermates were stained for MUC1, INS and CDH1 using the WMI protocol, and imaged in 3D using the Sp8 confocal. Segmentation of all channels was done using Imaris Surfaces wizard with identical settings for all embryos. Volume (µm^3^) CHD1, INS and MUC1 was exported to Excel, where volumes of INS and Muc1 were normalized by dividing with CHD1 volume.

### Data analysis

### Statistical analysis

All statistical tests were based on at least n=3, as indicated in figure legends. Before statistical testing, data was checked for normality and equal variance. Statistical analysis was performed in GraphPad Prism or Matlab.

In Graphpad Prism, data following a Gaussian distribution was analyzed by unpaired t test when F test for equal variance showed no statistical difference (p>0,05) and otherwise with unpaired t test with Welch’s correction.

For the count data pertaining to the tracking of early lumenal events contributing to secondary network formation, we performed a Chi-Squared test in GraphPad Prism to check that the observed counts were not likely to have occurred by chance.

In MATLAB normality was checked via the Anderson-Darling test using the function adtest(), equal variance was checked using vartest2(). The 2-sample t-test was performed using the function ttest2(). For distributions that did not pass the normality test, we used the Wilcoxon rank sum test (using the function ranksum()) to test for unequal medians. Rank correlation (Spearmans’ Rho) between variables X and Y was determined using the function corr(X,Y,’type’,’Spearman’).

### Models

#### Design of the ‘combined size feature’

We engineered a new combined size feature that is simply a linear combination of different features quantifying loop size. This was done for several reasons: 1) to reduce dimensionality of the feature space in a way where we retain information relevant to the prediction of the two target/response variables, 2) to tackle the problem of collinearity between these features and 3) to arrive at a combined size feature which is more robust to measurement error/noise in the individual features. This new combined size feature was used when dividing the loop data into our size categories S, M, L and XL, see S Fig10A. The original size features contributing to the combined feature were:

1. *’inner hole diameter’,*
2. *’convex area’,*
3. *’perimeter of convex hull’,*
4. *’minor axis length’,*
5. *’major axis length’*
6. *’lumen loop and hole total area’*.

(These were log transformed if needed and z scored). These original size features where weighted in the new combined size feature according to their correlation with the two target variables ‘Time Left’ (hereafter ‘TL’ - time left before loop closes) and ‘loop contraction rate’ (hereafter ‘dP/dt’ - signed contraction speed of loop perimeter per unit length). I.e. the feature weights are proportional to the sum of Spearmans rho with respect to rho_TL and rho_dPdt. We checked that our new combined size feature is as correlated with TL and dPdt as any of the individual size features are.

#### Linear model specifications

We used the ‘fitlm()’ MATLAB function when fitting the linear regression models, which contain an intercept (constant) term and one linear term per predictor variable, (*1. ‘combined size feature’, 2. ‘number of outgoing lumenal network connections’, 3. ‘mean distance to the epithelial mesenchymal border’, 4. ‘mean lumen width’, 5. ‘loop eccentricity’)*.

#### Neural network model specifications

We used the ‘fitrnet()’ MATLAB function for fitting our regression neural networks.

*Hyperparameters:*

Input layer = 5 features, (the features are: *1. ‘combined size feature’, 2. ‘number of outgoing lumenal network connections’, 3. ‘mean distance to the epithelial mesenchymal border’, 4. ‘mean lumen width’, 5. ‘loop eccentricity’*.

Number of hidden layers = 2. The first layer has 100 neurons and the second 10, first layer is followed by a ReLu activation layer. The final output layer is one neuron (with identity function). An explorative hyperparameters search (varying width and depth of the NN) was performed showing that other models with similar total numbers of parameters gave similar prediction results.

Loss function: Mean squared error, (MSE).

Solver/Optimizer: limited-memory Broyden-Fletcher-Goldfarb-Shanno quasi-Newton algorithm (LBFGS). Mini batch size = 1.

Training was stopped if the validation loss was smaller than the minimum validation loss recorded so far, for more than 6 iterations in a row, (since this indicates that further training is overfitting the model on the training data and causing an increase in validation loss).

#### Base models

The base models are very simple models that just return the median of time left in training data for one model and the median of contraction/expansion speed in the training data for the other model. The utilization of such basic models, (that merely returns the median value of the target variable), serves as a benchmark, representing the minimum predictive capability expected from more sophisticated modeling approaches.

#### Division of data into training and validation

One loop time course consists of all the images of the same loop tracked over time. We divided the time course dataset in two: training and validation. (Training - 106 loop time courses, 7515 images, Validation 66 loop time courses, 4146 images). Note that for some of the time courses (49 out of 172) we do not observe loop closure (either because the loop moves out of frame or because the movie ends before loop closure or in rare cases because the lumen breaks). This means that the dataset used for predicting ‘Time left’ is smaller than the one used for predicting ‘Contraction/expansion speed’. 76 time courses for training (4651 images) and 47 time courses for validation (2315 images).

The training/validation division of the data set was done by finding closets pairs of loop time courses via k nearest neighbor search in the space spanned by the time course features ‘*total time course duration’, ‘average number of lumen branch points on loop’, ‘average distance of loop to border of explant’, ‘average loop size’* and *’average loop long axis’*. Each member of such a *’closest pair’* was then put in the training set and in the validation set respectively. The remaining time courses were all put in the training dataset. This pairing and division of loop time courses was done to make sure that training and validation sets contained relatively similar types of data. Note that we never want training and validation sets to contain data/images from the same time course, since the similarities that a loop image has to itself taken at an earlier/later time point causes more complex models to overfit.

### Tracking plot scripts for plot generation

The tracking plots (*Fig 3E and SFig 6F-G*) were generated using custom scripts made in R (v 4.2.2) and several R packages (see STable6). The code and datasets used to generate the plots have been deposited in Zenodo and can be accessed using the following link https://doi.org/10.5281/zenodo.8248940. Continuous lumen growth over time is displayed as a solid horizontal line along the X-axis of the plots. Bifurcation points resulting in another tracked lumenal branch are plotted as a perpendicular solid line from the mother lumenal branch. Dashed horizontal lines plotted above the mother lumenal branch represent the frequency and persistence of BPTs. Derived (circles) and de novo foci (squares) are represented at the time frame in which they appear to contribute to the lumen.

## Supplementary Tables

**STable 1:** Live imaging acquisition specifications.

| Figure (Fig) or Movie (M) | Description | Imaging system | Imaging type | Time steps (min) | Laser Lines (%) | Gain | Pinhole | X,Y,Z (μm) |
| --- | --- | --- | --- | --- | --- | --- | --- | --- |
| Fig2A/ M1 | Explanted E9.5 posterior foregut expressing ZO1-GFP visualizing the evaginating pancreatic buds. | Zeiss LSM780 | Z-stack, sequential acquisition | 8 | 488:1%<br>561: 1.5%<br>(PMT for DIC acquisition in the 488 track) |  | GFP: 2.01AU<br>mCherry : 2.40AU | 0,69 x 0,69 x 1 μm |
| Fig2C-D and Fig3A-D/ M4, M6, and M8 | Explanted E10.5 posterior foregut expressing Muc1-mCherry:ZO1-GFP | Zeiss LSM780 | Z-stack of 40 μm, sequential acquisition (unidirectional scan) | 8 | 488:1%<br>561: 1.5%<br>(PMT for DIC acquisition in the 488 track) |  | GFP: 2.01AU<br>mCherry : 2.40AU | 0,69 x 0,69 x 1 μm |
| Fig3Gi-iii and M7 | Explanted E10.5 posterior foregut expressing Muc1-mCherry & ZO1-GFP recorded to track the frequency of each event during secondary network transformation. | Zeiss LSM880 | Z-stack of 40 μm, sequential acquisition | 4 | 488:1%<br>561: 1.5%<br>(PMT for DIC acquisition in the 488 track) | GFP: 1150<br>mCherry: 1000<br>T-PMT: 300 | GFP: 1.88AU<br>mCherry :2.25AU | 0,69 x 0,69 x 1 μm |
| SFig3A-C/ M2 | Explanted E10.5 posterior foregut expressing Muc1- | Zeiss LSM780 | Z-stack of 50μm, sequential | 8 | 488:1%<br>561: 1.5%<br>(PMT for | GFP: 1150<br>mCher | GFP: 1.88AU<br>mCherry | 0,69 x 0,69 x 1 μm |
|  | mCherry1-mCherry & ZO1-GFP demonstrating acquisition of Mucin in more mature lumens already marked by ZO1. |  | acquisition |  | DIC acquisition in the 488 track) | ry: 1000 T-PMT: 300 | :2.25AU |  |
| Fig5B-D, SFig9, Fig7E, SFig7, SFig9E, SFig 11 / M9-15 | Muc1-mCherry mid gestation live imaging | Zeiss LSM780 | Time-lapse, multi position, Z-stack, Simultaneous acquisition | 10 | 488:1% 561: 1,5% | GFP: 973 mCherry: 1009 T-PMT: 250 | 1,73AU | 0,35 x 0,35 x 1 µm |

**STable 2:** Primary Antibodies.

| Name | Host | Dilution | Source | Identifier |
| --- | --- | --- | --- | --- |
| Ezrin [3C12] | mouse | 1:1000 | Abcam | ab4069<br>RRID:AB_304261 |
| CHD1 | Rat | 1:100 | Takara Bio | M108<br>RRID:AB_2895157 |
| Fibronectin | Rabbit | 1:200 | Abcam | ab2413<br>RRID:AB_2262874 |
| GFP | Chicken | 1:500 | Abcam | ab13970<br>RRID:AB_300798 |
| GFP | Goat | 1:500 | Abcam | ab5450<br>RRID:AB_304897 |
| Glucagon | Mouse | 1:1000 | Sigma/Merck | G2654<br>RRID:AB_259852 |
| Integrin $\alpha 5$ | Rabbit | 1:100 | Abcam | ab150361<br>RRID:AB_2631309 |
| INS | Guinea Pig | 1:750 | Agilent | A056401-2<br>RRID:AB_2617169 |
| Laminin | Rabbit | 1:200 | Sigma | L9393<br>RRID:AB_477163 |
| MUC1 1 | Armenian Hamster | 1:1000 | Thermo Scientific | HM-1630-P0<br>RRID:AB_54620 |
| Pdx1 | Guinea Pig | 1:1000 | Beta Cell Biology Consortium | AB2028<br>RRID:AB_10014623 |
| Pdx1 | Guinea Pig | 1:1000 | Abcam | ab47308<br>RRID:AB_777178 |
| RFP-biotin IgG | Rabbit | 1:100 | Abcam | ab34771<br>RRID:AB_777699 |

**STable 3:** Secondary Antibodies.

| Name | Host | Dilution | Source | Identifier |
| --- | --- | --- | --- | --- |
| Cy3 anti Rat IgG | Donkey | 1:1000 | Jackson<br>ImmunoResearch | 712-165-150<br>RRID:AB_2340666 |
| Alexa Fluor® 647<br>anti Armenian<br>hamster IgG | Goat | 1:1000 | Jackson<br>ImmunoResearch | 127-605-160<br>RRID: AB_2339001 |
| Alexa Fluor® 488<br>anti Rabbit FAB<br>fragment | Donkey | 1:1000 | Jackson<br>ImmunoResearch | 711-547-003<br>RRID: AB_2340620 |
| Dylight® 405 anti<br>Guinea Pig IgG | Donkey | 1:1000 | Jackson<br>ImmunoResearch | 706-475-148<br>RRID: AB_2340470 |
| Alexa Fluor® 647<br>anti Rat IgG | Donkey | 1:500 | Jackson<br>ImmunoResearch | 712-605-153<br>RRID: AB_2340694 |
| Cy3 anti Armenian<br>hamster IgG | Goat | 1:500 | Jackson<br>ImmunoResearch | 127-165-160<br>RRID: AB_2338989 |

**STable 4:** Mouse strains.

| Name | Source | Identifier |
| --- | --- | --- |
| C57BL/6BomTac | Taconic | B6JBOM-F/M |
| Zo1-GFP | Terry Lechler lab, Duke University | (Foote, Sumigray, and Lechler 2013) |
| Muc1-mCherry | This paper | N/A |

**STable 5:**
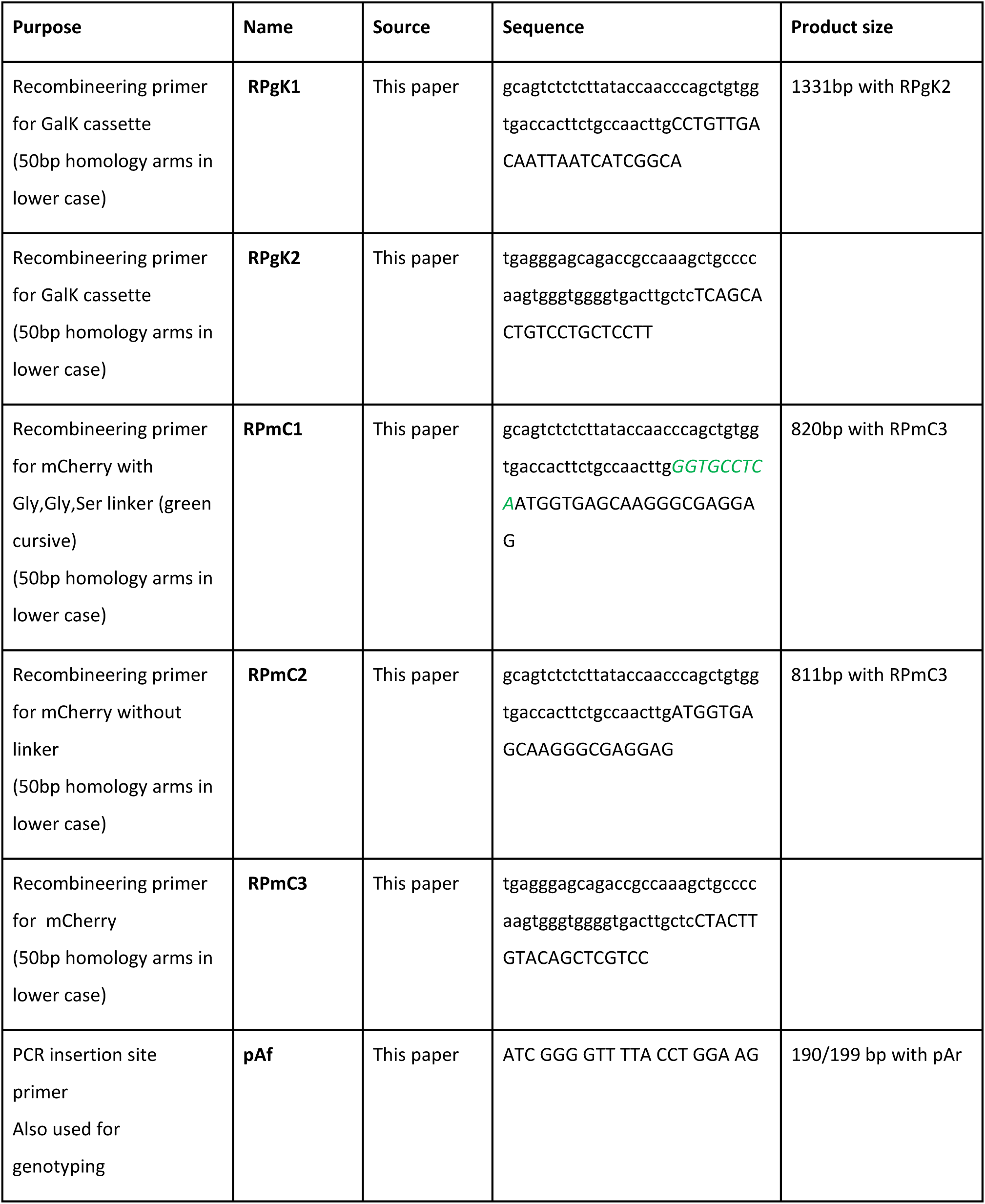
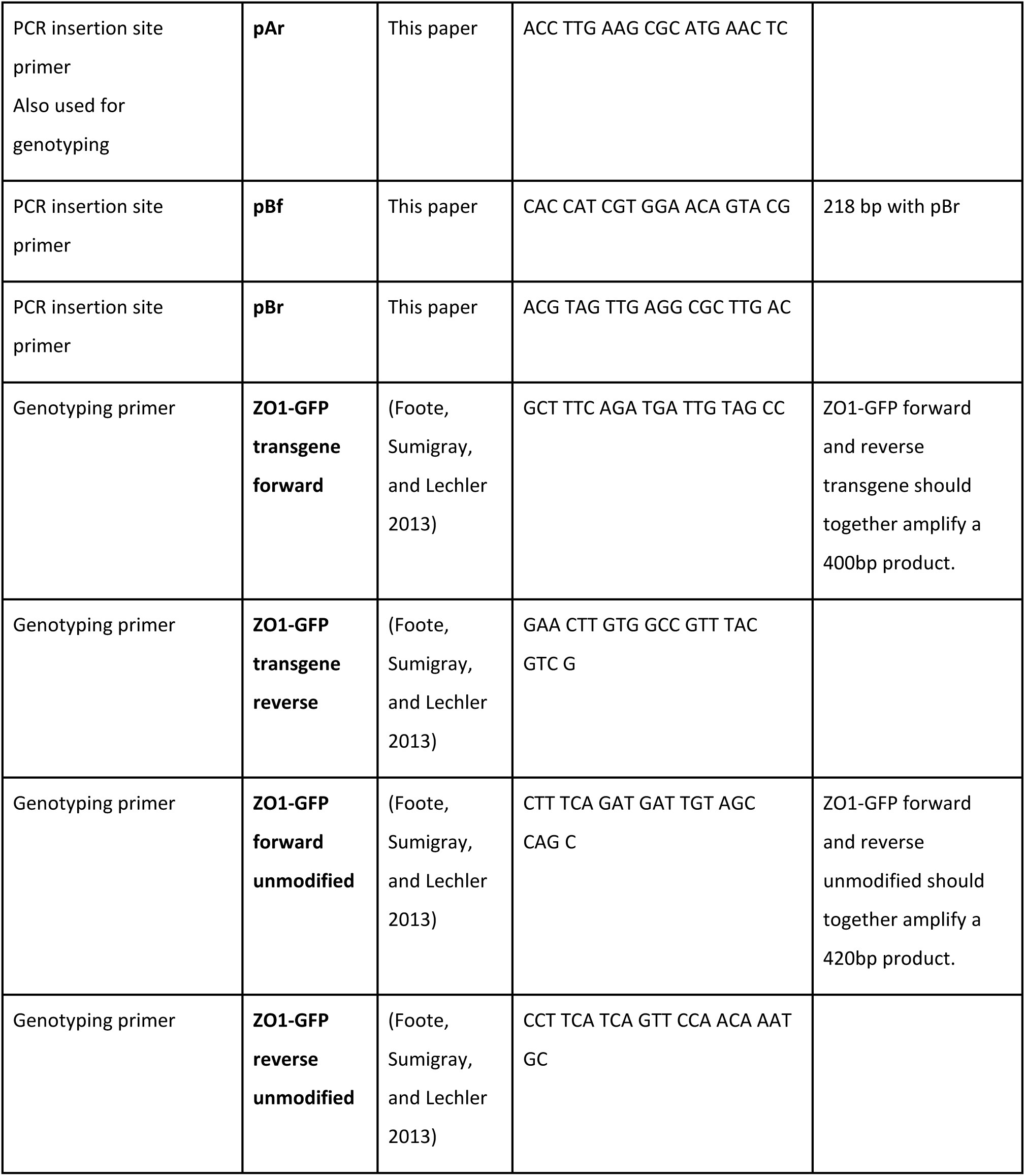
Oligonucleotides.

**STable 6:** Software and Algorithms.

| Name | Source | Webpage |
| --- | --- | --- |
| Fiji 2.0/ImageJ version 1.53t | (Rueden et al. 2017) | http://imagej.nih.gov/ij |
| Imaris Essentials version 8.4, 9.0-10.0 | Bitplane/Andor | https://imaris.oxinst.com |
| MATLAB R2022b with the Statistics and Machine learning toolbox and Computer Vision Toolbox | MathWorks | https://se.mathworks.com/products/matlab.html |
| R version 4.2.2 | (The R core Team 2021) | https://www.R-project.org |
| R packages | tidyverse (v 1.3.2)<br>(Wickham et al. 2019), plotly (v 4.10.1)<br>(Sievert 2020)<br>and GenomicRanges (v 1.50.2)<br>(Lawrence et al. 2013) | N/A |

**STable 7:** Additional key reagents, equipment and resources.

| Name | Source | Identifier |
| --- | --- | --- |
| 35 mm glass bottom dish | WillCo | GWST-3512 |
| μ-Slide 8 Well high | Ibidi | 80801 |
| TSA-Fluorescein kit | Akoya BioSciences | NEL701A001KT |
| Blocking Reagent | Akoya Biosciences | SKU FP1012<br>RRID: N/A |
| Fibronectin bovine protein | Thermo-Fisher | 33010018 |
| Media M-199 with phenol red | Gibco/Thermo Fisher | 31150022 |
| Media M-199 without phenol red | Gibco/Thermo Fisher | 11043023 |
| Zeiss LSM 780 inverted confocal | Zeiss and PECON | RRID:SCR_02092<br>2 |
| scanning microscope equipped with PECON XL incubator, heating insert, and temperature, humidification, and CO2 modules |  |  |
| Objective: C-Apochromat 40x/1.20 Water Corr | Zeiss | RRID: N/A |
| Objective: Plan-Apochromat 40x/1.30 Oil DIC | Zeiss | RRID: N/A |
| Objective: Plan-Apochromat 63x/1.40 Oil DIC | Zeiss | RRID: N/A |
| Zeiss LSM 880 with Airyscan inverted confocal scanning microscope equipped with PECON XL incubator, heating insert, and temperature, humidification, and CO2 modules | Zeiss and PECON | RRID:SCR_020925 |
| Leica SP8 inverted laser scanning confocal microscope | Leica | RRID: N/A |
| Objective: Plan-Apochromat CS2 20x/0.75 IMM CORR | Leica | RRID: N/A<br>11506343 |
| Objective: Plan-Apochromat CS2 63x1.30 Glyc CORR | Leica | RRID: N/A<br>11506353 |
| Zeiss axioplan2 widefield fluorescence microscope | Zeiss | RRID: N/A |

## Acknowledgements

We thank Jette Larsen, Anna Månsson and Diana Klüver Hansen for technical assistance. We thank Roger Tsien, Howard Hughes Medical Institute Laboratories for the pmCherry-N1 plasmid, and Terry Lechler at Duke University for the ZO1-GFP mice. We acknowledge assistance and support for computational resources and infrastructure via the Health Data Science Sandbox (https://hds-sandbox.github.io) funded by the Novo Nordisk Foundation (grant number NNF20OC0063268). We acknowledge the Danstem Imaging platform and the Core Facility for Integrated Microscopy, Faculty of Health and Medical Sciences, University of Copenhagen for use of microscopes and image analysis work stations. We acknowledge the Lund University Transgenic Core and the University of Copenhagen Core Facility for Transgenic Mice for oocyte injections. We thank and acknowledge Casper Øbro for his expertise and skills used to generate the 3D illustrations used in the schematic figures (Fig. 8) of this paper. PN and JMK were supported by the Novo Nordisk Foundation grant number NNF17OC0028360 and JARH by the Novo Nordisk grant number NNF20OC0063268. SH was supported by DFF grant number 6110-00091B. The Novo Nordisk Foundation Center for Stem Cell Biology was supported by Novo Nordisk Foundation grant number NNF17CC0027852.

## Author contributions

PN and AJ conceived the study; designed, conducted, and analyzed the experiments and wrote the manuscript. SH conceived the study, analyzed the experiments, developed the image analysis methods, performed the computational modeling, created artwork and wrote the manuscript. CE performed the ECM staining and imaging and edited the manuscript. JMK assisted with embryo dissection, explant culture, immunofluorescence staining, and image analysis. JARH developed the tracking graph plots (computational script and aesthetics). HS conceived the study, acquired funding and edited the manuscript.

## Supplementary Figure Legends

**SFigure 1:**
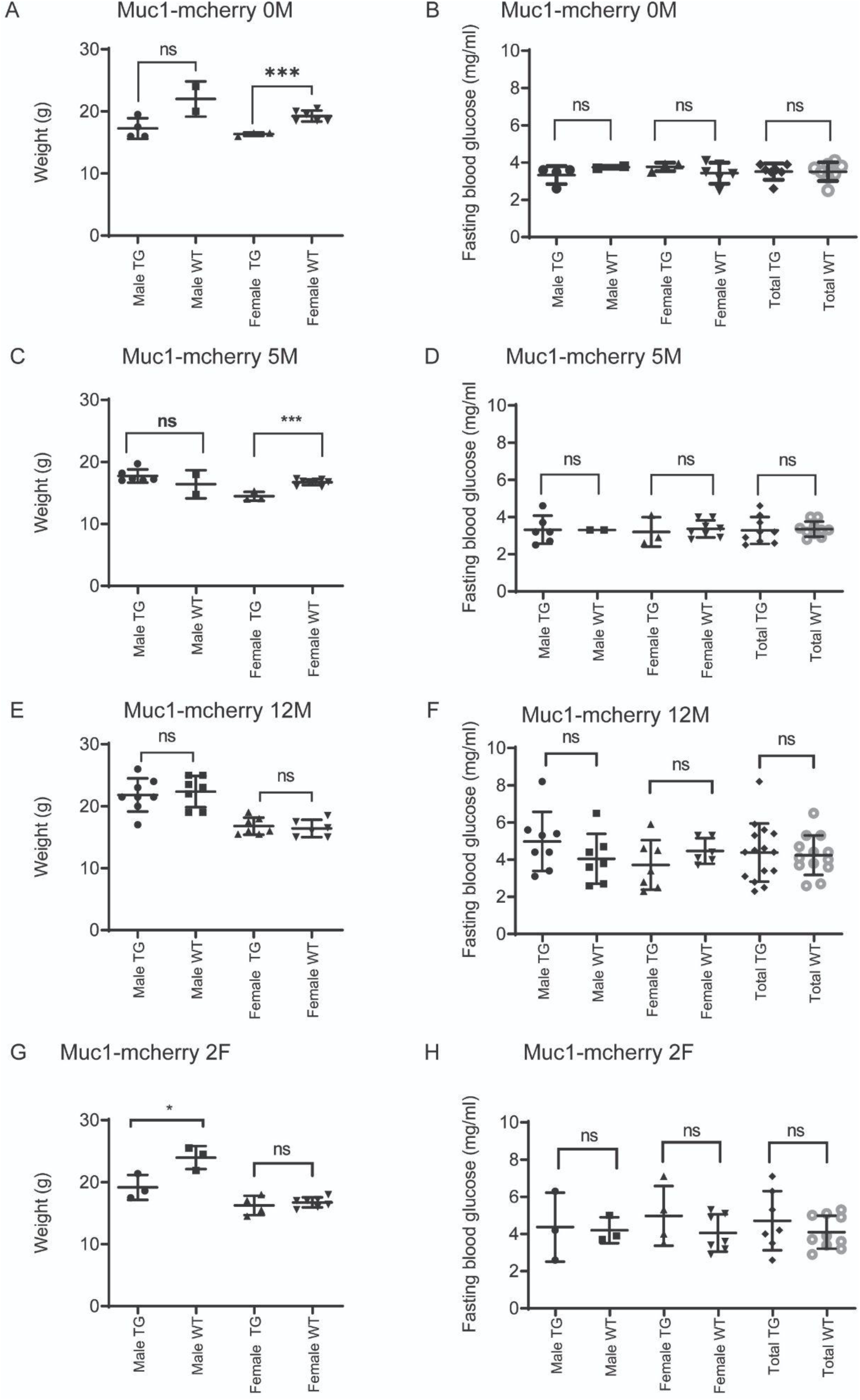
Weight and fasting blood glucose is not perturbed by the expression of the Muc1-mCherry transgene. **A:** Muc1-mCherry 0M females had reduced body weight (p=0,001 unpaired t-test), while males had a tendency for reduced body weight at 8-week of age compared to WT. **B:** Muc1-mCherry 0M had normal fasting blood glucose at 8-weeks. n=4 TG males, n=2 WT littermate males, n=3 TG females, n=6 WT females. **C:** Muc1-mCherry 5M females had a slightly reduced body weight at 8-week of age compared to WT (p=0,0002 unpaired t-test), while males had the same bodyweight. **D:** Muc1-mCherry 5M had normal fasting blood glucose at 8-weeks. N=6 TG males, n=2 WT littermate males, n=3 TG females, n=8 WT females. **E:** Muc1-mCherry 12M mice had a normal body weight at 8-week of age compared to WT. **F:** Muc1-mCherry 12M had normal fasting blood glucose at 8-weeks. N=8 TG males, n=7 WT littermate males, n=7 TG females, n=6 WT females. **G:** Muc1-mCherry 2F male mice had a slightly reduced body weight (p=0,04 unpaired t-test, while female mice had a normal body weight at 8-week of age compared to WT. **H:** Muc1-mCherry 2F had normal fasting blood glucose at 8-weeks. N=3 TG males, n=3 WT littermate males, n=4 TG females, n=7 WT females.

**SFigure 2:**
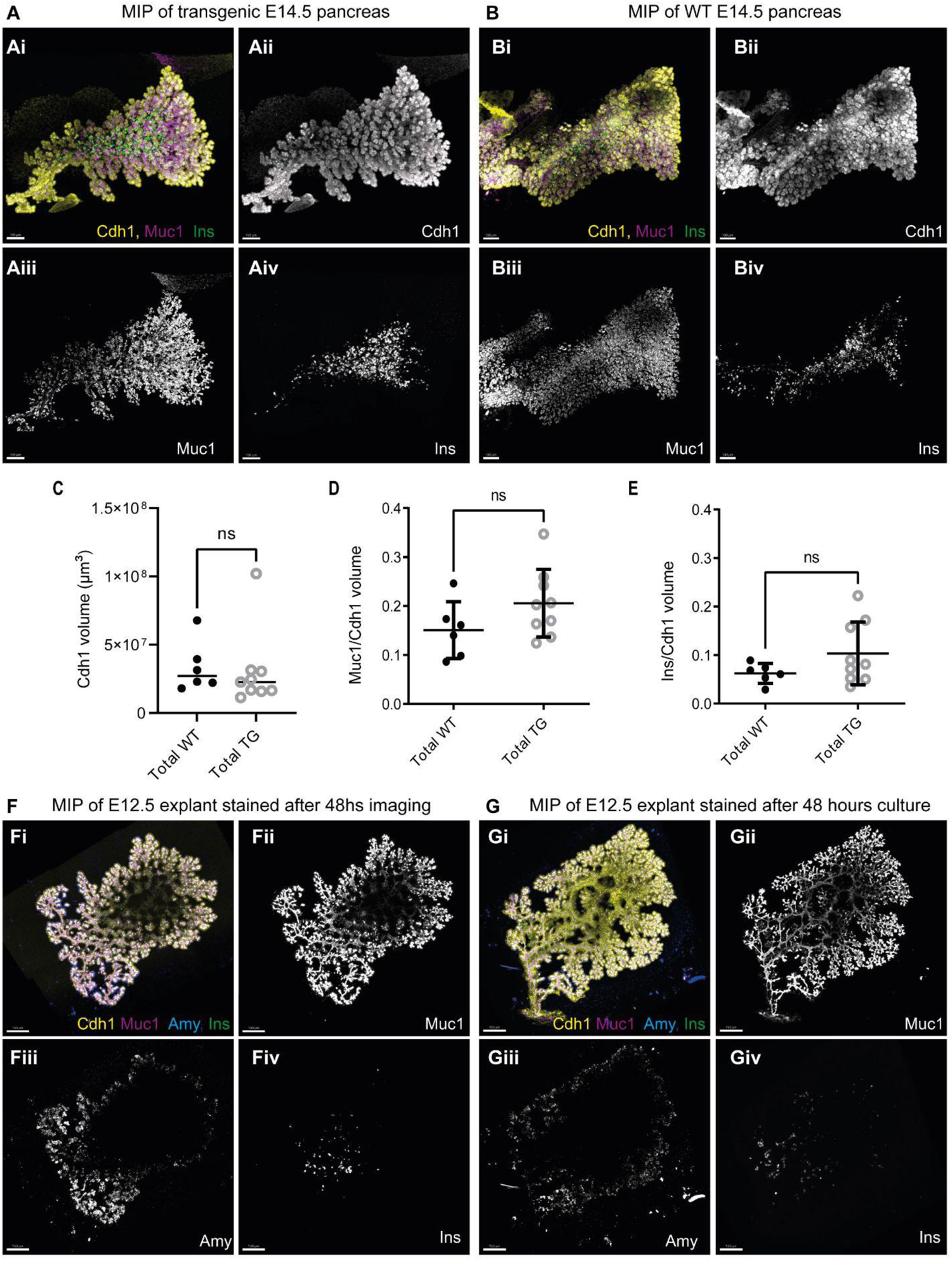
Muc1 expression and the differentiation of pancreatic precursors to mature cell types is not perturbed by the expression of the Muc1-mCherry transgene. **A-B :** Representative maximum intensity projections of WMIF of E14.5 Muc1-mCherry **(A)** and WT litter mate **(B)** pancreata stained for INS, CHD1 and MUC1. Bar=100µm in A-B. **C-D:** Quantification of antibody positive volumes as segmented using Imaris software. **C:** CHD1 total volume (µm3). **D:** MUC1 volume as fraction of CHD1 volume. **E:** INS volume as fraction of CHD1 volume. T-tests comparing all WT and all TG E14.5 embryos shows no significant difference between TG and WT. n=9 TG embryos and n=6 WT littermate embryos. n=2-4 embryos of 5M, 12M and 2F substrains were tested and the results pooled, as these substrains were not significantly different by one-way ANOVA (p=0,60 for MUC1/CDH1, and p=0,1 for INS/CDH1). **F-G:** Representative maximum intensity projection of IF staining of E12.5 explants from the 5M substrain after 4 days ex vivo growth including either 48 hours imaging **(F)** or 48 hours culture in the same on-stage incubator without imaging **(G)**. Stained for CHD1, MUC1, Amylase and INS. Bar=150µm in F-G.

**SFigure 3:**
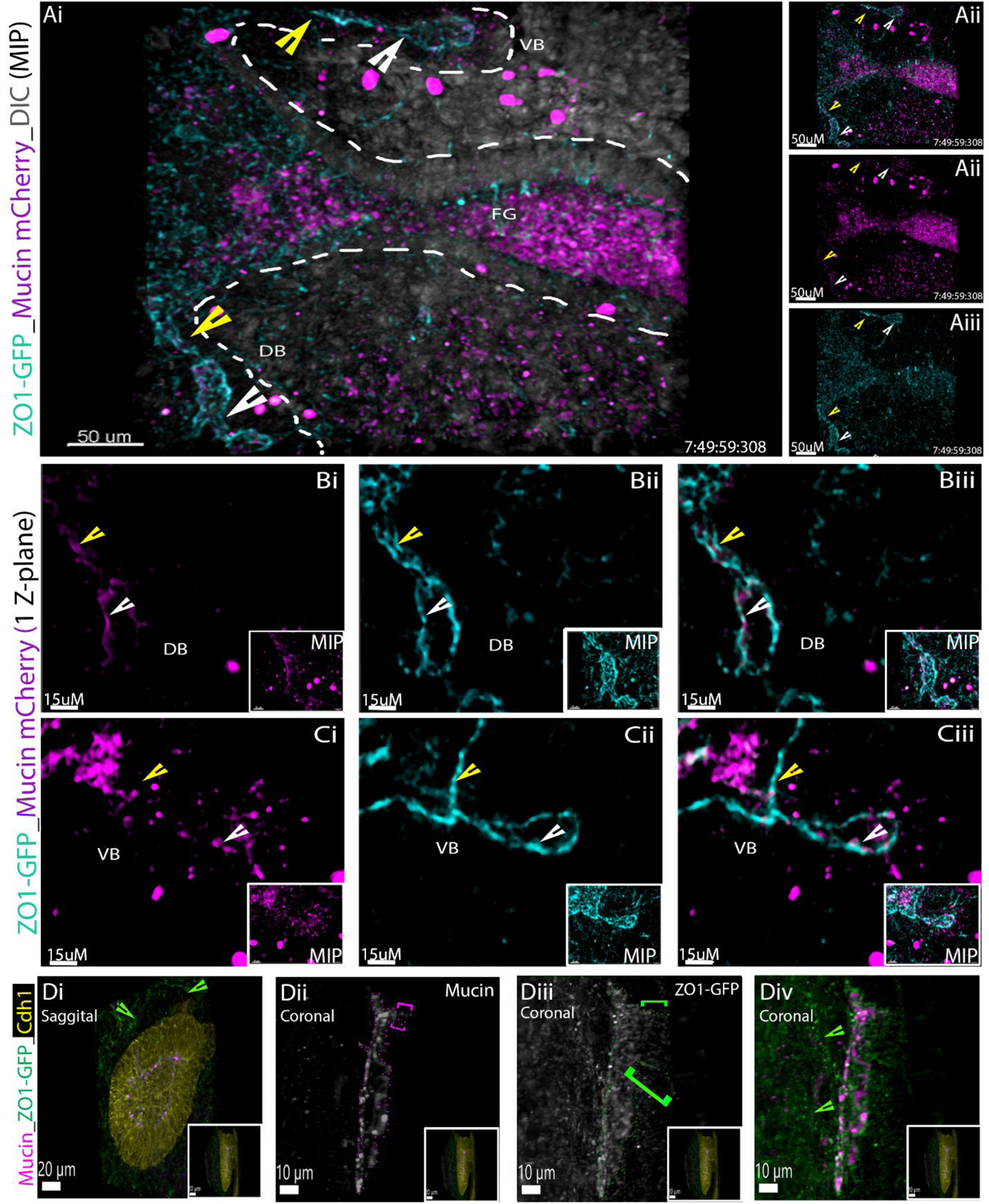
Hierarchy of endogenous and reporter visualized Mucin expression comparative to ZO1-GFP expression in live and WMIF pancreatic foreguts. **Ai-iii:** Representative time frames from 4D imaging of an E9.75 dorsal pancreatic bud expressing Muc1-mCherry (Magenta). The TIFFs depict the gradual onset of MUC1 expression (absent in time point T0 (ai, asterisk)), that begins as distal foci (aii, white arrowheads) that over time expands proximally toward the foregut (i.e. toward the time stamp). White dashed outline marks the edge of the pancreatic epithelium approximated from the DIC channel (shown as inserts top right). **Bi:** Representative time frames from 4D imaging of an E10.5 foregut expressing Muc1-mCherry (Magenta) and ZO1-GFP (Cyan) reveals consistent ZO1-GFP expression and variable Muc1-mCherry reporter expression in dorsal and ventral buds (visible by DIC, outlined in white dashed line), **Bii-iv:** same frame as (B) with Muc1-mCherry (Magenta) and ZO1-GFP (Cyan) alone in tandem, or alone. **Ci-iii and Di-iii:** Magnified views of the expression data shown in B panels displayed as a single optical Z-plane focusing on the dorsal (C panels) and ventral (D panels) buds (inset shows the same area as a MIP). We observe very little Muc1-mCherry expression in the ventral bud CPL compared to the dorsal bud CPL (Di vs Ci). In contrast ZO1-GFP expression is expressed apically throughout both buds lumens (Dii vs Cii). Arrowheads highlight yellow CPL stalk and white CPL cap, VB= ventral bud, DB= dorsal bud. Ei-iv MIP of an E10.0 dorsal bud stained for endogenous MUC1, CDH1 and the reporter ZO1-GFP shows a similar hierarchy of MUC1 and ZO1-GFP expression in fixed in vivo samples. Eii shows MUC1 is absent in most of the CPL stalk (inner yellow dashed line and in the foregut, in contrast to ZO1-GFP was detected throughout the CPL stalk (inner yellow dashed line) and into the foregut. Green arrows highlight ZO1-GFP marked endothelial cells.

**SFigure 4:**
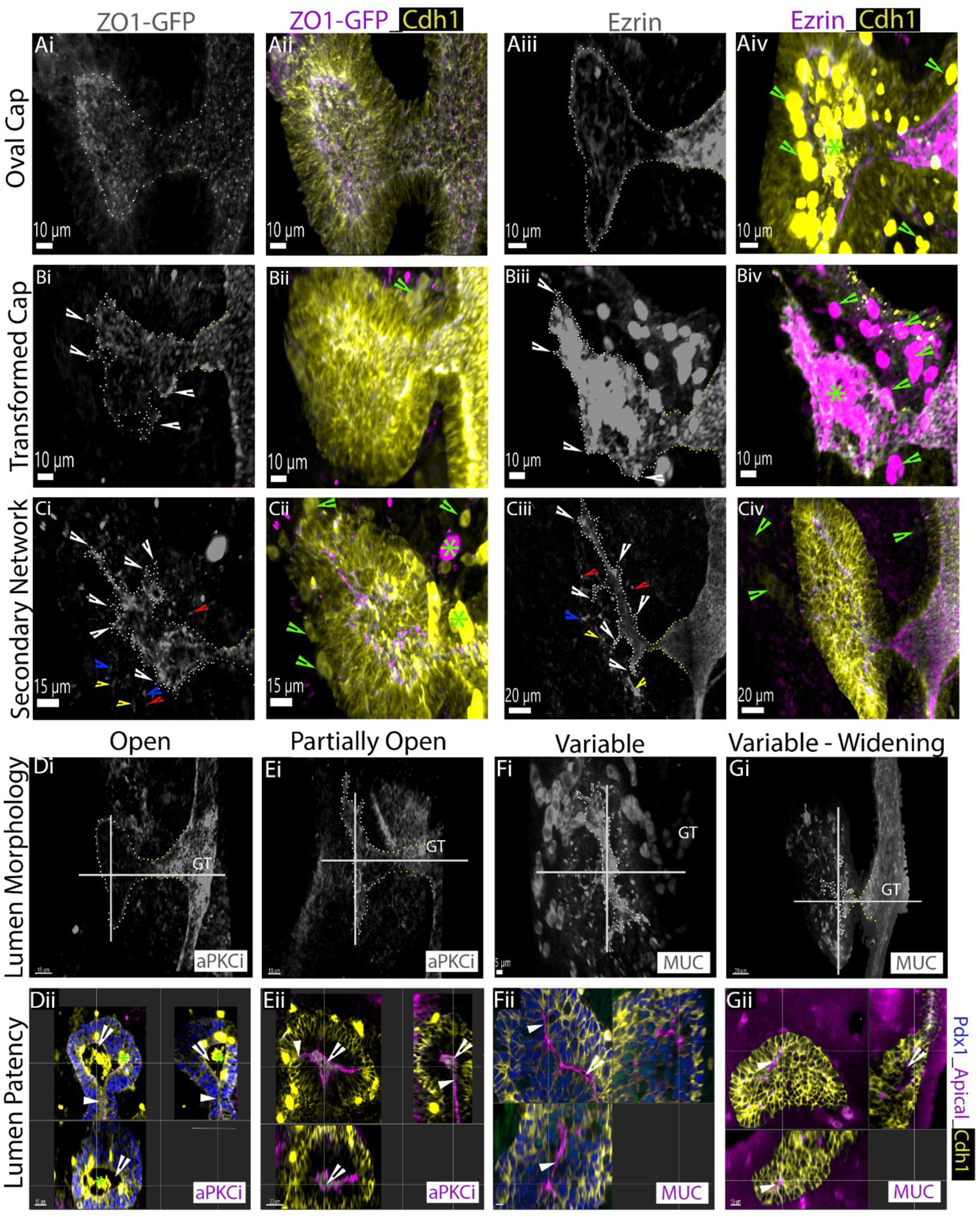
WMIF of multiple apical proteins reveal consistent CPL transitions and variable luminal patency during secondary network establishment. **A-C :** MIP of WMIF imaged embryonic foreguts (ranging from E10.0 to E11.0) stained for either ZO1-GFP (Ai, ii – Ci,Cii) or EZRIN (Ai, ii – Ci,Cii) depict similar CPL morphologies also visualized with MUC1 WMIF in Fig 2Di-iiii. Di-Gi and Hiii show MIP of WMIF imaged embryonic foreguts (ranging from E10.0 to E11.0) stained for various apical proteins to visualize different stages of CPL and network morphology. **Dii-Gii, Hi and Hii:** For each MIP we viewed the sample orthogonally and identified areas of luminal patency (open arrowhead) and lack of patency (filled arrowhead), both between samples and within samples. Panels Hi and Hii are orthogonal slices through the MIP (Hiii) at different levels of the CPL.

**SFigure 5:**
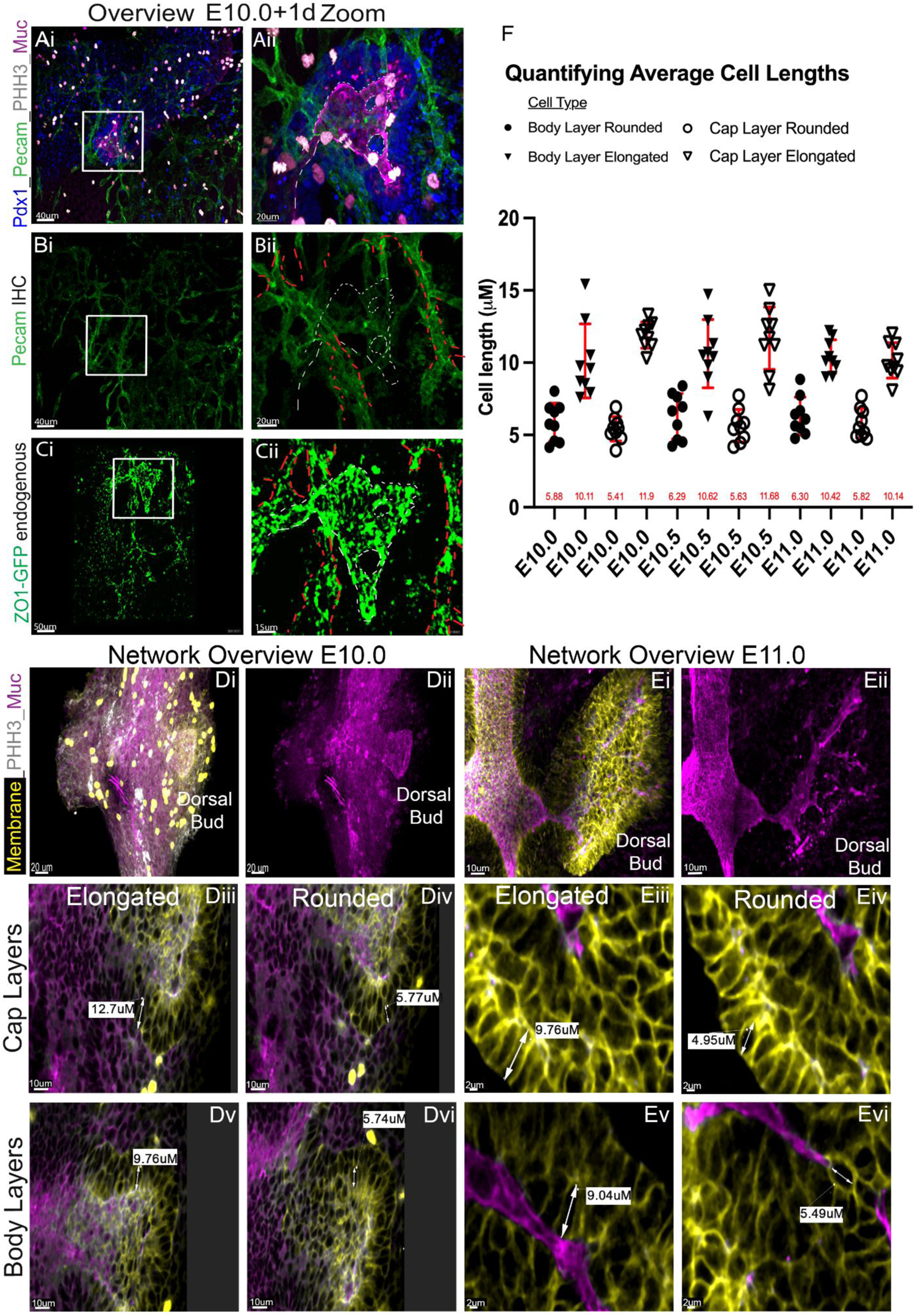
Identifying endothelial expression of ZO1-GFP outside the pancreatic epithelium and estimating the average cell length of a pancreatic epithelial cell. **A and B:** MIPs of an explant (i = overview of all tissue, ii = zoom in on pancreatic tissue) fixed directly after live imaging and WMIF stained for endothelial (PECAM, green), pancreas (PDX1, blue) and luminal (MUC1, magenta) proteins. Panel C is the last frame from the live imaging movie (endogenous ZO1-GFP signal, green). We found that the non-lumenal domains of ZO1-GFP in Ci and ii were comparable to the domain of the PECAM expression in Bi and ii (outlined in dashed red line). We could further see that the MUC1 expression domain (outlined in white dashed line in Bii) was absent from the PECAM expression. **E and F:** representative samples used to quantify average cell length Di, ii, Ei ii: MIPs of pancreata stained for membrane (D panels B-CATENIN, E panels ECDD2, yellow) markers and luminal markers (MUC1, magenta). From the cap and body layers of 3 independent E10.0, E10.5 and E11.0 dorsal buds, 3 rounded and 3 elongated cell lengths were measured. Diii-iv and Ei ii are representative optical Z-slices through the MIP illustrating examples of how cell measurements were performed. **F:** Scatter plot of cell length measurements, including the mean (red values along the X-axis) and SD (bars in red) of lengths grouped by cell morphology (elongated or rounded) and cell layer (cap or body) for E10.0-E11.0 dorsal buds samples. The average cell length of all the data was 8.35uM and the average of the longest (elongated cells) was 10.81uM, thus we settled on a length of more than 10uM to be a reasonable cell length cut off for categorizing a foci as de novo generated.

**SFigure 6:**
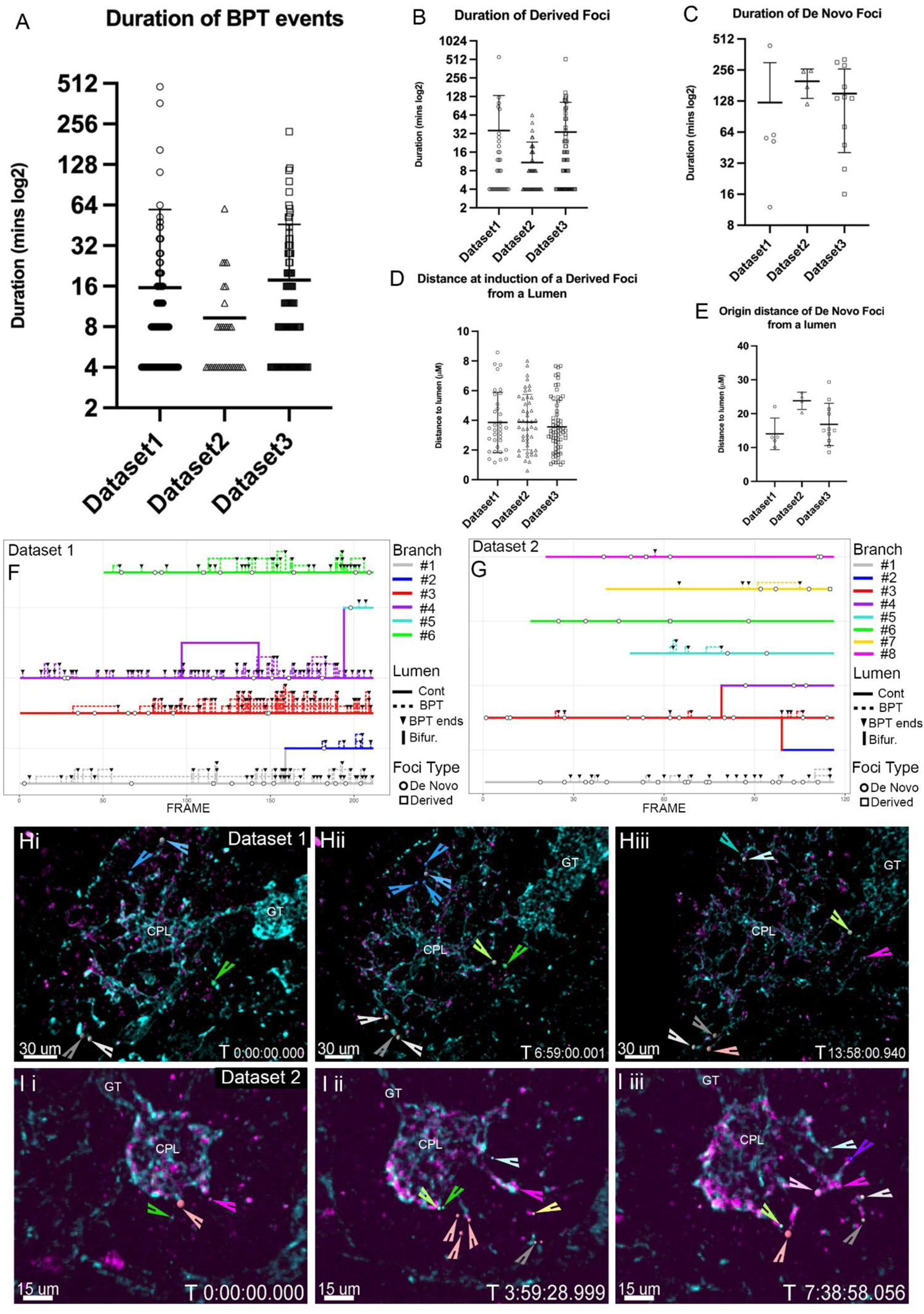
Quantitative data to further support the conclusion that lumenal network is largely derived through expansion and rearrangement of apical membranes. **A-C :** Scatter Plots visualizing the duration (i.e. length of time) - respectively - of every BPT, derived and de novo foci tracked in 3 live imaged explants forming a secondary network. **D-E**: Scatter Plots visualizing at detection, the distance - respectively - of every derived and de novo tracked foci to a pre-existing lumen in 3 live imaged explants forming a secondary network. All scatter plots display the mean and SD bars. **F and G:** Tracking plots displaying the frequency of tracked lumenal events, occurring over time, to contribute to the proximal distal growth and transformation of discrete lumenal branches in two additional explants. The plot visualizes the high frequency of lumenal breaking events (dashed lines, downward arrow marks the point at which the lumen unifies) and derived foci (unfilled circles) are observed during lumen growth. In contrast de novo foci (unfilled square) occur relatively infrequently. **Hi-iii:** Representative time frames from 4D imaging of the remodeling lumenal pancreatic network expressing ZO1_GFP (Cyan) and Muc1-mCherry (magenta). The TIFFs shown are a visual representation of the quantitative tracking data shown in F and data set 1 in Fig 2F. **Gi-iii:** Representative time frames from 4D imaging of the remodeling lumenal pancreatic network expressing ZO1_GFP (Cyan) and Muc1-mCherry (magenta). The TIFFs shown are a visual representation of the quantitative tracking data shown in G and data set 2 in Fig 2F. In panels H and G each spot color represents a different lumenal branch, light spots mark lumenal extension, dark spots mark breaking events and foci. Imaris tracking spots are further highlighted with an arrow of the same color.

**SFigure 7:**
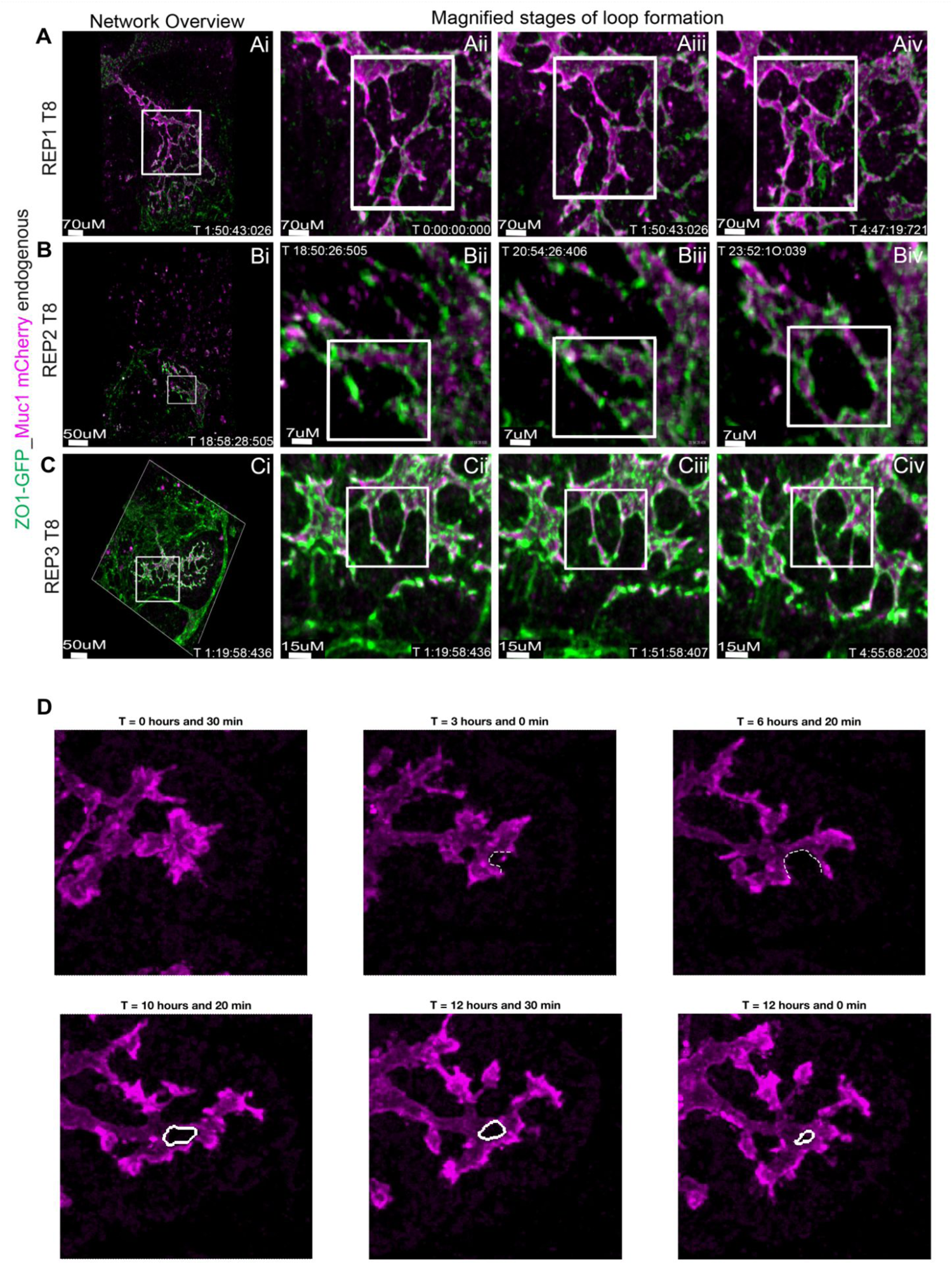
Loop formation is a continuous feature through early and mid-gestation lumenal network transformations. **A-C:** Representative time frames showing loop formation in the 4D imaging of pre-plexus pancreatic networks found in early gestation pancreata expressing ZO1_GFP (Green) and Muc1-mCherry (magenta). A-Ci gives an overview of the whole pancreatic explant within the imaging frame. A-Cii-iv are magnified areas, (white box in Ai-Ci) in which loops are forming over time, (inside the white box). A-C are 3 independent biological explants, in C one can see that the loop forms transiently and is not present in Civ. **D:** Example of loop formation in live *ex vivo* data. Magenta: MUC1, white dashed line: lumen outline before loop formation. White line: inner loop segmented area.

**SFigure 8:**
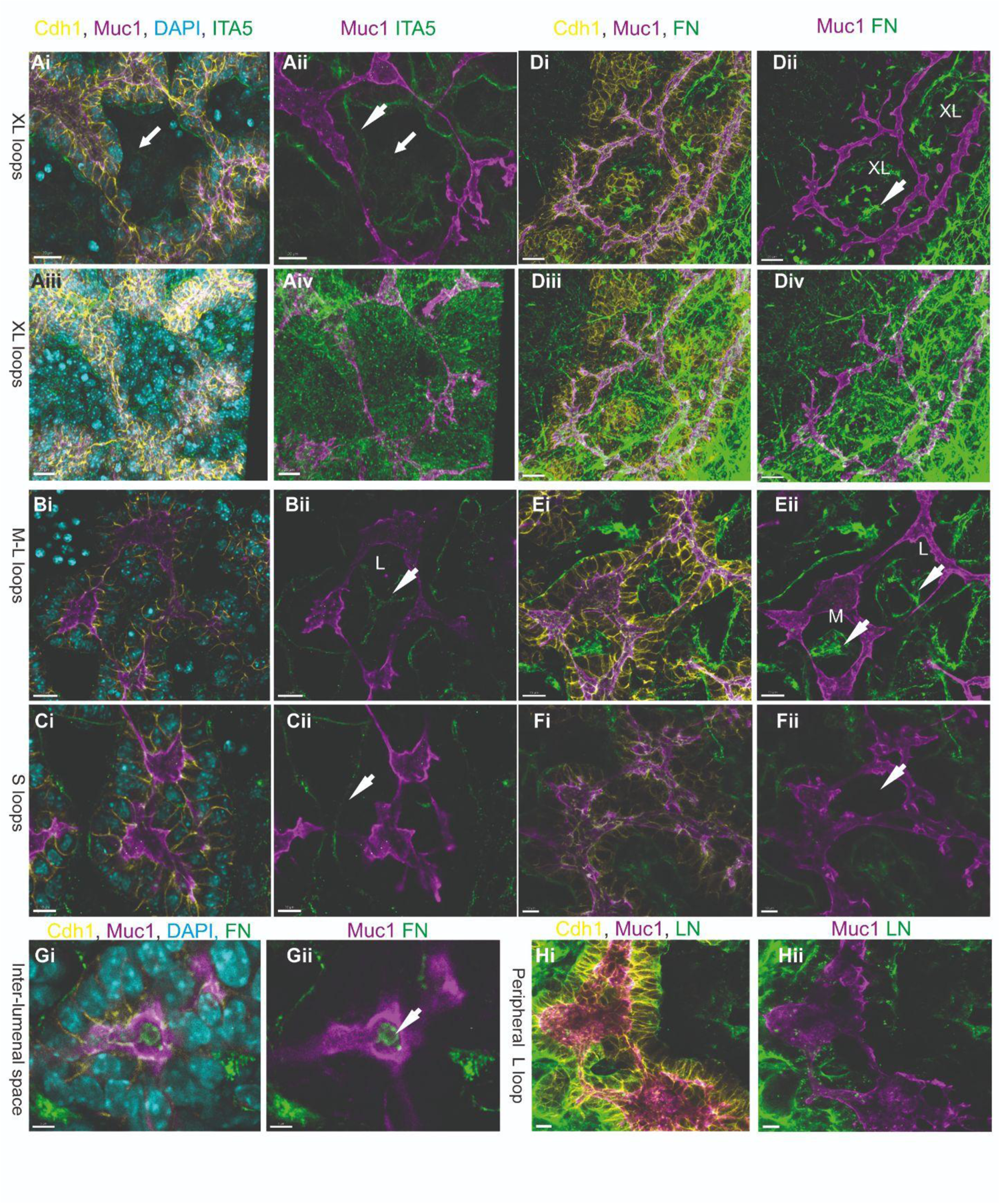
Representative maximum intensity projections (MIP) of loops in E11.5 + 3 days *ex vivo* pancreas. **Ai-Civ**: Stained for CHD1, MUC1, DAPI and ITG5. Ai-ii & Bi-Cii: MIP of z=5-6µm around loops making the inner hole visible. Arrow in Ai-ii: Mesenchymal cells. Arrowhead in Aii & B-Cii: ITA5 expression at the basal side of epithelial cells facing the basement membrane. Aiii-iv: MIP of the whole explant including regions above and below the loop. **Di-Gi:** Stained for CDH1, MUC1, Fibronectin (FN), and DAPI in Gi. Gi-ii shows a single 1µm section, all other images are MIPs of z=5-6µm around the loops. Arrowhead in Dii, Eii and Fii: FN expression in the inner hole of a loop. **Hi-ii:** Stained for CHD1, MUC1 and Laminin. MIPs of z=5-6µm. Bar is 20µm in Ai-ii, Di-iv and Hi-ii, 15 µm in Bi-ii and Ei-ii, and 10 µm in Ci-ii, Fi-ii & Gi-Hii.

**SFigure 9:**
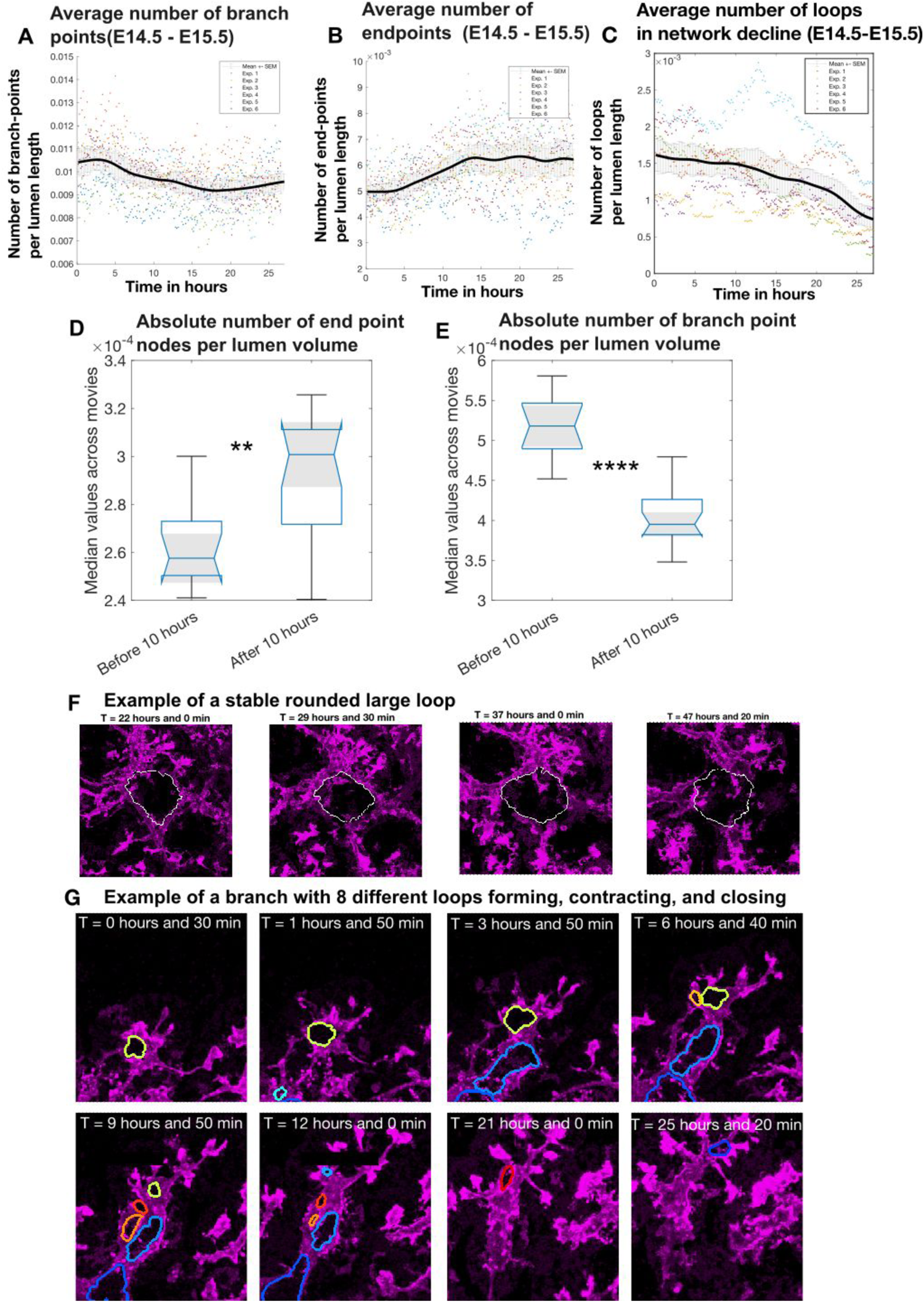
**A:** Data from segmentation and tracking of lumenal structures in 4D data. Colored dots: Number of branch points measured in the XY plane for each experiment/movie separately. Black line: Mean number of branch points across all experiments/movies with moving mean smoothing (window size 30 timepoints) +-SEM. Pearson’s correlation coefficients (ρ) between mean number of branch points (BPs) per lumen length and time for each experiment/movie separately are [−0.5075 −0.4790 −0.3447 −0.6437 −0.7963], median ρ = −0.51, (all p values are below 1.0e-5). **B:** Data from segmenting and tracking lumenal structures in live data. Colored dots: Number of end points measured in the XY plane for each experiment/movie separately. Black line: Mean number of end points across all experiments/movies with moving mean smoothing (window size 30 timepoints) +-SEM. Pearson’s correlation coefficients (ρ) between mean number of branch points (BPs) per lumen length and time for each experiment/movie separately are [0.5781 0.3593 0.7327 0.5270 0.6805], median ρ = 0.5755, (all p values are below 0.003). **C:** Colored dots: Median number of loop structure per lumen length for each experiment/movie separately. Black line: Mean number of loops per lumen length across all experiments/movies with moving mean smoothing (window size 30 timepoints) +-SEM. Pearson’s correlation coefficients (ρ) between median number of loops and time for each experiment/movie separately are [−0.8848 −0.9327 −0.2786 −0.9659 −0.6325 −0.9624], median ρ = - 0.78, (all p values are below 0.001). **D-E:** Analysis of absolut number of end points (C) and branch points (D) before and after 10 hours of imaging normalized with total lumen length or total lumen volume visible in the movies. n=6 experiments. p= 0.0012 (C) and p= 5.44e-6 (D) with Wilcoxon rank sum test. **G:**. Example of a branch with 8 different loops forming, contracting, and closing over time. Magenta: MUC1. Colored lines: segmentation of loops.

**SFigure 10:**
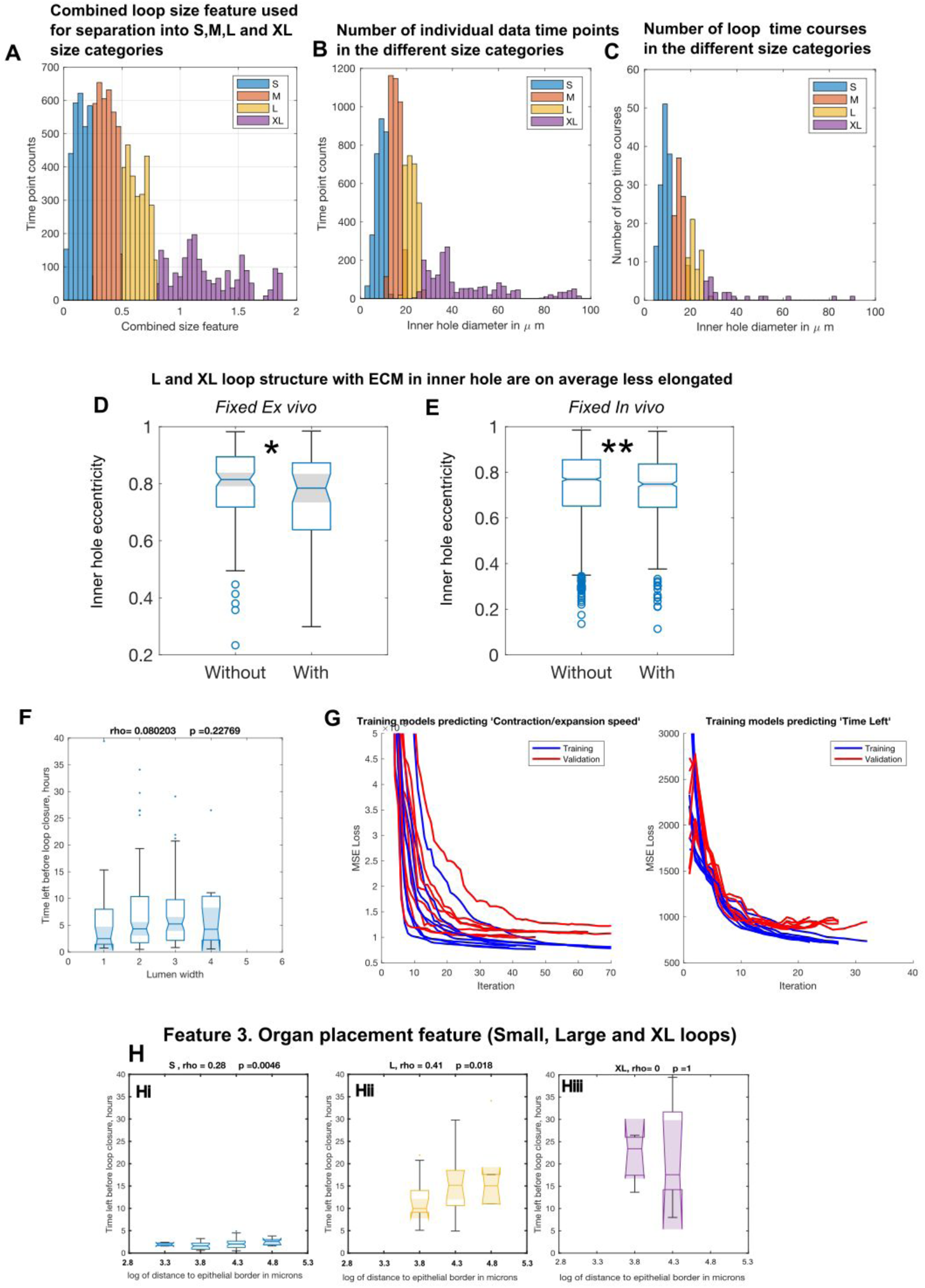
**A-C:** Distribution of loop sizes. We studied dynamic loop behavior in 4 size categories (S, blue M, orange L, yellow and XL, purple). Number of data time points per size category: small n= 2986, medium n= 3708, large n= 2706, extra large n=2122. Number of loop time courses in the different size categories: S n=133, M n= 95, L n= 54, XL n=24. Number of loop time courses in the different size categories ending with loop closure: S n=114, M n= 72, L n= 33, XL n=9. **A:** The combined size feature was used when dividing the loop data into size categories (S x>0.25, M 0.25<x <0.5, L 0.5<x <0.8, XL 0.8<x). The comb. size feature is a linear combination of the following highly correlated features: *’inner hole diameter’, ‘convex area’, ‘perimeter of convex hull’, ‘minor axis length’, ‘major axis length’, ‘lumen loop and hole total area’*. **B:** Inner loop hole equivalent diameter (the diameter of a circle with the same area as the segmented inner loop hole). Distribution of time points. **C:** Inner loop hole equivalent diameter (the diameter of a circle with the same area as the segmented inner loop hole). Distribution of loop time courses. **D:** Fixed *ex vivo* data of lumens and epithelium E14.5 *in vivo* pancreas (n=3) shows that L and XL lumenal loops without non epithelial cell types present in the middle are more elongated. p value 0.028 (Wilcoxon rank sum test). **E:** Fixed *in vivo* data of lumens and epithelium E14.5 *in vivo* pancreas (n=7) shows that L and XL lumenal loops without non epithelial cell types present in the middle are more elongated. p value 0.003 (Wilcoxon rank sum test). **F**:Time left before loop closure is not significantly associated with average lumen width in loop. **G:** Mean squared error loss monitored during training of 8 NN models predicting either ‘Time left before loop closure’ or ‘loop contraction speed’. Training was stopped if the validation loss was smaller than the minimum validation loss recorded so far, for more than 6 iterations in a row, (since this indicates that further training is overfitting the model on the training data and causing an increase in validation loss). **H:** We found a positive correlation between *time left* and *mean distance to epithelial/mesenchymal interface*. Since there is also a slight positive correlation between category and distance to the organ edge we look at correlation in our size categories separately (S Hi, L Hii, XL Hiii, see figure 6H for M). The strongest effect of placement in the organ on *time left* seems to be in the M and L groups, where Spearman’s rho is >0.4. Number of time courses in each category is, Small: n=114. Medium: n=72. Large: n=33. XL n=9. Time left before loop closure in XL loop size groups is not significantly associated with placement in the organ.

**SFigure 11:**
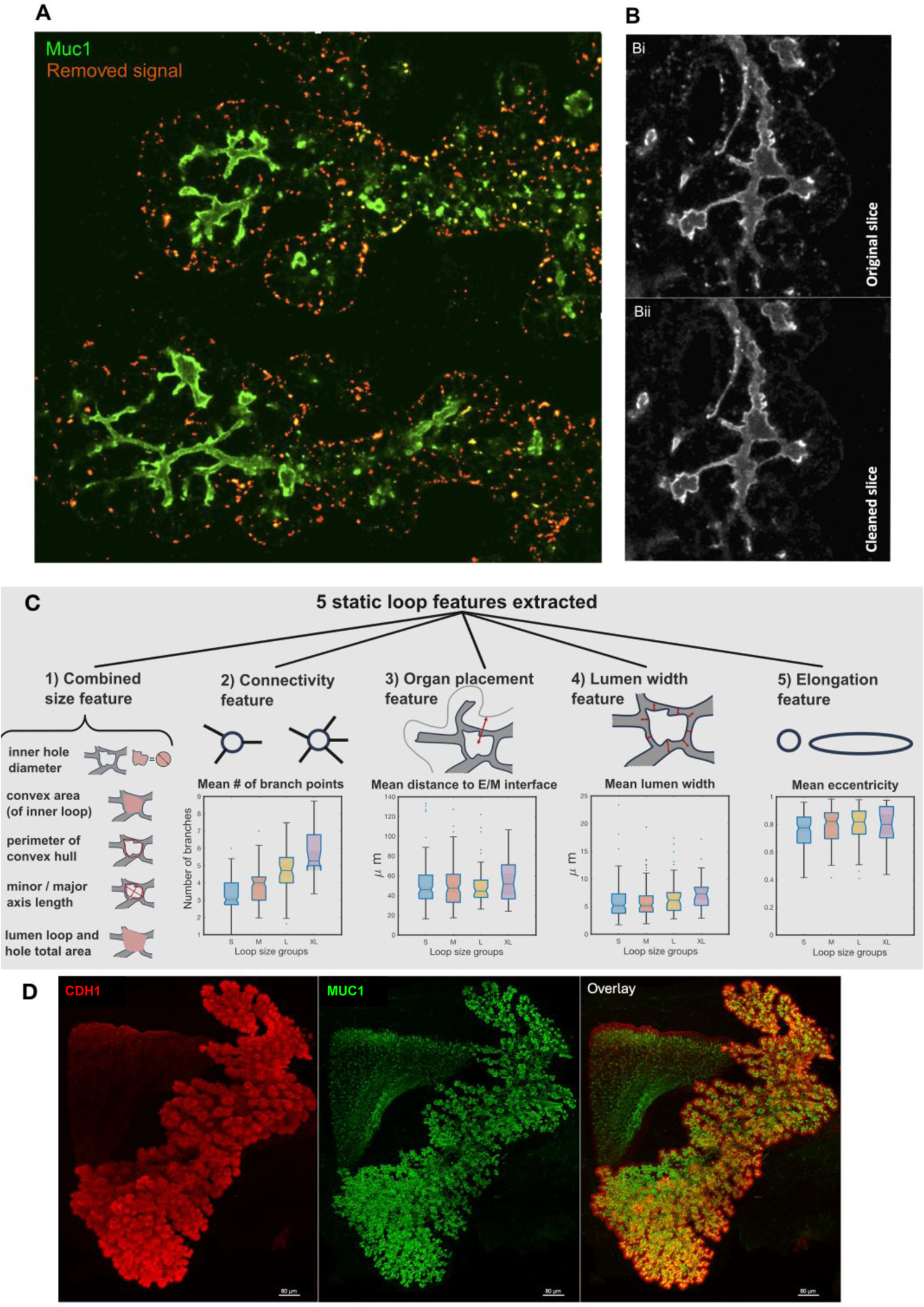
**A:** Single slice (middle of z stack) from live movie time point. Green Muc1 one signal. Orange/yellow: Areas where Muc1 signal was removed. **B:** Cleaned slice from live movie time point before (Bii) and after (Bi) removal of small round bright Muc1 signals. **C:** Overview of the 5 static loop features extracted from the live imaging data. 1) Combined size feature. The loop features that are most correlated with both the two long term and short term target variables (*time left* and *contraction speed*) are those that quantify loop size in different ways. We therefore designed a combined size feature and divided our loop time courses into 4 size categories (S, M, L, XL see also SFigure 10) in order to study the effect of other static features relating to shape and placement in the organ within groups of similar loop size. The original six size features contributing to the combined size feature were: *’inner hole diameter’, ‘convex area’, ‘perimeter of convex hull’, ‘minor/major axis length of an ellipse fitted to the inner loop hole segmentation’, ‘major axis length’, ‘lumen loop and hole total area’*, z scored and weighted in the new combined size feature according to their correlation with the two target variables. 2) Connectivity feature, the mean number of branch points/outgoing lumens. 3) Organ placement feature, the mean distance to epithelial/mesenchymal interface in microns. 4) Lumen width feature, the mean lumen width in microns. 5) Elongation feature, the eccentricity of an ellipse fitted to the inner loop hole segmentation. **D**: MIPs of 3D images used to produce data shown in figure 7A. Scale bar signifies 80 microns.

## Movie legends

Movie metadata is included in STable1

***Movie 1:* ZO1-GFP reporter expression illustrating CPL cap and stalk formation by lumenal evagination in the ex vivo pancreatic bud.**

A posterior foregut explant, expressing ZO1-GFP (cyan), dissected from an E9.75 embryo was imaged every 8 minutes. The movie depicts the CPL (stalk and cap) budding out from and perpendicular to the foregut over a 20 hour period. Unlike the pattern of Muc1-mCherry expression, ZO1 GFP expression marking the central primary lumen is ubiquitous and continuous, with no peripheral signal appearing de novo during the imaging period. White outline marks the budding CPL. Annotations: DB = Dorsal Bud, VB= Ventral Bud, CPL = central primary lumen, Green arrow heads are examples of ZO1-GFP expressing endothelial cells moving stochastically and at higher frequency in comparison to the lumenal ZO1-GFP expression in the lumen.

***Movie 2:* Hierarchy of MUC1-mCherry expression comparative to ZO1-GFP expression in live pancreatic foreguts.**

A posterior foregut explant, expressing Muc1-mCherry (Magenta):ZO1-GFP (cyan), dissected from an E10.5 embryo was imaged every 8 minutes. The movie reveals the expression of ZO1-GFP to be consistent/ubiquitous through the lumen of dorsal and ventral buds. In contrast, expression of the Muc1-mCherry reporter appeared variable and to have a maturity in its onset.. Annotations: Arrowheads highlight yellow CPL stalk and white CPL cap, VB= ventral bud, DB= dorsal bud. Green arrow marks autofluorescence from red blood cells.

***Movie 3:* In vivo 3D lumenal morphologies are found to be equivalent, at comparative developmental stages, to the 4D ex vivo budding topology of the CPL observed through live imaging (Movie 1).**

***3D Clip 1*:** E9.0 in vivo pancreatic bud in which CPL evagination (white outline) has just begun, echoing the shape of the outer epithelium. ***3D Clip 2***: an E9.5 in vivo pancreatic bud with more pronounced CPL evagination, (white outline) at the distal tip of the lumen dorsal ventral cap widening has begun (CPL bending, magenta arrows). ***3D Clip 3:*** E10.0 in vivo pancreatic bud in which budding is complete and the CPL has transformed/grown to form an oval cap (white outline) and stalk (yellow outline) CPL morphology. The magenta arrows denote extensions from the CPL, no peripheral foci or lumens are visible. **All movies:** Ezrin expressing lumens in magenta, Cell membranes expressing CDH1 in yellow and Pdx1 expressing nuclei in blue. Annotations: FG = foregut epithelium, CPL = central primary lumen VB= ventral bud, DB= dorsal bud. Green arrows are examples of autofluorescence from red blood cells (RBC’s).

***Movie 4:* 4D secondary network generation through lumenal expansion and rearrangement initiated at the CPL.**

A posterior foregut explant, expressing Muc1-mCherry (Magenta):ZO1-GFP (cyan), dissected from anE10.5 embryo imaged every 8 minutes for ca.20 hours. The imaged ventral pancreatic bud depicts the transformation of the CPL into a pre-plexus secondary network with multiple interconnected lumens. Arrows/spots highlight examples of different transformative topological events observed in the 4D imaging dataset (white = extension, yellow = extension becoming a BPP event, red = De Novo Foci, Blue = derived foci). Green arrows are examples of autofluorescence from red blood cells.

***Movie 5: I*n vivo 3D lumenal morphologies are found to be equivalent, at comparative developmental stages, to the 4D transformation and extension of the CPL documented in Movie 4.**

Whole Mount ImmunoFluorescence (WMIF) visualizes the morphologies of the lumenal network in 3D static data during secondary network establishment. ***3D Clip 1*:** WMIF in an E10.5 in vivo pancreatic bud illustrates transformation and extension of the CPL, secondary lumenal sites primarily have a foci/microlumen like morphology. ***3D Clip 2:*** WMIF in an E10.5 in vivo pancreatic bud illustrates a highly transformed CPL secondary lumenal sites are a mix of peripheral lumens (reminiscent of break points) and microlumens and foci. ***3D Clip 3*:** WMIF in an E11.0 in vivo pancreatic bud illustrates in 3D an elaborate, abundant and highly disjoined secondary lumenal network with peripheral sites of varying sizes. ***3D Clip 4*:** WMIF in an E11.5 in vivo pancreatic bud illustrates in 3D a secondary lumenal network with a more connected, pre-plexus-like appearance in which the secondary lumenal sites are also beginning to branch and unify with other secondary sites. All movie panels: MUC1 expressing lumens in magenta, Cell membranes expressing CDH1 in yellow and PDX1 expressing nuclei in blue. Annotations: CPL is more clearly defined by white (CPL) and yellow (FG) dashed lines. Arrows in all movies denote topologies similar to the transformative lumenal events seen in dynamic (4D) datasets: White large open = extension, white small closed = bifurcation, red = De Novo, Blue =Derived, Yellow = BP. Green arrows are examples of autofluorescence from red blood cells.

***Movie 6*: Zoomed in examples of tracked lumenal events that occur during secondary network transformation.**

Pancreatic foregut explants, expressing Muc1-mCherry (Magenta):ZO1-GFP (cyan), dissected from an E10.5 embryo imaged every 4 or 8 minutes, DIC gray. **4D Clip 1 - Extension and bifurcation:** Lumen of interest is highlighted and tracked by gray spots, intermittent white arrows highlight extension, intermittent unfilled arrow highlights a merging event which gives a bifurcated lumen. **4D Clip 2 - Break Point Temporary (BPT)** of an initially unified CPL extension (white arrow), merges with another lumen (pink arrow, 14hr57), then fragments (grey and cream arrow, ca 16hr50) and eventually reunifies (white arrow), as highlighted and tracked by spots. **4D Clip 3 - Break Point Permanent (BPP)** The lumen of interest which extends from the CPL is initially unified, fragments and remains discontinuous (BPP). A gray spot initially highlights and tracks the CPL extension. At 4hrs9 the spot tracks the BPP as the extension fragments and at the end of imaging (6hr39) remains disconnected. **4D Clip 4 - Derived and de novo foci** A red spot and arrow marks and tracks a de novo foci (>10uM, de novo) measured (T0hr) 23.6uM from the CPL, which at the end of imaging had contributed to the peripheral lumenal network. A blue spot and arrow marks and tracks a derived foci (<10uM, derived)) measured (T2hr31) 6.64uM from the CPL, which temporarily contributes to the secondary lumenal network but after T10hr29 fragments and is unfeasible to track. **In all clips: insets, top right/left of the movies:** Overview of whole explants growth and transformation shown, zoom in of lumenal event highlighted by yellow box.

***Movie 7:* Tracking and lumenal events that can contribute to branch growth and transformation during secondary network formation.**

Three independent posterior foregut explants, expressing Muc1-mCherry (Magenta):ZO1-GFP (cyan), dissected from embryos imaged every 4 minutes, DIC gray. Each movie visualizes the real time tracking events displayed graphically in the tracking plots and event frequency graphs (Fig2 and sFig6). Each spot colour tracks the events within and growth of an individual lumenal branch. Light spots mark lumenal extension, dark spots mark breaking events and foci. **Left: dataset 2 -**The lumenal network is at the earliest stage of secondary network establishment i.e at T0 the CPL is not transformed and there are only foci, no lumens, peripherally. **Middle: dataset 3** - The lumenal network is at the beginning of secondary network transformation i.e at T0 the CPL is transformed/branched but only very few lumens are established peripherally (e.g. branch 7, yellow spot). **Right: dataset 1 -**The secondary lumenal network is in a period of growth and continued transformation i.e at T0 the CPL is already very branched and multiple peripheral lumens are established (e.g. branch 1, 3 and 4 respectively gray, red and purple spots. Gray arrows are examples of autofluorescence from red blood cells, Green arrow heads are examples of ZO1-GFP expressing endothelial cells moving with high frequency in comparison to the lumenal ZO1-GFP expression in the lumen.

***Movie 8:* Loop formation is a continuous feature through early and mid-gestation lumenal network transformation*s*.**

Two independent posterior foregut explants, expressing Muc1-mCherry (Magenta):ZO1-GFP (green), dissected from embryos imaged every 8 minutes, DIC gray. Each movie visualizes loop formation in the 4D imaging of pre-plexus pancreatic networks found in early gestation pancreata. Coloured boxes highlight areas where loops are forming, numbers record the amount of loops generated, bold numbers mark loops at the end of imaging which were outside of the field of view during imaging.

**Movie 9: *Example of loop forming and closing in Muc1-mCherry expressing mid-gestation pancreas***

Partial view of dorsal pancreas explanted at E12.5 and grown *ex vivo* for 48 hs, followed by live time lapse imaging for up to 48 hs, expressing Muc1-mCherry (Magenta), imaged every 10 min. Movie shows maximum intensity projection (MIP) of z stack. White line is drawn along the inner rim of the loop structure and is inserted at the time the loop forms and disappears again when the loop closes. This loop belongs to the small size category. White scale bar shows 50 microns.

**Movie 10: Loop segmentation and tracking in Muc1-mCherry expressing mid-gestation pancreas**

This movie relates to Fig 5A-B. Dorsal pancreas explanted at E12.5 and grown *ex vivo* for 48 hs, followed by live time lapse imaging for 48 hs (every 10 min), expressing Muc1-mCherry (Magenta). Movie shows maximum intensity projection (MIP) of the z stack. Loop structures were tracked and segmented. Lines (with unique color for each loop) are drawn along the inner rim of the loop structure and is inserted at the time a loop forms and disappears again when the loop closes/breaks. Note that there may be loop structures besides those highlighted in the movie. The ones highlighted have all been inspected and verified as bonafide loop structures in 3D. For some parts of the tissue, the thickness was too high and signal thus too low to properly distinguish loop structures. White scale bar shows 50 microns.

**Movie 11: Example of loop cinching in Muc1-mCherry mid-gestation pancreas**

This movie relates to Fig 6D. Dorsal pancreas explanted at E12.5 and grown *ex vivo* for 48 hs, followed by live time lapse imaging for up to 48 hs (every 10 min), expressing Muc1-mCherry (Magenta). This movie relates to Fig 6d. Movie shows maximum intensity projection (MIP) of the z stack.

A white line is drawn along the inner rim of the loop structure that cinches. The white line is inserted at the time a loop forms and disappears again when a loop closes. Initially this loop belongs to the large size category. After cinching the two newly formed loops belong to the medium and small size categories respectively. At approximately 20 hours the medium loop passes into the small size category and finally closes about 3 hours later. White scale bar shows 50 microns.

**Movie 12: Example of one branch with 8 different loops forming, contracting, and closing over time in Muc1-mCherry mid-gestation pancreas**

This movie relates to SFig 9E. Dorsal pancreas explanted at E12.5 and grown *ex vivo* for 48 hs, followed by live time lapse imaging for 48 hs (every 10 min), expressing Muc1-mCherry (Magenta). Movie shows maximum intensity projection (MIP) of the z stack. Loop structures were tracked and segmented. Lines (with unique color for each loop) are drawn along the inner rim of the loop structure and is inserted at the time a loop forms and disappears again when the loop closes. White scale bar shows 50 microns.

**Movie 13: Example of a loop structure contracting and closing (lower right corner)**

This movie relates to Fig 6B. Dorsal pancreas explanted at E12.5 and grown *ex vivo* for 48 hs, followed by live time lapse imaging for 48 hs (every 10 min), expressing Muc1-mCherry (Magenta). The movie shows maximum intensity projection (MIP) of the z stack. A white line is drawn along the inner rim of the loop structure that contracts and closes. White scale bar shows 50 microns.

**Movie 14: Example of a loop structure breaking (upper left corner)**

This movie relates to Fig 6C. Dorsal pancreas explanted at E12.5 and grown *ex vivo* for 48 hs, followed by live time lapse imaging for 48 hs (every 10 min), expressing Muc1-mCherry (Magenta). The movie shows maximum intensity projection (MIP) of the z stack. A white line is drawn along the inner rim of the loop structure that breaks. The white line disappears at the time point when the loop breaks. White scale bar shows 50 microns.

**Movie 15: Example of an XL loop structure widening/maintaining size**

This movie relates to Fig 6A. Dorsal pancreas explanted at E12.5 and grown *ex vivo* for 48 hs, followed by live time lapse imaging for 48 hs (every 10 min), expressing Muc1-mCherry (Magenta). The movie shows maximum intensity projection (MIP) of the z stack. A white line is drawn along the inner rim of the XL loop structure. White scale bar shows 50 microns.

## References

Araújo, Sofia J., and Marta Llimargas. 2023. “Time-Lapse Imaging and Morphometric Analysis of Tracheal Development in Drosophila.” Methods in Molecular Biology 2608: 163–82.

Azizoglu, D. Berfin, Caitlin Braitsch, Denise K. Marciano, and Ondine Cleaver. 2017. “Afadin and RhoA Control Pancreatic Endocrine Mass via Lumen Morphogenesis.” Genes & Development 31 (23-24): 2376–90.

Bafna, S., S. Kaur, and S. K. Batra. 2010. “Membrane-Bound Mucins: The Mechanistic Basis for Alterations in the Growth and Survival of Cancer Cells.” Oncogene 29 (20): 2893–2904.

Bankaitis, Eric D., Matthew E. Bechard, and Christopher V. E. Wright. 2015. “Feedback Control of Growth, Differentiation, and Morphogenesis of Pancreatic Endocrine Progenitors in an Epithelial Plexus Niche.” Genes & Development 29 (20): 2203–16.

Barlow, Haley R., Neha Ahuja, Tyler Bierschenk, Yadanar Htike, Luke Fassetta, D. Berfin Azizoglu, Juan Flores, et al. 2023. “Rab11 Is Essential to Pancreas Morphogenesis, Lumen Formation and Endocrine Mass.” Developmental Biology 499 (July): 59–74.

Bergmann, Carsten, Lisa M. Guay-Woodford, Peter C. Harris, Shigeo Horie, Dorien J. M. Peters, and Vicente E. Torres. 2018. “Polycystic Kidney Disease.” Nature Reviews. Disease Primers 4 (1): 50.

Cano, David A., Noel S. Murcia, Gregory J. Pazour, and Matthias Hebrok. 2004. “Orpk Mouse Model of Polycystic Kidney Disease Reveals Essential Role of Primary Cilia in Pancreatic Tissue Organization.” Development 131 (14): 3457–67.

Chambers, J. A., M. A. Hollingsworth, A. E. Trezise, and A. Harris. 1994. “Developmental Expression of Mucin Genes MUC1 and MUC2.” Journal of Cell Science 107 (Pt 2) (February): 413–24.

Cnossen, Wybrich R., and Joost P. H. Drenth. 2014. “Polycystic Liver Disease: An Overview of Pathogenesis, Clinical Manifestations and Management.” Orphanet Journal of Rare Diseases 9 (May): 69.

Dahl-Jensen, Svend Bertel, Siham Yennek, Lydie Flasse, Hjalte List Larsen, Dror Sever, Gopal Karremore, Ivana Novak, Kim Sneppen, and Anne Grapin-Botton. 2018. “Deconstructing the Principles of Ductal Network Formation in the Pancreas.” PLoS Biology 16 (7): e2002842.

Darrigrand, Jean-Francois, Anna Salowka, Alejo Torres-Cano, Rafael Tapia-Rojo, Tong Zhu, Sergi Garcia-Manyes, and Francesca M. Spagnoli. 2024. “Acinar-Ductal Cell Rearrangement Drives Branching Morphogenesis of the Murine Pancreas in an IGF/PI3K-Dependent Manner.” Developmental Cell 59 (3): 326–38.e5.

Debnath, Jayanta, Kenna R. Mills, Nicole L. Collins, Mauricio J. Reginato, Senthil K. Muthuswamy, and Joan S. Brugge. 2002. “The Role of Apoptosis in Creating and Maintaining Luminal Space within Normal and Oncogene-Expressing Mammary Acini.” Cell 111 (1): 29–40.

Dimitriou, Ioannis, Anastasios Katsourakis, Eirini Nikolaidou, and George Noussios. 2018. “The Main Anatomical Variations of the Pancreatic Duct System: Review of the Literature and Its Importance in Surgical Practice.” Journal of Clinical Medicine Research 10 (5): 370–75.

Fanni, D., N. Iacovidou, A. Locci, C. Gerosa, S. Nemolato, P. Van Eyken, G. Monga, S. Mellou, G. Faa, and V. Fanos. 2012. “MUC1 Marks Collecting Tubules, Renal Vesicles, Comma- and S-Shaped Bodies in Human Developing Kidney Tubules, Renal Vesicles, Comma- and S-Shaped Bodies in Human Kidney.” European Journal of Histochemistry: EJH 56 (4): e40.

Flasse, Lydie, Coline Schewin, and Anne Grapin-Botton. 2020. “Pancreas Morphogenesis: Branching in and Then out.” Curr Top Dev Biol 143 (November): 75–110.

Foote, Henry P., Kaelyn D. Sumigray, and Terry Lechler. 2013. “FRAP Analysis Reveals Stabilization of Adhesion Structures in the Epidermis Compared to Cultured Keratinocytes.” PloS One 8 (8): e71491.

Hannezo, Edouard, Colinda L. G. J. Scheele, Mohammad Moad, Nicholas Drogo, Rakesh Heer, Rosemary V. Sampogna, Jacco van Rheenen, and Benjamin D. Simons. 2017. “A Unifying Theory of Branching Morphogenesis.” Cell 171 (1): 242–55.e27.

Heilmann, Silja, Henrik Semb, and Pia Nyeng. 2021. “Quantifying Spatial Position in a Branched Structure in Immunostained Mouse Tissue Sections.” STAR Protocols 2 (4): 100806.

Hick, Anne-Christine, Jonathan M. van Eyll, Sabine Cordi, Céline Forez, Lara Passante, Hiroshi Kohara, Takashi Nagasawa, et al. 2009. “Mechanism of Primitive Duct Formation in the Pancreas and Submandibular Glands: A Role for SDF-1.” BMC Developmental Biology 9 (December): 66.

Hogan, Brigid L. M., and Peter A. Kolodziej. 2002. “Organogenesis: Molecular Mechanisms of Tubulogenesis.” Nature Reviews. Genetics 3 (7): 513–23.

Jørgensen, Mette Christine, Jonas Ahnfelt-Rønne, Jacob Hald, Ole D. Madsen, Palle Serup, and Jacob Hecksher-Sørensen. 2007. “An Illustrated Review of Early Pancreas Development in the Mouse.” Endocrine Reviews 28 (6): 685–705.

Kesavan, Gokul, Fredrik Wolfhagen Sand, Thomas Uwe Greiner, Jenny Kristina Johansson, Sune Kobberup, Xunwei Wu, Cord Brakebusch, and Henrik Semb. 2009. “Cdc42-Mediated Tubulogenesis Controls Cell Specification.” Cell 139 (4): 791–801.

Kopinke, Daniel, and L. Charles Murtaugh. 2010. “Exocrine-to-Endocrine Differentiation Is Detectable Only prior to Birth in the Uninjured Mouse Pancreas.” BMC Developmental Biology 10 (April): 38.

Lacunza, E., V. Ferretti, C. Barbeito, A. Segal-Eiras, and M. V. Croce. 2010. “Immunohistochemical Evidence of Muc1 Expression during Rat Embryonic Development.” European Journal of Histochemistry: EJH 54 (4): e49.

Larsen, Hjalte List, Laura Martín-Coll, Alexander Valentin Nielsen, Christopher V. E. Wright, Ala Trusina, Yung Hae Kim, and Anne Grapin-Botton. 2017. “Stochastic Priming and Spatial Cues Orchestrate Heterogeneous Clonal Contribution to Mouse Pancreas Organogenesis.” Nature Communications 8 (1): 605.

Lawrence, Michael, Wolfgang Huber, Hervé Pagès, Patrick Aboyoun, Marc Carlson, Robert Gentleman, Martin T. Morgan, and Vincent J. Carey. 2013. “Software for Computing and Annotating Genomic Ranges.” PLoS Computational Biology 9 (8): e1003118.

Löf-Öhlin, Zarah M., Pia Nyeng, Matthew E. Bechard, Katja Hess, Eric Bankaitis, Thomas U. Greiner, Jacqueline Ameri, Christopher V. Wright, and Henrik Semb. 2017. “EGFR Signalling Controls Cellular Fate and Pancreatic Organogenesis by Regulating Apicobasal Polarity.” Nature Cell Biology 19 (11): 1313–25.

Luan, Zhou, Yoshihiro Morimoto, Atsushi Fushimi, Nami Yamashita, Wenhao Suo, Atrayee Bhattacharya, Masayuki Hagiwara, Caining Jin, and Donald Kufe. 2022. “MUC1-C Dictates Neuroendocrine Lineage Specification in Pancreatic Ductal Adenocarcinomas.” Carcinogenesis 43 (1): 67–76.

Lubarsky, Barry, and Mark A. Krasnow. 2003. “Tube Morphogenesis: Making and Shaping Biological Tubes.” Cell 112 (1): 19–28.

Mamidi, Anant, Christy Prawiro, Philip A. Seymour, Kristian Honnens de Lichtenberg, Abigail Jackson, Palle Serup, and Henrik Semb. 2018. “Mechanosignalling via Integrins Directs Fate Decisions of Pancreatic Progenitors.” Nature 564 (7734): 114–18.

Mangan, Anthony J., Daniel V. Sietsema, Dongying Li, Jeffrey K. Moore, Sandra Citi, and Rytis Prekeris. 2016. “Cingulin and Actin Mediate Midbody-Dependent Apical Lumen Formation during Polarization of Epithelial Cells.” Nature Communications 7 (August): 12426.

Martín-Belmonte, Fernando, Wei Yu, Alejo E. Rodríguez-Fraticelli, Andrew J. Ewald, Zena Werb, Miguel Alonso, and Keith Mostov. 2008. “Cell-Polarity Dynamics Controls the Mechanism of Lumen Formation in Epithelial Morphogenesis.” Current Biology: CB 18 (7): 507–13.

Marty-Santos, Leilani, and Ondine Cleaver. 2016. “Pdx1 Regulates Pancreas Tubulogenesis and E-Cadherin Expression.” Development 143 (1): 101–12.

Mederacke, Malte, Lisa Conrad, Roman Vetter, and Dagmar Iber. 2022. “Geometric Effects Position Renal Vesicles During Kidney Development.” bioRxiv. 10.1101/2022.08.30.505859.

Metzger, Ross J., Ophir D. Klein, Gail R. Martin, and Mark A. Krasnow. 2008. “The Branching Programme of Mouse Lung Development.” Nature 453 (7196): 745–50.

Mullapudi, Sri Teja, Giulia L. M. Boezio, Andrea Rossi, Michele Marass, Ryota L. Matsuoka, Hiroki Matsuda, Christian S. M. Helker, Yu Hsuan Carol Yang, and Didier Y. R. Stainier. 2019. “Disruption of the Pancreatic Vasculature in Zebrafish Affects Islet Architecture and Function.” Development 146 (21). 10.1242/dev.173674.

Nyeng, Pia, Silja Heilmann, Zarah M. Löf-Öhlin, Nina Fransén Pettersson, Florian Malte Hermann, Albert Reynolds, and Henrik Semb. 2019. “p120ctn-Mediated Organ Patterning Precedes and Determines Pancreatic Progenitor Fate.” Developmental Cell 49 (1): 31–47.e9.

Packard, Adam, Kylie Georgas, Odyssé Michos, Paul Riccio, Cristina Cebrian, Alexander N. Combes, Adler Ju, et al. 2013. “Luminal Mitosis Drives Epithelial Cell Dispersal within the Branching Ureteric Bud.” Developmental Cell 27 (3): 319–30.

Pan, Xinchao, Ulrike Schnell, Courtney M. Karner, Erin V. Small, and Thomas J. Carroll. 2015. “A Cre-Inducible Fluorescent Reporter for Observing Apical Membrane Dynamics.” Genesis 53 (3-4): 285–93.

Percival, A. C., and J. M. Slack. 1999. “Analysis of Pancreatic Development Using a Cell Lineage Label.” Experimental Cell Research 247 (1): 123–32.

Pierreux, Christophe E. 2021. “Shaping the Thyroid: From Peninsula to de Novo Lumen Formation.” Molecular and Cellular Endocrinology 531 (July): 111313.

Puri, Sapna, and Matthias Hebrok. 2007. “Dynamics of Embryonic Pancreas Development Using Real-Time Imaging.” Developmental Biology 306 (1): 82–93.

Reichert, Maximilian, and Anil K. Rustgi. 2011. “Pancreatic Ductal Cells in Development, Regeneration, and Neoplasia.” The Journal of Clinical Investigation 121 (12): 4572–78.

Riccio, Paul, Cristina Cebrian, Hui Zong, Simon Hippenmeyer, and Frank Costantini. 2016. “Ret and Etv4 Promote Directed Movements of Progenitor Cells during Renal Branching Morphogenesis.” PLoS Biology 14 (2): e1002382.

Rueden, Curtis T., Johannes Schindelin, Mark C. Hiner, Barry E. DeZonia, Alison E. Walter, Ellen T. Arena, and Kevin W. Eliceiri. 2017. “ImageJ2: ImageJ for the next Generation of Scientific Image Data.” BMC Bioinformatics 18 (1): 529.

Sakurai, J., N. Hattori, M. Nakajima, T. Moriya, T. Suzuki, A. Yokoyama, and N. Kohno. 2007. “Differential Expression of the Glycosylated Forms of MUC1 during Lung Development.” European Journal of Histochemistry: EJH 51 (2): 95–102.

Scheele, Colinda L. G. J., Edouard Hannezo, Mauro J. Muraro, Anoek Zomer, Nathalia S. M. Langedijk, Alexander van Oudenaarden, Benjamin D. Simons, and Jacco van Rheenen. 2017. “Identity and Dynamics of Mammary Stem Cells during Branching Morphogenesis.” Nature 542 (7641): 313–17.

Shaner, Nathan C., Robert E. Campbell, Paul A. Steinbach, Ben N. G. Giepmans, Amy E. Palmer, and Roger Y. Tsien. 2004. “Improved Monomeric Red, Orange and Yellow Fluorescent Proteins Derived from Discosoma Sp. Red Fluorescent Protein.” Nature Biotechnology 22 (12): 1567–72.

Shih, Hung Ping, Devin Panlasigui, Vincenzo Cirulli, and Maike Sander. 2016. “ECM Signaling Regulates Collective Cellular Dynamics to Control Pancreas Branching Morphogenesis.” Cell Reports 14 (2): 169–79.

Shu, Xiaokun, Nathan C. Shaner, Corinne A. Yarbrough, Roger Y. Tsien, and S. James Remington. 2006. “Novel Chromophores and Buried Charges Control Color in mFruits.” Biochemistry 45 (32): 9639–47.

Sievert, Carson. 2020. Interactive Web-Based Data Visualization with R, Plotly, and Shiny. CRC Press.

Sugimura, Kaoru, Pierre-François Lenne, and François Graner. 2016. “Measuring Forces and Stresses in Situ in Living Tissues.” Development 143 (2): 186–96.

Sznurkowska, Magdalena K., Edouard Hannezo, Roberta Azzarelli, Steffen Rulands, Sonia Nestorowa, Christopher J. Hindley, Jennifer Nichols, et al. 2018. “Defining Lineage Potential and Fate Behavior of Precursors during Pancreas Development.” Developmental Cell 46 (3): 360–75.e5.

The R core Team. 2021. R: A Language and Environment for Statistical Computing. https://www.R-project.org/.

Türkvatan, Aysel, Ayşe Erden, Mehmet Akif Türkoğlu, and Özlem Yener. 2013. “Congenital Variants and Anomalies of the Pancreas and Pancreatic Duct: Imaging by Magnetic Resonance Cholangiopancreaticography and Multidetector Computed Tomography.” Korean Journal of Radiology: Official Journal of the Korean Radiological Society 14 (6): 905–13.

Varner, Victor D., and Celeste M. Nelson. 2014. “Cellular and Physical Mechanisms of Branching Morphogenesis.” Development 141 (14): 2750–59.

Villasenor, Alethia, Diana C. Chong, Mark Henkemeyer, and Ondine Cleaver. 2010. “Epithelial Dynamics of Pancreatic Branching Morphogenesis.” Development 137 (24): 4295–4305.

Warming, Søren, Nina Costantino, Donald L. Court, Nancy A. Jenkins, and Neal G. Copeland. 2005. “Simple and Highly Efficient BAC Recombineering Using galK Selection.” Nucleic Acids Research 33 (4): e36.

Wesseling, J., S. W. van der Valk, and J. Hilkens. 1996. “A Mechanism for Inhibition of E-Cadherin-Mediated Cell-Cell Adhesion by the Membrane-Associated Mucin episialin/MUC1.” Molecular Biology of the Cell 7 (4): 565–77.

Wickham, Hadley, Mara Averick, Jennifer Bryan, Winston Chang, Lucy McGowan, Romain François, Garrett Grolemund, et al. 2019. “Welcome to the Tidyverse.” Journal of Open Source Software 4 (43): 1686.

Zhou, Qiao, Anica C. Law, Jayaraj Rajagopal, William J. Anderson, Paul A. Gray, and Douglas A. Melton. 2007. “A Multipotent Progenitor Domain Guides Pancreatic Organogenesis.” Developmental Cell 13 (1): 103–14.

